# Direct Assessment of Nitrative Stress in Lipid Environments: Applications of a Designer Lipid-Based Biosensor for Peroxynitrite

**DOI:** 10.1101/2022.06.22.497268

**Authors:** Bryan Gutierrez, Tushar Aggarwal, Huseyin Erguven, M. Rhia L. Stone, Changjiang Guo, Alyssa Bellomo, Elena Abramova, Emily R. Stevenson, Andrew J. Gow, Enver Cagri Izgu

## Abstract

Lipid environments can be chemically impacted by peroxynitrite (ONOO^−^), a reactive species generated under nitrative stress. Molecular tools used for investigating ONOO^−^ reactivity in biological membranes remain underdeveloped, available probes lack the ability of subcellular localization, and the standard methods for detecting ONOO^−^ *in vivo* are indirect. Here we investigated ONOO^−^ in diverse lipid environments (biomimetic giant vesicles, live mammalian cells, and within the lung lining) using a biocompatible and membrane-localized phospholipid named **DPPC-TC-ONOO^−^**. This designer lipid and 1-palmitoyl-2-oleoyl-sn-glycero-3-phosphocholine self-assemble to giant vesicles that respond to ONOO^−^ by generating fluorescence. These vesicles remain intact after sensing ONOO^−^ and exhibit excellent selectivity against other redox species. We delivered **DPPC-TC-ONOO^−^** into live HeLa and RAW cells via lipid nanoparticles (LNPs). Cytokine-induced nitrative stress led to enhanced fluorescence of the lipid clusters, primarily in the endoplasmic reticulum. These LNPs allowed the detection of ONOO^−^ reactivity and nitrative stress around bronchioles within precision cut lung slices in response to acute lung injury (ALI). Furthermore, the use of the LNPs allowed for the detection of pulmonary macrophages from bronchoalveolar lavage following ALI in C57BL6/J but not in *Nos2^−/–^* mice. These investigations revealed significant advantages of **DPPC-TC-ONOO^−^** over its non-amphiphilic analog. Our work presents (i) an unprecedented function for biomimetic membranes, (ii) the potential of LNPs for delivering designer lipids into cells and tissues, (iii) real-time imaging of endogenous ONOO^−^ at the organelle level in mammalian cells, and (iv) a direct method of studying nitrative stress due to ALI *ex vivo* and *in vivo*.

## INTRODUCTION

Lipid membranes and lipid-rich environments are potential targets of reactive oxygen and nitrogen species (RONS).^1, 2^ A major source of lipid membrane damage is peroxynitrite (ONOO^−^)^3^ which is produced through the reaction between nitric oxide (^•^NO) and superoxide (O_2_^•–^) at near diffusional rates (∼5 x 10^9^ mol^-1^s^-1^).^4, 5^ It has been proposed that biological membranes and hydrophobic tissue compartments are critical locations for formation of ^•^NO-derived reactive species.^6^ The majority of ONOO^−^ formation is likely to occur in proximity to the lipid portion of the cell, as ^•^NO is over 1000 times more soluble in hydrophobic environments.^5^ Molecular tools for investigating the presence and reactivity of ONOO^−^ in lipid membranes and lipid-rich organelles or tissues will provide significant insights into pathophysiologies associated with nitrative stress.^3, 7, 8^

The mechanism of peroxynitrite-mediated lipid damage has been proposed to involve hydrogen-abstraction from unsaturated lipids, such as linolenic acid and arachidonic acid, generating carbon-centered lipid radicals.^1, 9^ These radicals react with nearby unsaturated lipids and are trapped by molecular oxygen (O_2_), forming lipid peroxides. Peroxynitrite reactivity has been linked to protein aggregation in neurons associated with Parkinson’s disease^10–12^ and other synucleinopathies^13^ while the same reactivities have also been associated with the progression of cardiovascular diseases like diabetic cardiomyopathy.^14^ Inducible nitric oxide synthase (iNOS) catalyzes the production of ^•^NO from L-arginine and O_2_ in macrophages in response to various stimuli, including bacterial or viral infections, inflammation, and proinflammatory cytokines.^15^ In the case of RAW 264.7, a widely used macrophage model, peroxynitrite production can be stimulated by lipopolysaccharide (LPS). LPS binds Toll-like receptor 4 (TLR4), which triggers the activation of nuclear factor kappa B (NF-KB) signaling, resulting in the production of iNOS^16, 17^ and ROS^18, 19^ In HeLa, peroxynitrite production through iNOS is facilitated by the combination of interferon-γ (IFN-γ), LPS, and phorbol myristate acetate (PMA)^20^ The expression and activity of iNOS must be tightly regulated, because excessive or prolonged iNOS activation is detrimental to the host, leading to tissue damage and fibrosis^21^ Macrophage upregulation of iNOS production in response to injury is a characteristic of pro-inflammatory activation. One such model is intratracheal bleomycin (ITB)-induced ALI.^22^ The extent of pulmonary ITB injury has been demonstrated to be heavily iNOS dependent.^23^ This system, therefore, presents a relevant biological model to examine the efficacy of ONOO^−^ detection *in vivo*.

The protonated form of ONOO^−^, peroxynitrous acid (ONOOH, p*K*_a_ = 6.8), has a chemical half-life of 0.63 seconds^3^ The biological half-life of ONOO^−^ and ONOOH depends on the concentration of the biological targets with which they react and has been estimated at ∼10–20 ms.^2^ As a result, investigating the generation and reactivity of ONOO^−^ in biological systems is challenging.^24^ In addition, although there is a significant and evolving interest in developing biosensing tools to study lipid membranes,^25–28^ technologies for investigating peroxynitrite reactivity in biological lipid membranes remain underdeveloped^29, 30^ Furthermore, investigating the reactivity of ONOO^−^ in biological systems has relied on indirect measurements, including immunohistochemistry,^23^ chemiluminesce,^23^ or EPR (electron paramagnetic resonance)^31^ or bite marks, such as tyrosine nitration,^32^ or lipid peroxidation.^33^ These markers are neither specific nor continuous, making cellular visualization of ONOO^−^ generation and reactivity challenging. Taken together, there is a need for direct and membrane-specific peroxynitrite probes that function in living systems without interfering with redox events.

Herein, we report the development, characterization, membrane localization, and multifaceted utilization of **DPPC-TC-ONOO^−^**, a high-fidelity peroxynitrite probe derived from 1,2-dipalmitoyl-*rac*-glycero-3-phosphocholine (*rac*-DPPC). This designer phospholipid senses ONOO^−^ proximal to lipid membranes in both biomimetic systems and mammalian cells with diverse origins. Using **DPPC-TC-ONOO^−^** and 1-palmitoyl-2-oleoyl-*sn*-glycero-3-phosphocholine (POPC), we built models of protocell membranes in the form of giant vesicles (GVs). These GVs responded to ONOO^−^ by lighting up at the membrane. The biophysical integrity of the GVs is retained after sensing ONOO^−^ and they displayed excellent selectivity against other RONS. Lipid nanoparticles (LNPs) obtained from 1,2-di-*O*-octadecenyl-3-trimethylammonium propane (DOTMA) and 1,2-dioleoyl-*sn*-glycero-3-phosphoethanolamine (DOPE) successfully delivered **DPPC-TC-ONOO^−^** to live HeLa and RAW 264.7 cells. Real-time confocal imaging demonstrated substantial fluorescence enhancement of the ER in HeLa under nitrative stress induced through endogenous stimulation with IFN-γ/LPS/PMA. Assays for RAW 264.7 cells used LPS-mediated stimulation, which led to enhanced fluorescence of lipid clusters in both the ER and Golgi. The redox selectivity of **DPPC-TC-ONOO^−^** was confirmed to be iNOS-dependent through control experiments that use an inhibitor of iNOS. The biological potential of **DPPC-TC-ONOO^−^**was further demonstrated through murine models of *ex vivo* and *in vivo* ALI. For the *ex vivo* ALI study, we used mechlorethamine hydrochloride, a carcinogen historically categorized as nitrogen mustard (NM), a well-characterized pulmonary toxicant known to induce nitrative stress^34^ An organotypic culture of precision cut lung slices (PCLS) was used to determine the cellular localization of **DPPC-TC-ONOO^−^** within the tissue. Through LNP-assisted delivery, iNOS-dependent nitrative stress was detected around bronchioles upon treatment of PCLS with NM. For the *in vivo* investigation of ALI, immunostaining of bronchoalveolar lavage (BAL) cells, which are 95% or more macrophages,^35^ indicated that **DPPC-TC-ONOO^−^**responds to iNOS-dependent nitrative stress upon intratracheal bleomycin (ITB) challenge. Control experiments revealed that **TEG-TC-ONOO^−^**, which is the lipid-free, triethylene glycol (TEG) substituted analog of **DPPC-TC-ONOO^−^**, lacked the ability to localize in membranes or generate detectable signal in PCLS under nitrative stress. The modular design, LNP-assisted delivery, and localized function of **DPPC-TC-ONOO^−^** demonstrate the possibility of developing targeted lipid-based molecular tools for sensing redox species in proximity to membranes.

## RESULTS AND DISCUSSIONS

### The peroxynitrite-sensing phospholipid and its non-amphiphilic analog were constructed through a modular chemical design

We developed a concise synthetic route for the peroxynitrite-sensing phospholipid, **DPPC-TC-ONOO^−^** (Figure 1A), which includes *rac*-DPPC as the phospholipid framework, 3-triazole-coumarin as the fluorophore, and *p*-(4,4,4-trifluoro-3-oxobutyl)phenyl as the peroxynitrite-reactive module. To facilitate investigations of the peroxynitrite-mediated florescence activation mechanism and fidelity, we synthesized its non-amphiphilic analog, **TEG-TC-ONOO^−^**, which contains TEG in place of *rac*-DPPC (Figure 1). **TEG-TC-ONOO^−^** allows us to make comparisons between the amphiphilic and non-amphiphilic probe designs in terms of localization. **TEG-TC-ONOO^−^** lacks the ability to localize in biological membranes as observed in our *in vitro* and *ex vivo* studies. In addition, **TEG-TC-ONOO^−^** exists in a non-aggregated state in aqueous media, making the spectroscopic and spectrophotometric analyses of its reaction with ONOO^−^ and other redox species straightforward. For both **DPPC-TC-ONOO^−^** and **TEG-TC-ONOO^−^**, the *p*-(4,4,4-trifluoro-3-oxobutyl)phenyl and 3-triazole-7-hydroxycoumarin groups were connected through a carbonate linkage. We reasoned that this connection could mask 7-*O* of the coumarin, diminishing its fluorescence intensity. In the presence of ONOO^−^, the *p*-(4,4,4-trifluoro-3-oxobutyl)phenyl moiety would undergo an oxidative spirocyclization, which is in agreement with the previously reported transformation of a dichlorofluorescein caged at its xantane oxygen as diaryl ether.^36^ We hypothesized that for our probe design the spirocyclization should generate a zwitterionic carbonate that reacts with water, eliminating the oxa-spiro[4,5] decenone ***6***, uncaging the coumarin motif via decarboxylation, subsequently forming **DPPC-TC** or **TEG-TC** (Figure 1B).

**Figure 1.**
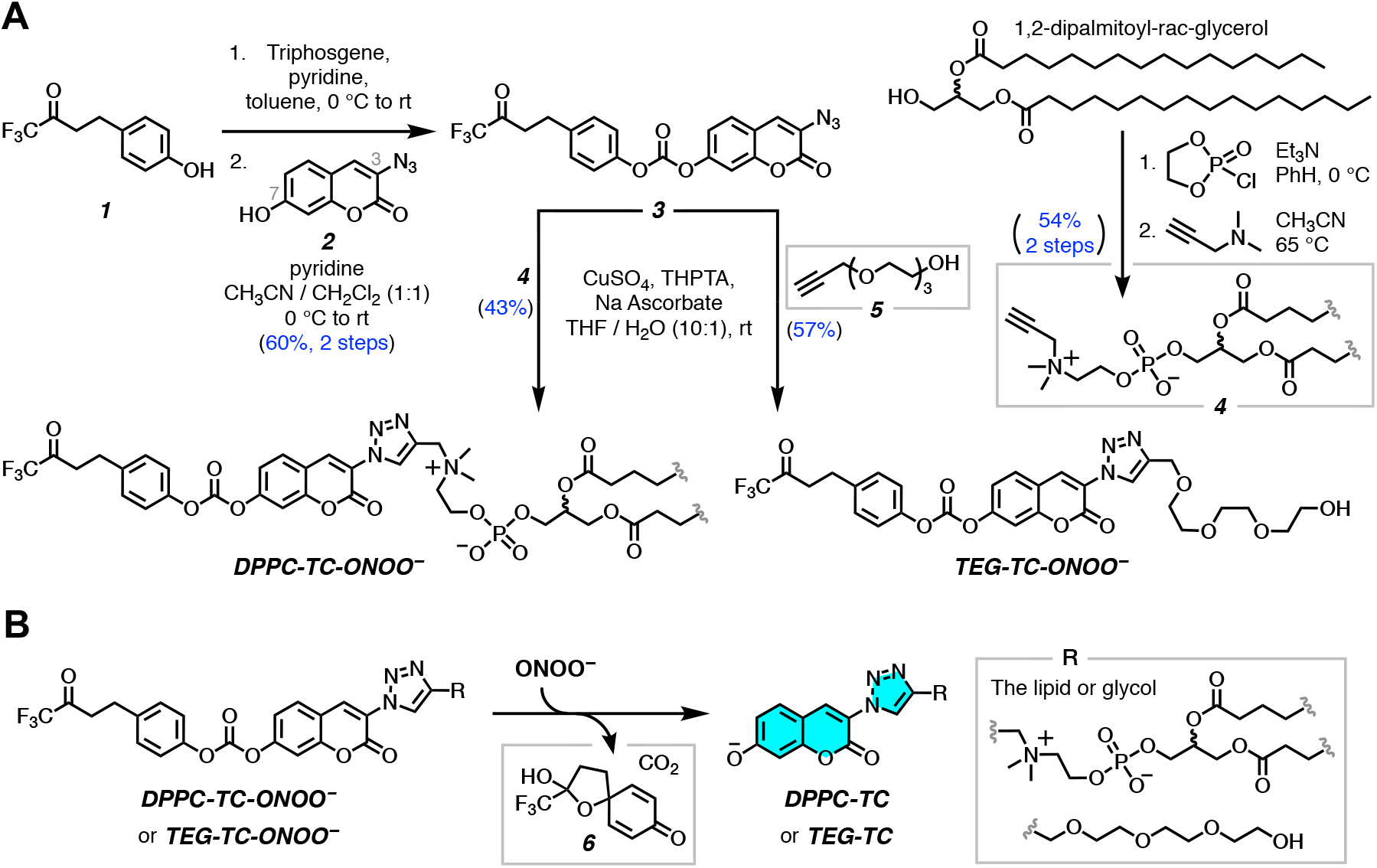
(**A**) The chemical synthesis of peroxynitrite probes **DPPC-TC-ONOO^−^** and **TEG-TC-ONOO^−^**. **(B)** Peroxynitrite-mediated conversion of **TEG-TC-ONOO^−^** and **DPPC-TC-ONOO^−^** to their fluorogenically active products **DPPC-TC** and **TEG-TC**.

Both **DPPC-TC-ONOO^−^** and **TEG-TC-ONOO^−^**were synthesized through convergent routes in which the longest linear sequence required three chemical steps. First, 1,1,1-trifluoro-4-(4-hydroxyphenyl)butan-2-one (***1***) was converted to its chloroformate using triphosgene and successively connected with 3-azido-7-hydroxycoumarin (***2***) to furnish the carbonate **3** in 60% yield over two steps. This compound served as a common intermediate in the construction of **DPPC-TC-ONOO^−^** and **TEG-TC-ONOO^−^**. Through the copper-catalyzed azide alkyne cycloaddition reaction, the carbonate **3** was conjugated with either the propargylated *rac*-DPPC ***4*** or propargyl-TEG-OH ***5*** to afford **DPPC-TC-ONOO^−^**(43%) or **TEG-TC-ONOO^−^** (57%), respectively. To obtain the alkynyl lipid ***4***, we converted 1,2-dipalmitoyl-*rac*-glycerol to its cyclic phosphate triester using ethylene chlorophosphite and opened the subsequent ring by the nucleophilic addition of 3-dimethylamino-1-propyne (see Supporting Information for details).

### Sensing occurs via oxidative uncaging of the coumarin motif and is specific to peroxynitrite

To gain a mechanistic insight into the reaction of the probes with ONOO^−^, we examined the products by liquid chromatography followed by high-resolution mass spectroscopy (LC-HRMS) (Supplemental Figure 1). The amphiphilic probe, **DPPC-TC-ONOO^−^**, displayed poor electrospray ionization profile, while **TEG-TC-ONOO^−^** provided evidence for the formation of its oxidative decarboxylation product, **TEG-TC**, upon mixing with ONOO^−^. This mass spectroscopy data do not provide direct evidence of a reactive carbonate intermediate, and the identification of such is outside the scope of this project. However, we observed the progressive formation of the oxa-spiro[4,5] decenone ***6***, which provides support to a potential spirocyclization/elimination cascade. We then determined the p*K*_a_ of **TEG-TC** to be 7.1 ± 0.1 using UV-Vis spectrophotometric measurements (Supplemental Figure 2). Under our confocal imaging conditions (pH 7.5– 8.5 with cell-free assays, pH 7.4 with cells), a substantial population of the 7-hydroxycoumarin motif should exist in its fluorogenically active aryloxide (deprotonated) form. Indeed, measuring fluorescence (405/475 nm) of **TEG-TC** for different pHs (5.8–8.8) showed that the signal intensities from the samples at pHs 7.5–8.8 were similar and 18–20-fold higher than that at pH 5.8 (Supplemental Figure 3). We assessed the change in triazole-coumarin fluorogenicity upon uncaging the probes by measuring the relative fluorescence quantum yield (Φ_F*rel*_) values (Table 1) (for absorbance and emission spectra, see Supplemental Figure 4). As the standard, we used coumarin 343, whose absolute fluorescence quantum yield (Φ_F_) is 0.63 in ethanol.^37^ The Φ_F*rel*_ values of **TEG-TC-ONOO^−^**(entry 1) and **DPPC-TC-ONOO^−^** (entry 3) were measured as 0.08 and 0.09, respectively. Each with a low Φ_F*rel*_ and molar extinction coefficient, these probes were validated to exhibit weak brightness due to derivatization of their coumarin 7-*O* atom as a carbonate group. The Φ_F*rel*_ values for **TEG-TC** (entry 2) and **DPPC-TC** (entry 4), which were synthesized independently (see Supporting Information for details), were measured as 0.66 and 0.64, respectively. Taken together, these results suggested that conversion of the caged coumarins to their uncaged forms enhances the fluorescence quantum yield by ≥7-fold and brightness by ∼30-to-35 fold.

**Table 1.** Characterization of the peroxynitrite probes and their uncaged forms.

| Entry | Compound | Max Ex (nm) | Max Em (nm) | Extinction Coefficient ($M^{-1} cm^{-1}$) | Quantum Yield | Brightness | Relative Brightness |
| --- | --- | --- | --- | --- | --- | --- | --- |
| 1 | TEG-TC-ONOO<sup>-</sup> | 400.0 | 472.0 | 4475 | 0.08 | 358 | 1.4 |
| 2 | TEG-TC | 397.0 | 471.4 | 15613 | 0.66 | 10305 | 41.1 |
| 3 | DPPC-TC-ONOO<sup>-</sup> | 395.0 | 471.2 | 2784 | 0.09 | 251 | 1.0 |
| 4 | DPPC-TC | 398.0 | 472.0 | 13749 | 0.64 | 8799 | 35.1 |

The fluorogenic response rate and magnitude of the probe were evaluated by a series of fluorescence measurements with **TEG-TC-ONOO^−^**(Figure 2A–C). Physiologically normal ONOO^−^ production rate in cells has been estimated to be in the single-digit µM/s range.^3, 38^ Guided by this information, and in agreement with the fluorescence titration conditions employed for cytosolic resorufin-based peroxynitrite sensors,^39^ we reasoned that administering 600 μM of ONOO^−^ into a pH 7.5 medium would provide an effective level of ONOO^−^ exposure equivalent to a biological flux rate of ∼10–20 μM/s. Existence of ONOO^−^ at pH 7.5 was recorded with both CO_2_-rich and CO_2_-deprived aqueous solutions using UV-Vis spectrophotometry (Supplemental Figure 5). We tested pH dependance of fluorescence generation from **TEG-TC-ONOO^−^** using buffers with varying pHs, including 6.8, which is the p*K*_a_ of ONOOH.^3^ Addition of ONOO^−^ (600 μM) led to a significant increase in fluorescence at pHs 7.8 and 8.8, which continued until 30 minutes, reaching up to 6.5-to-8 times of the untreated samples (t = 0 min) (Figure 2A). This trend of increasing fluorescence output was mirrored in the level of conversion of **TEG-TC-ONOO^−^**into **TEG-TC** at pH 7.5, based on UV-Vis spectrophotometric measurements (Supplemental Figure 6). In contrast, those at lower pHs (5.8 and 6.8) showed little or no fluorescence generation, largely due to the predominant background consumption of ONOO^−^ (e.g., protonation to ONOOH)^3, 24^ or that surviving ONOOH does not lead to activation of the probe. It is also worth noting that for pH (5.8 or 6.8) which is below the p*K*_a_ of **TEG-TC** (7.1), weakens the fluorescence output (cf. Supplemental Figure 3), likely due to the decreased ratio of the deprotonated coumarin over its protonated form. The observation that fluorescence increases over 30 minutes supports a multi-step mechanism for the formation of **TEG-TC** from **TEG-TC-ONOO^−^**. To gain further insight, we conducted a competitive oxidation assay where we measured F/F_0_ with and without the ketone ***1*** (Figure 2B). This ketone generates no fluorescence when treated with ONOO^−^ (600 equiv) (gray square). If the starting mixture contains **TEG-TC-ONOO^−^** (1 equiv) and excess of ***1*** (600 equiv), ONOO^−^ (600 equiv) addition generates only weak fluorescence (orange triangle), indicating that ***1*** can largely scavenge ONOO^−^. When ***1*** (600 equiv) is introduced after 5 minutes of incubation of **TEG-TC-ONOO^−^** (1 equiv) with ONOO^−^ (600 equiv), fluorescence generation is substantial (inverted blue triangle), following a trend similar to that for **TEG-TC-ONOO^−^** treated with ONOO^−^ (red circle). These results suggest that the reaction between the probe and ONOO^−^ might be relatively fast for the tested condition, yet fluorescence generation from the probe is likely a complex, multistep mechanism.

**Figure 2.**
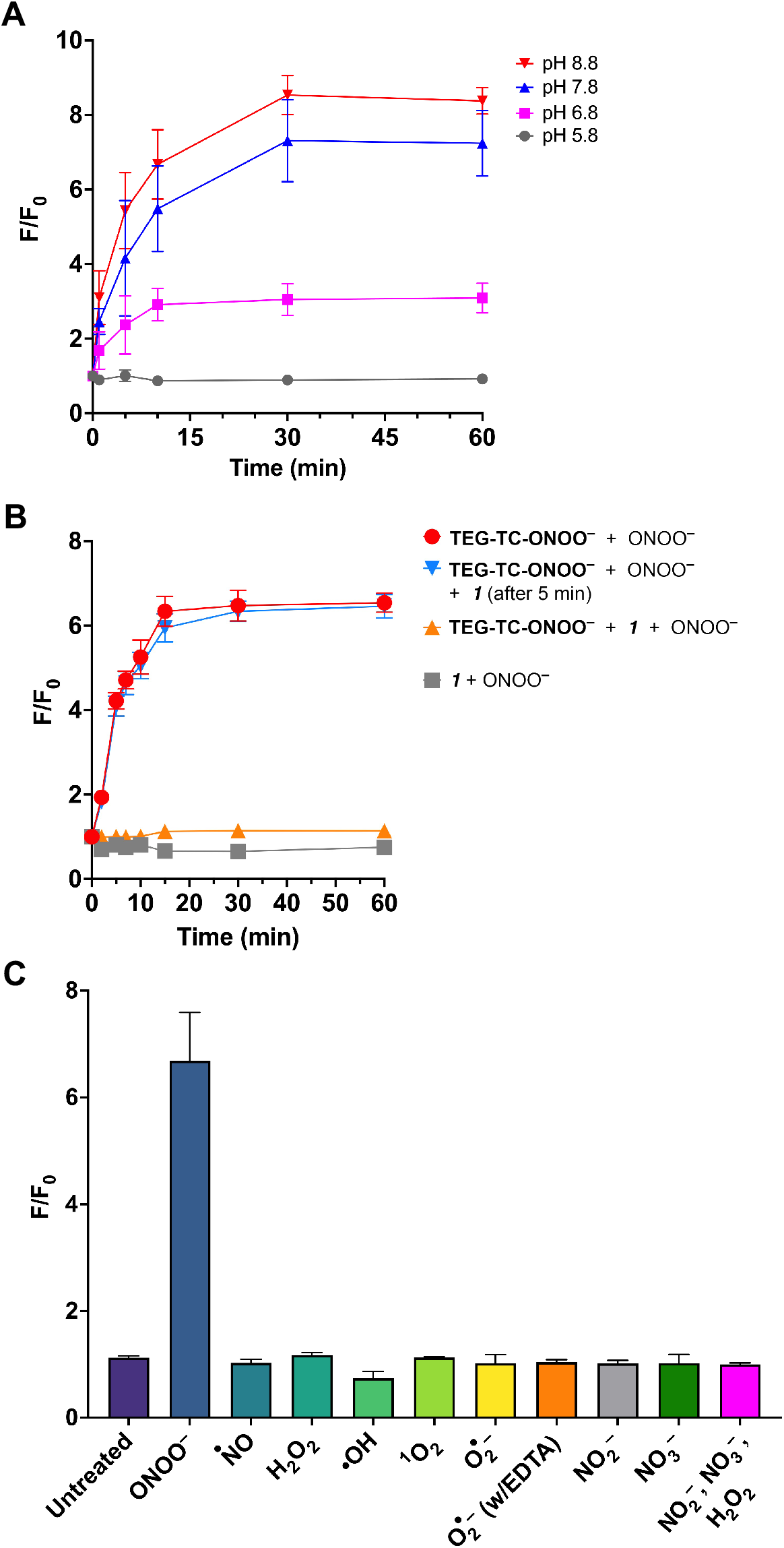
(**A**) The time-dependent change in relative fluorescence intensity (F) with respect to the background fluorescence intensity (F_0_) of **TEG-TC-ONOO^−^** (1 µM) when treated with ONOO^−^ (600 µM) at various pHs: 5.8, 6.8, 7.8, and 8.8. Buffer: 50 mM Tris (pH 6.8, 7.8, or 8.8) or MES (pH 5.8). (**B)** The time-dependent change in F/F_0_ for **TEG-TC-ONOO^−^**and/or 1,1,1-trifluoro-4-(4-hydroxyphenyl)butan-2-one (compound ***1***). **TEG-TC-ONOO^−^**(1 µM), ***1*** (600 µM), and ONOO^−^ (600 µM). The solutions were prepared using Tris 7.5 (50mM) buffer. (**C)** Selectivity of **TEG-TC-ONOO^−^** against RONS. **TEG-TC-ONOO^−^** (1 µM) was dissolved in Tris buffer (50 mM, pH 7.5) and subjected to RONS with an estimated initial concentration of 600 µM. EDTA (600 µM) was added to chelate transition metals. Data represent F/F_0_ measurements of samples incubated for 60 min. **(A–C)** Ex/Em: 405/475 nm. Error bars represent standard deviation, *n* = 3.

The peroxynitrite-specificity of the sensor moiety was assessed by subjecting **TEG-TC-ONOO^−^** to other redox species, including ^•^NO, hydrogen peroxide (H_2_O_2_), singlet oxygen (^1^O_2_), hydroxyl radicals (HO^•^), and O_2_^•–^ (Figure 2C). Change in fluorescence intensity following excitation at 405 nm was measured over 60 minutes of incubation of **TEG-TC-ONOO^−^** with each species and presented based on the signal-to-background ratio *F/F_0_*, where F and F_0_ are defined as the fluorescence with and without the analyte. In the absence of a redox species, F/F_0_ for **TEG-TC-ONOO^−^** increased only 13% (F/F_0_ = 1.13) over 60 minutes. Treatment of **TEG-TC-ONOO^−^** with ONOO^−^ gave rise to ∼570% increase in fluorescence (F/F_0_ = 6.69). Notably, incubating **TEG-TC-ONOO^−^** with species other than ONOO^−^ resulted in only negligible levels of change in fluorescence. Among those, ^•^NO had essentially no effect on fluorescence intensity (F/F_0_ = 1.03). The most significant fluorescence enhancement was observed for H_2_O_2_, with an intensity 16% stronger than that of the background (F/F_0_ = 1.16). These results indicate that the relative fluorescence enhancement of **TEG-TC-ONOO^−^** is nearly 36-fold for ONOO^−^ compared to the other RONS tested in this study.

### Giant vesicles respond to peroxynitrite by lighting up at the membrane

Biomimetic membranes can be represented by GVs, which have dimensional properties (1–30 μm in diameter) relevant to those of mammalian cells (10–100 μm), display physical characteristics feasible for confocal fluorescence imaging, and are utilized for bottom-up construction of biomimetic systems.^40, 41^ We studied the membrane localization and chemical reactivity of **DPPC-TC-ONOO^−^** in GVs of mixed unilamellar and multilamellar populations (Figure 3A). We set the pH of GV imaging buffers to 8.5 to extend the lifetime of ONOO^−^, because the sample handling for confocal microscopy requires a longer timescale than that for plate reader fluorescence measurement.

**Figure 3.**
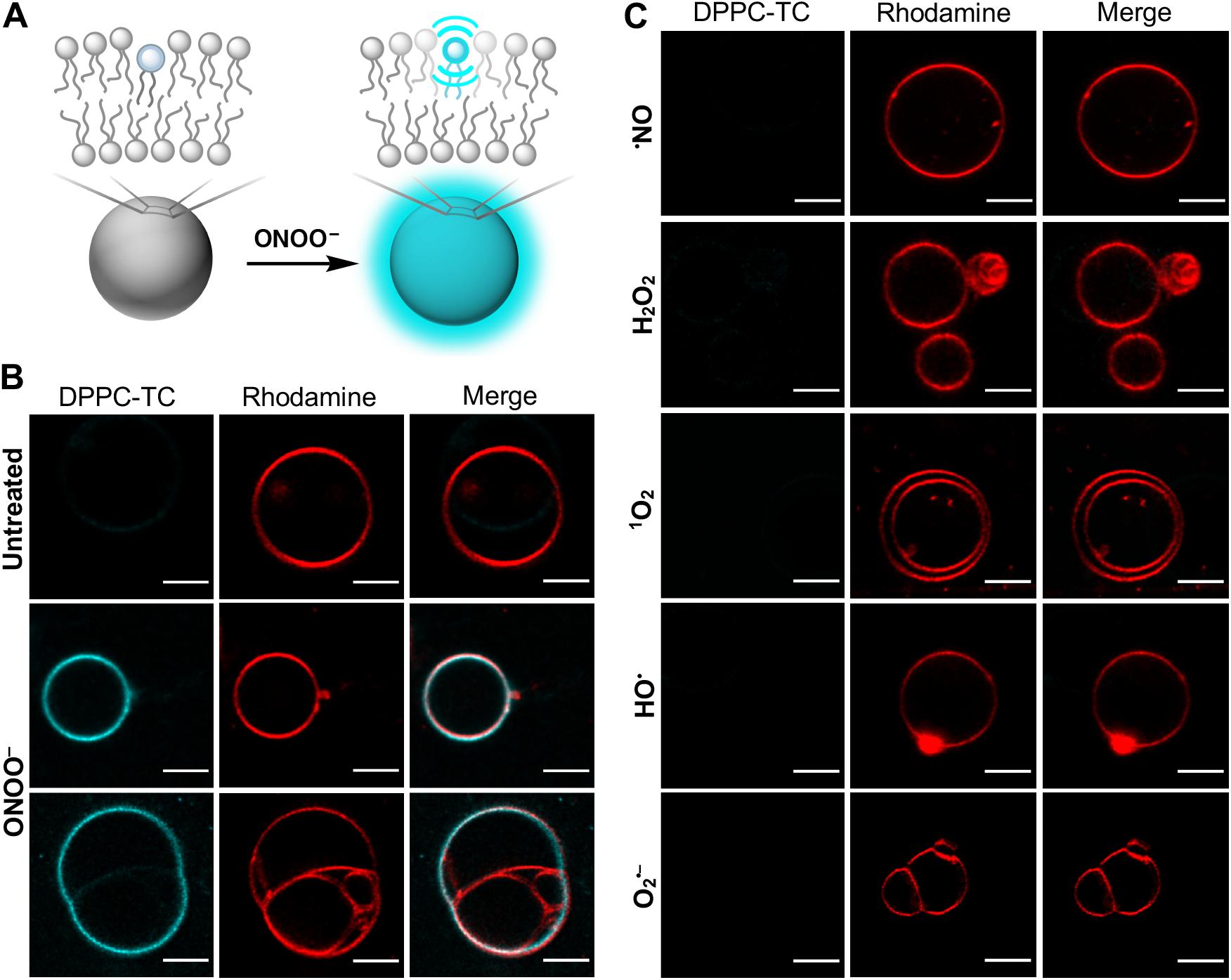
(**A**) Conceptual illustration of vesicle membrane sensing ONOO^−^. **(B)** Confocal images acquired for untreated (top row) and peroxynitrite-treated (middle and bottom rows) GVs. **(C)** Confocal images acquired for GVs treated with other RONS. **(B-C)** Vesicles were prepared from POPC (98.5%, molar ratio) and **DPPC-TC-ONOO^−^** (1%) via electroformation. Liss-Rhod PE (red, 560 nm excitation) was used at 0.5 mol% to label vesicle membranes. In the absence of ONOO^−^ or in the presence of other RONS (100 µM), vesicles emit minimal fluorescence following excitation at 405 nm (DPPC-TC channel). GVs emit strong fluorescence only when ONOO^−^ is present, not when other RONS are used. Fluorescence images are shown for each field of view to demonstrate the localization of the fluorescent phospholipid product **DPPC-TC** with respect to the vesicle membranes. Scale bar = 5 µm.

We produced GVs from POPC and **DPPC-TC-ONOO^−^** (∼99:1 molar ratio) using electroformation^42^ (Supplemental Figure 7). To label vesicle membranes, we employed 1,2-dioleoyl-*sn*-glycero-3-phosphoethanolamine-*N*-(lissamine rhodamine B sulfonyl) (18:1 Liss-Rhod PE) in 0.05 mol percent of the total lipid composition. Sucrose solution (400 mM) was used to hydrate the lipid films. A high solute concentration of sucrose facilitated the formation and stabilization of GV. Following electroformation, solutions of GVs were used within 24 hours. Vesicle size distribution was analyzed by dynamic light scattering (DLS), confirming the presence of GVs. DLS analysis showed that 91% of vesicles had diameters ranging from ∼5 to 20 µm, with 48% having an average size of 5 µm in diameter (Supplemental Figure 8). GVs were then imaged using fluorescence confocal microscopy. The vesicles were treated with the redox species, then mounted directly onto a microscope slide. Although significant vesicle motility was observed, the population density was sufficiently high to locate some static vesicles. Addition of ONOO^−^ (estimated initial concentration of 100 µM) into GVs led to an observable fluorescence (405 nm excitation, “DPPC-TC channel”) throughout the vesicle membrane after 5 minutes of incubation (Figure 3B, middle row). A potential conclusion one can draw from this observation is that **DPPC-TC-ONOO^−^**and **DPPC-TC** lipid product stay in bilayer membranes, with a near homogenous lateral distribution.

Interestingly, confocal imaging of a selected multilamellar vesicle system (Figure 3B, bottom row) showed partial fluorescence light-up following excitation at 405 nm upon administration of ONOO^−^. While the outermost vesicle (either a single or two vesicles fused) interacts with ONOO^−^, the entrapped smaller vesicles exhibited no observable DPPC-TC fluorescence. This observation can be reasoned by the possibilities that (a) ONOO^−^ may have poor membrane permeability in our imaging buffer conditions and/or (b) most of the available ONOO^−^ is consumed by reacting with the probe within the outermost vesicle membrane.

Notably, GVs exhibited selectivity towards ONOO^−^. Incubation (5 minutes) with other redox species resulted in no significant fluorescence generation at 405 nm excitation (Figure 3C). Taken together, these results indicate that (i) **DPPC-TC-ONOO^−^** can localize into the vesicle membranes made of POPC, a natural phospholipid, (ii) it induces no biophysical alteration to the membrane upon reaction with ONOO^−^, and (iii) the detection of ONOO^−^ in the membranes can be achieved with high redox selectivity.

### LNP-assisted delivery of DPPC-TC-ONOO^−^ allows direct imaging of ER targeted by peroxynitrite in live mammalian cells

With the utility of **DPPC-TC-ONOO^−^** proven in synthetic protocell membranes, we sought to incorporate it into live HeLa and RAW 264.7 cells, which are relevant to redox-induced stress and have well-recognized pathways for RONS production. We utilized lipid nanoparticles (LNPs) (∼100 nm in diameter) to deliver **DPPC-TC-ONOO^−^**, which circumvented its limited solubility due to the relatively hydrophobic butanone-coumarin motif. LNPs were obtained via sonication from DOTMA, DOPE, and **DPPC-TC-ONOO^−^** (47.5: 47.5: 5.0 molar ratio), which were hydrated with a 300 mM sucrose solution. DOTMA and DOPE were chosen as the carrier lipids due to their cationic/zwitterionic natures, which has been postulated to enhance cellular uptake through the cell membrane.^43^

The impact of different LNP compositions and incubation times on the cellular incorporation of **DPPC-TC-ONOO^−^** was assessed. Initial trials indicated that the molar concentration of **DPPC-TC-ONOO^−^**in LNPs influenced both the homogeneity of the LNPs and the fluorescence signal generation within stimulated cells. Specifically, LNPs containing greater than 10 mol% **DPPC-TC-ONOO^−^** were considerably more heterogenous in size (up to 20 μm observed, in contrast to ∼100 nm) and nonuniform in vesicle lamellarity. Those with less than 4 mol% **DPPC-TC-ONOO^−^**led to insufficient fluorescence signal. To balance these counteracting factors, 5 mol% **DPPC-TC-ONOO^−^** was used, with 47.5 mol% DOTMA and DOPE, each. In these initial assays, cells were incubated with LNPs for up to 15 hours to complete the uptake and intracellular localization of **DPPC-TC-ONOO^−^**. Shorter incubation times (3-to-5 hours) led to its uptake but not subcellular localization. To facilitate extended incubation timeframes, LNPs were prepared in DMEM (Dulbecco’s Modified Eagle Medium) instead of HBSS (Hanks’ Balanced Salt Solution), which we observed to be deleterious to the cells for incubations exceeding 3 hours. However, high serum concentration in DMEM caused interference between LNPs and serum-protein, resulting in aggregate formation as observed by microscopy. To overcome this obstacle, we used LNPs (50 µM) in a mixture of 2/3 volumetric ratio of sucrose solution and Opti-MEM™, a reduced serum medium known to increase the efficiency of lipofection.^44^

With a feasible LNP preparation and delivery protocol established for live cell imaging, we then turned our attention to assessing the cytocompatibility of the LNPs, with or without **DPPC-TC-ONOO^−^**, **DPPC-TC**, and the oxa-spiro[4,5] decenone ***6***. We employed a standard 3-(4,5-dimethylthiazol-2-yl)-2,5-diphenyl tetrazolium bromide (MTT) cell viability assay in HeLa and RAW 264.7 cells treated with these molecules (8–250 µM) over a 24-hour period. The dose-response curves (Supplemental Figure 9) provided high IC_50_ values for each compound: **DPPC-TC-ONOO^−^** (≥100 µM), **DPPC-TC** (≥50 µM), and 6 (≥70 µM) (Supplemental Table 1). These results suggested that the probe-containing LNPs (50 µM) and the byproducts of the reaction from peroxynitrite (**DPPC-TC** and **6**) do not induce cytotoxicity under our cell imaging conditions.

The biological utility of **DPPC-TC-ONOO^−^** was first demonstrated by imaging live HeLa cells that are under oxidative and nitrosative stress (Figure 4 and Supplemental Figure 10). For a physiologically relevant stress model, we employed the standard stimulation condition involving IFN-γ/LPS/PMA.^19^ Treatment of cells with IFN-γ/LPS leads to the endogenous production of nitric oxide and PMA provokes increased levels of superoxide. These two reactive species combine to form ONOO^−^, as described above. HeLa cells treated with LNPs containing **DPPC-TC** (positive control) exhibited strong fluorescence at 405/475 nm (Supplemental Figure 10A). In contrast, those treated with LNPs containing **DPPC-TC-ONOO^−^** and left unstimulated (negative control) generated no observable signal (Supplemental Figure 10B). Upon endogenous ONOO^−^ generation through stimulation with IFN-γ/LPS/PMA, we observed a substantial increase in fluorescence at 405/475 nm (Figures 4A). Notably, upon PMA addition post IFN-γ/LPS stimulation, the intracellular fluorescence was observable at the 3-minute time point, the earliest possible image acquisition time after cell treatment in our imaging setup. This indicated that, in the background of stimulated iNOS function, PMA instantly leads to ONOO^−^ production (presumably by activating NADPH oxidase directly) and that **DPPC-TC-ONOO^−^**can rapidly sense proximal ONOO^−^. The change in signal intensity after 30 minutes was negligible, suggesting the possibility of probe consumption and/or lack of proximal peroxynitrite. We also evaluated the intracellular redox selectivity of **DPPC-TC-ONOO^−^** using *N*-(3-(aminomethyl)benzyl)-acetamidine, known as 1400W, which inhibits the production of ^•^NO by irreversibly binding to iNOS.^45^ The cells treated with 1400W only (Figure 4B) or treated with 1400W and IFN-γ/LPS/PMA (Figure 4C) generated no observable signal at 405/475 nm. These controls showed that **DPPC-TC-ONOO^−^**selectively senses peroxynitrite generated from iNOS in response to nitrosative but not oxidative stress alone. To determine whether **DPPC-TC-ONOO^−^**localizes and functions at a specific subcellular compartment in HeLa, we carried out a comprehensive colocalization study with dyes specific for lipid-rich and membrane-enclosed organelles, including the ER, mitochondrion, Golgi apparatus, and lysosome (Figures 4D-E). In addition, we used actin stain to gain a visual insight into the cytoskeletal structure of the whole cell. To assess the degree of colocalization, we determined the Pearson’s correlation coefficients (PCC) of the signal from **DPPC-TC** overlapping with that from the selected organelle tracker dye. The **DPPC-TC** signal overlapped almost perfectly (PCC = 0.938) with the perinuclear signal of ER-Tracker™ Green, suggesting a high statistical correlation and preference of **DPPC-TC-ONOO^−^** to localize in the ER. Cells stained with dyes specific for Golgi, mitochondria, and lysosomes displayed less significant correlation.

**Figure 4.**
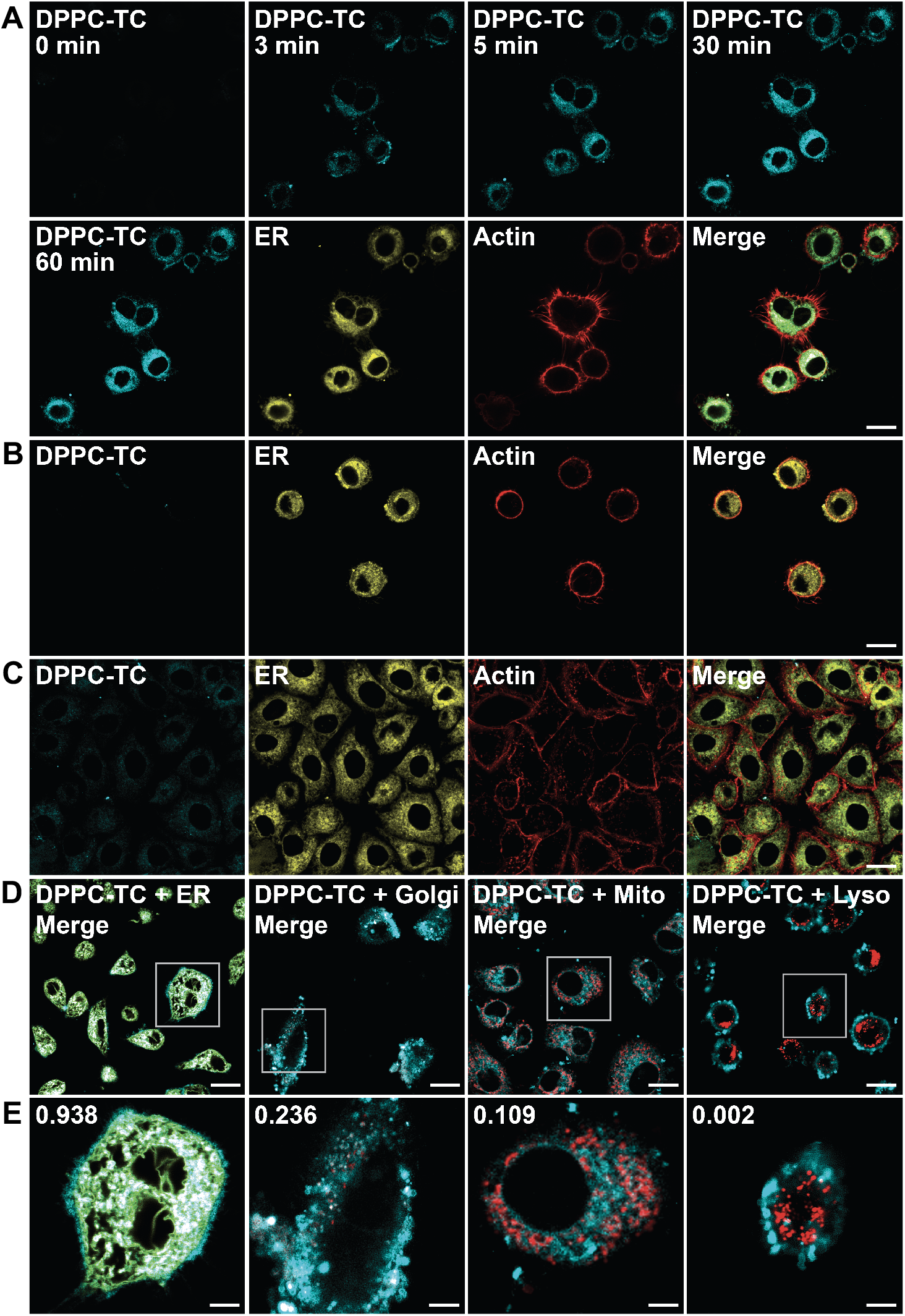
Confocal imaging of lipid environments targeted by ONOO^−^ in live HeLa cells. Cells were incubated with LNPs (50 μM) and stained with organelle trackers and/or actin dye (CellMask™ Deep Red Actin Tracking Stain) prior to imaging. LNPs obtained from a 47.5: 47.5:5.0 molar ratio of DOTMA, DOPE, and **DPPC-TC-ONOO^−^**. **(A)** Cells treated with LNPs, stimulated with IFN-γ/LPS/PMA. Time coarse images acquired upon PMA treatment. **(B)** Cells treated with LNPs and 1400W. **(C)** Cells treated with LNPs, then with 1400W and IFN-γ/LPS/PMA. **(D–E)** Quantitative colocalization study of cells treated with LNPs and stimulated with IFN-γ/LPS/PMA. Organelle trackers: ER-Tracker™, MitoTracker™, CellLight™ Golgi-RFP, LysoTracker™. DPPC-TC channel: 405/475 nm. Scale bars: **(A–D)** 20 µm, **(E)** 5 µm.

We expanded our investigations to include murine-derived macrophage model, RAW 264.7 (Figure 5 and Supplemental Figure 11). RNS play important roles in macrophage activation and differentiation as part of the inflammatory response.^46^ Treatment of RAW 264.7 cells with LPS leads to the expression of iNOS, which produces ^•^NO, as well as increasing O_2_^•–^ formation from multiple sources, generating ONOO^−^. RAW 264.7 cells were treated with LNPs containing either **DPPC-TC** (positive control, Supplemental Figure 11A) or **DPPC-TC-ONOO^−^** (Figure 5 and Supplemental Figure 11B–F). Those treated with LNPs containing **DPPC-TC-ONOO^−^** and left unstimulated generated negligible levels of signal at 405/475 nm (Supplemental Figure 11B). Endogenous ONOO^−^ production through stimulation with LPS led to a substantial increase in fluorescence intensity over 16 hours (Figure 5A). The intracellular redox selectivity of **DPPC-TC-ONOO^−^** was evaluated using 1400W. The cells treated with 1400W only (Figure 5B) or treated with 1400W and LPS (Figure 5C) generated little-to-no signal in the DPPC-TC channel. Assessment of **DPPC-TC-ONOO^−^** localization in RAW 264.7 cells was carried out using the ER, Golgi, mitochondrion, and lysosome trackers (Figures 5D–E). The PCC values for the merged images of ER (0.946) and Golgi (0.729) suggested that **DPPC-TC-ONOO^−^** localizes primarily in the ER and then Golgi. This observation suggests that the probe localization in the ER is predominant but not exclusive, likely due to the abundance of ER-Golgi intermediate compartment (ERGIC) in RAW 264.7 cells and increased prevalence in cross-talk between the ER and Golgi.^47^ For both cell types, the use LNPs appears to be critical for the probe localization. Our control experiments with LNPs containing Liss-Rhod PE showed high statistical correlations (PCC of 0.76–0.94) with the ER tracker (Supplemental Figure 12A-B). Additionally, stimulated cells showed overlapping signals of Liss-Rhod PE and **DPPC-TC** (Supplemental Figure 12C-D). In contrast to the observations with **DPPC-TC-ONOO^−^**, its non-amphiphilic counterpart, **TEG-TC-ONOO^−^**, displayed inferior specificity in regard to subcellular localization (Supplemental Figure 13). It is worth noting that the use of LNPs proved unsuccessful for cellular uptake of either **TEG-TC** or **TEG-TC-ONOO^−^**. Therefore, cells were treated with **TEG-TC** (positive control, Supplemental Figures 13A,B and 13E,F) or **TEG-TC-ONOO^−^**(Supplemental Figures 13C,D and 13G,H) directly and incubated for 20 minutes. Those treated with **TEG-TC-ONOO^−^** were then stimulated via the methods described above. Both the positive controls and stimulated cells displayed **DPPC-TC** signal throughout the cells, including the nucleus. In the case of RAW 264.7 cells, the intracellular homogeneity of the signals were especially substantial (Supplemental Figure 13E,G).

**Figure 5.**
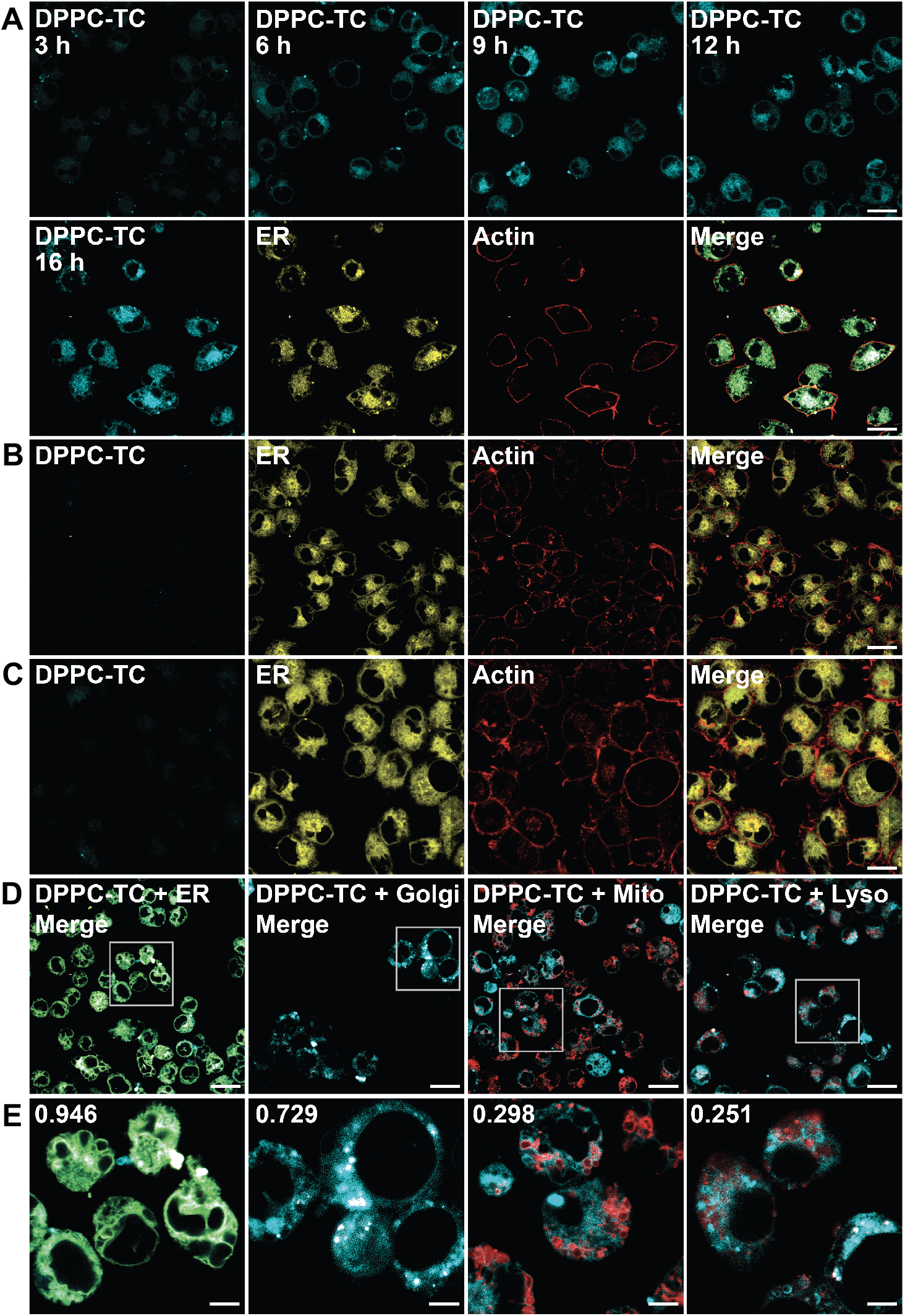
Confocal imaging of lipid environments targeted by ONOO^−^ in live RAW246.7 cells. Cells were seeded into glass microscope dishes, incubated with LNPs (50 μM) and stained with organelle trackers and/or actin dye prior to imaging. LNPs obtained from a 47.5: 47.5:5.0 molar ratio of DOTMA, DOPE, and **DPPC-TC-ONOO^−^**. **(A)** Cells treated with LNPs, stimulated with LPS. Time coarse images acquired after addition of LPS. **(B)** Cells treated with LNPs and 1400W. **(C)** Cells treated with LNPs, then with 1400W and LPS. **(D–E)** Quantitative colocalization study of cells treated with LNPs and stimulated with LPS. Scale bars = **(A–D)** 20 µm or **(E)** 5 µm.

### LNPs carrying DPPC-TC-ONOO^−^ allow for detecting nitrative stress in lung tissue murine models of *ex vivo* and *in vivo* ALI

After successfully imaging the iNOS-dependent ONOO^−^ activity in both HeLa and RAW 264.7, we sought to explore the potential of **DPPC-TC-ONOO^−^**for detecting ONOO^−^ generated in lung tissue murine models of *ex vivo*^35, 48^ and *in vivo* ALI^35^ (Figure 6). To first find an optimal LNP concentration for PCLS, we performed a dose-response assay with **DPPC-TC** (25, 50, 75, 100, 150 µM) in unexposed PCLS (Supplemental Figure 14), which would serve as the positive control in confocal images of PCLS (Supplemental Figure 15). No cytotoxicity was observed for LNP concentrations of 25–100 µM. Based on this result, subsequent studies were performed with 75 µM LNPs in PCLS exposed to NM (Figure 6A and Supplemental Figure 15). In control PCLS (NM-untreated) from C57BL6/J (WT) mice, LNPs carrying **DPPC-TC-ONOO^−^** led to a weak fluorescence signal (405 nm excitation) around the bronchial epithelium, showing basal nitrative stress (Figure 6A, column 1). In PCLS exposed to NM, the fluorescence signal was qualitatively stronger and was primarily localized to the bronchial epithelium (Figure 6A, column 2). The DPPC-TC signal did not overlap with MitoTracker™. Notably, in PCLS obtained from *Nos2^−/–^* mice, weak fluorescence signals were detected for both the control and NM-treated samples in comparison to PCLS from C57BL6/J mice (Figure 6A, columns 3 and 4). These results indicate that the fluorescence generation from **DPPC-TC-ONOO^−^** in the lung tissue is both NM-dependent and iNOS-dependent, validating that the function of **DPPC-TC-ONOO^−^** is specific to nitrative stress. Strikingly, **DPPC-TC** signal is more uniformly accumulated within PCLS compared to **TEG-TC**. PCLS incubated directly with **TEG-TC** (75 µM) showed fluorescence signals largely diffused throughout the tissue (Supplemental Figure 15, column 2). Use of **TEG-TC-ONOO^−^** (75 µM) generated relatively weak signal within PCLS exposed to NM following excitation at 405 nm (Supplemental Figure 15, column 4), suggesting that the non-amphiphilic nature of the probe may hinder its ability to localize in the tissue.

**Figure 6.**
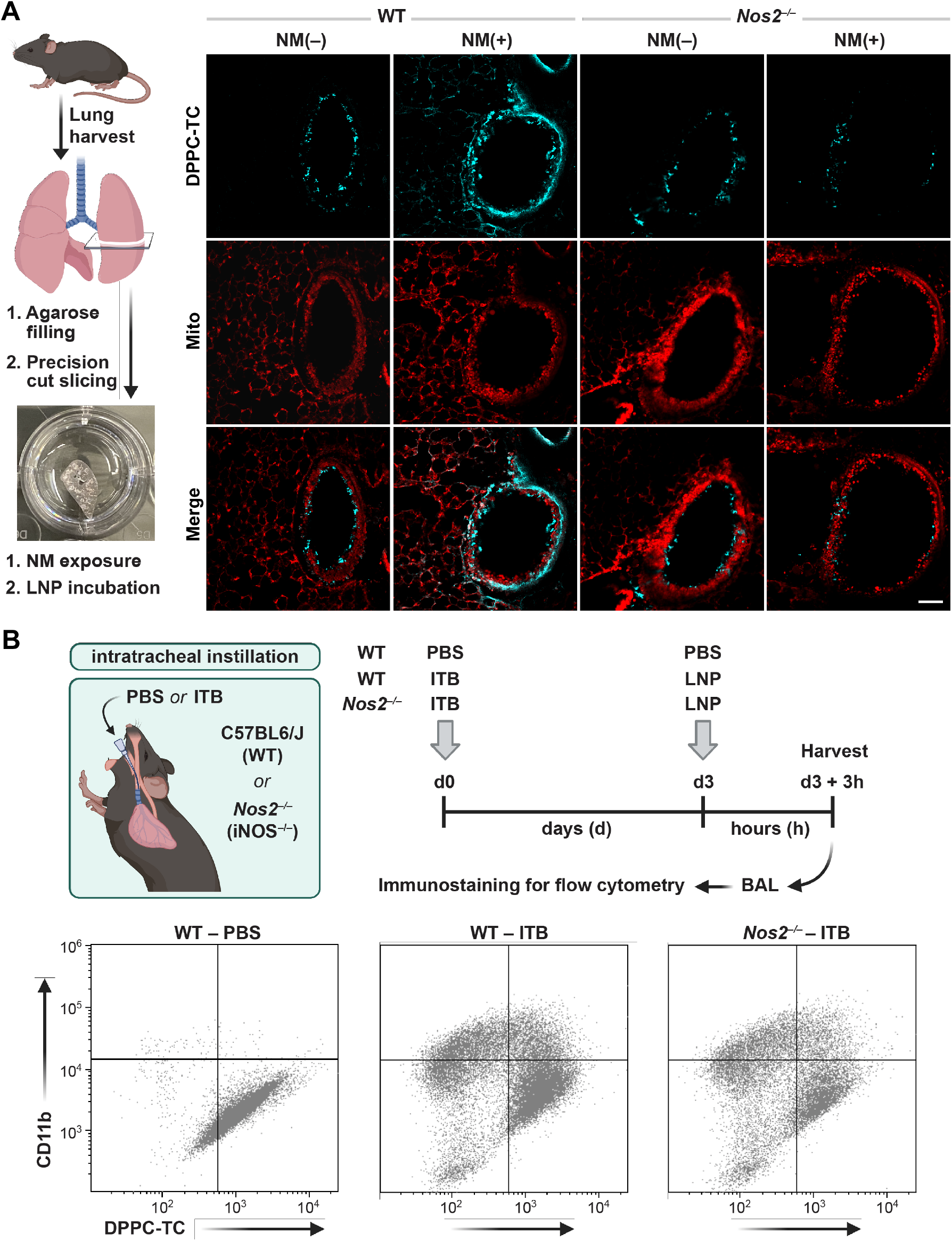
Investigation of the alveolar nitrative stress *ex vivo* and *in vivo.* **(A)** PCLS obtained from C57BL6/J (WT) and *Nos2^−/–^* mice were either untreated (NM(–)) or exposed to nitrogen mustard (NM(+)). After 24 hours, PCLS were incubated with LNPs carrying **DPPC-TC-ONOO^−^** for 1 hour and then imaged. **(B)** ITB-induced ALI and LNP instillation Flow cytometry analysis of myeloid-derived BAL cells for migratory phenotype and **DPPC-TC-ONOO^−^** probe activity in murine model of ALI. **(B)** Illustration and timeline of ITB-induced ALI model and LNP instillation. Quadrant plots were constructed from flow cytometry analyses of myeloid-derived BAL cells for migratory phenotype (CD11b+) and **DPPC-TC-ONOO^−^** probe activity. Quadrant plots are shown for individual mice, which were used to quantify **DPPC-TC** as a measure of ONOO^−^ generation. Gating threshold applied for each marker and **DPPC-TC** was based on fluorescent signals of antibody or DPPC-TC-stained and unstained BAL cells.

We employed a previously reported ITB method, which causes a severe inflammatory response characterized by infiltration of immune cells, which is dependent upon iNOS activation.^23^ During *in vivo* ALI, release of cytokines and chemokines, such as tumor necrosis factor-alpha (TNF-α) and interleukin-1 beta (IL-1β), recruit macrophages and other immune cells to the site of injury.^49^ During the initial phase of injury (d0–d3), pro-inflammatory macrophages are dominant, which is associated with higher levels of iNOS expression.^23^ For a generalizable and biocompatible instillation protocol, we first explored LNP concentrations ranging from 0.5 to 2.0 mM. We used LNPs carrying **DPPC-TC** (5 mol% of the lipid content) in C57BL6/J mice and analyzed the viability of the cells from BAL via flow cytometry (Supplemental Figure 16). Results indicated 94% viability at 0.5 mM LNP concentration, which we employed for the subsequent experiments. On d0, mice (C57BL6/J or *Nos2^−/–^*) were anesthetized and received an intratracheal instillation of either PBS (control) or bleomycin (Figure 6A). To detect the ONOO^−^ generation during the early inflammatory response, mice were again anesthetized and instilled with 0.5 mM LNPs containing **DPPC-TC-ONOO^−^**(5 mol% of the lipid content) on day 3. The animals were sacrificed, and BAL was collected for immunostaining 3 hour-post instillation. Flow staining^50^ was used to define myeloid derived cells (CD45+), migratory (CD11b+) and pulmonary in nature (CD11c+) (Supplemental Figure 17). We observed a significant increase in recruited population as a result of ITB for both WT (13 ± 2.4%) and *Nos2^−/–^* (17 ± 6.1%) compared to healthy WT (2 ± 0.8%) (Supplemental Figure 18). This increase was also observed for the migratory, and hence activated, population due to ITB in WT (28 ± 6.6%) and *Nos2^−/–^* (20 ± 8.2%), compared to control WT (2 ± 0.5%) (Supplemental Figure 18). The migratory population was further analyzed for **DPPC-TC** signal quantification (Figure 6B and Table 2). As indicated by the increase in migratory macrophages in both C57BL6/J and *Nos2^−/–^*mice following ITB, inflammatory activation was not altered significantly by the lack of ^•^NO generation. The degree of **DPPC-TC** fluorescence generated in response to ITB in *Nos2^−/–^* (entry 3) was no different from control (entry 1). However, DPPC-TC fluorescence was significantly increased (∼30%) in C57BL6/J with ITB (entry 2). These data indicate that **DPPC-TC-ONOO^−^** was successfully delivered via LNPs to the BAL cells *in vivo* and was able to quantify the production of ONOO^−^ with ALI.

**Table 2.**
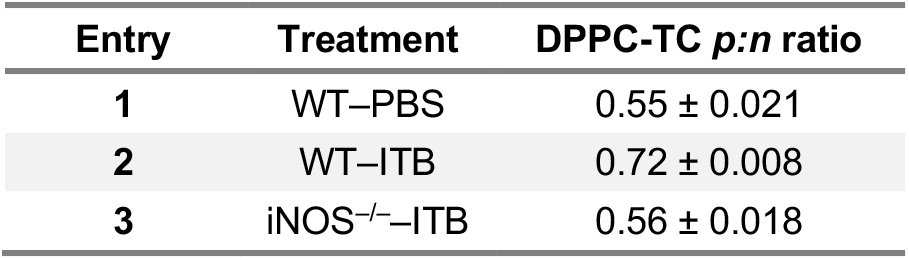
Characterization of the **DPPC-TC** positive-to-negative *(p:n)* ratio in CD11b+ macrophage population from BAL cells (*n* = 3 per group).

| Entry | Treatment | DPPC-TC *p:n* ratio |
| --- | --- | --- |
| 1 | WT-PBS | 0.55 ± 0.021 |
| 2 | WT-ITB | 0.72 ± 0.008 |
| 3 | iNOS<sup>-/-</sup>-ITB | 0.56 ± 0.018 |

## CONCLUSIONS

In situ, real-time imaging of ONOO^−^ in proximity to lipid environments in live mammalian cells and tissues has been a fundamental challenge yet to be addressed. In this work, we describe the development and diverse applications of **DPPC-TC-ONOO^−^**, a membrane-localized and biocompatible probe that allows direct investigation of lipid environments targeted by ONOO^−^. This designer phospholipid and POPC self-assemble to model protocell membranes in the form of GVs. These GVs respond to ONOO^−^ by lighting up at the membrane, remain physically intact, and show excellent selectivity against other RONS. Through cationic/zwitterionic LNPs, **DPPC-TC-ONOO^−^** was delivered into HeLa (Figure 4 and Supplemental Figure 10) and RAW 264.7 (Figure 5 and Supplemental Figure 11). These cell types can generate both NOS-derived ^•^NO and increased oxidative stress upon stimulation. Within HeLa cells the production of these reactive species can be separately stimulated. NOS activation is dependent upon IFNγ/LPS treatment, while NADPH oxidase is stimulated to produce O_2_^•–^ by PMA administration. This was clearly demonstrated via detection of fluorescence signal within just 3 minutes of PMA stimulation following IFNγ/LPS (Figure 4A). The iNOS-mediated production of ONOO^−^ upon LPS treatment results in significant subcellular fluorescence enhancement in RAW 264.7 cells. Importantly, within macrophages oxidative stress results from multiple sources and, thus, the LPS-mediated induction of ^•^NO production is sufficient to produce fluorescence (Figure 5A). Comprehensive colocalization studies indicated that **DPPC-TC-ONOO^−^** localizes primarily in the ER for each cell type. Time-dependent imaging showed that the subcellular detection of ONOO^−^ reactivity is rapid. Of significant note, LNP-mediated delivery plays a predominant role for the localization of **DPPC-TC-ONOO^−^** to the ER, which was supported through the use of LNPs carrying a fluorogenic, control lipid (Supplemental Figure 13). **TEG-TC-ONOO^−^**, the non-amphiphilic analog of **DPPC-TC-ONOO^−^**, exhibits poor localization or site-specificity, especially in RAW 264.7 (Supplemental Figure 13). Additionally, control samples using the iNOS inhibitor 1400W revealed that **DPPC-TC-ONOO^−^** is selective to iNOS. Furthermore, both **DPPC-TC** and **DPPC-TC-ONOO^−^** can localize in the lung lining of bronchioles and parenchyma (Figure 6A). Unlike **TEG-TC-ONOO^−^**, which diffuses non-selectively throughout the lung lining, **DPPC-TC-ONOO^−^** allows for the localized visualization of nitrative stress due to ALI. Qualitative confocal imaging of PCLS incubated with LNPs revealed nitrative stress primarily around bronchioles upon NM exposure. Finally, flow cytometry of activated alveolar macrophages from a murine model of ALI demonstrated ONOO^−^ production upon ITB challenge via fluorogenic activation of **DPPC-TC-ONOO^−^** (Figure 6B and Table 2). Through the use of C57BL6/J and *Nos2^−/–^* mice, both of the *ex vivo* and *in vivo* experiments validated that the function of **DPPC-TC-ONOO^−^** is iNOS-specific.

Interest in expanding the chemical toolbox of synthetic lipids is surging and demand for applying these tools in both basic research and biomedicine is substantial.^51–55^ Our work opens a platform for the development of lipid self-assemblies capable of sensing a chemical cue, offering a new design opportunity for synthetic protocell compartments^56–59^ and a purely chemical perspective to cell-free sensing.^60^ In addition, delivery of unnatural lipids to cells and tissues via LNPs expands the current landscape of soft-material carriers, which are typically employed for small molecules, sugars, peptides, proteins, and nucleic acids.^61^ LNPs containing **DPPC-TC-ONOO^−^** enables real-time imaging of the ER during nitrative stress. This bioimaging tool, in conjunction with LNP delivery, can be leveraged to help untangle redox processes that damage eukaryotic cells. The ER, which functions in protein synthesis and folding,^62^ must maintain its redox homeostasis to facilitate disulfide bridge formation in proteins.^63^ Additionally, the ER membrane is a key site for lipid peroxidation in ferroptosis^64^, a non-apoptotic cell death mechanism characterized by iron-dependent accumulation of lipid peroxides.^65, 66^ These highlight the importance of ER-specific redox probing in live mammalian cells. Finally, compared to the widely used techniques that measure redox species indirectly, the use of LNPs with **DPPC-TC-ONOO^−^** offers a facile and direct approach for investigating nitrative stress both *ex vivo* and *in vivo*. The modular design of **DPPC-TC-ONOO^−^** serves as a blueprint for lipid-based probes of other reactive species and the knowledge gained through our methodology can advance the understanding of redox biochemistry.

## MATERIALS AND METHODS

### Materials

1-2-dipalmitoyl-*rac*-glycerol, copper (II) sulfate (CuSO_4_), 2-{2-[2-(2-propynyloxy)-ethoxy]-ethoxy}ethanol, 1,1,1-trifluoro-4-(4-hydroxyphenyl)butan-2-one, ethylene chlorophosphite, coumarin 343 were purchased from Millipore-Sigma. 3-azido-7-hydroxycoumarin was purchased from Biosynth International Inc. Tris(3-hydroxypropyltriazolyl-methyl)amine (THPTA) was purchased from Click Chemistry Tools. 1-Palmitoyl-2-oleoyl-*sn*-glycero-3-phosphocholine (POPC), 1,2-palmitoyl-*sn*-glycero-3-phosphocholine (DPPC), and 1,2-dioleoyl-*sn*-glycero-3-phosphoethanolamine-*N*-(lissamine rhodamine B sulfonyl) (ammonium salt) were purchased from Avanti Polar Lipids. 3-(4,5-Dimethylthiazol-2-yl)-2,5-diphenyl tetrazolium bromide (MTT) cell proliferation assay kit was purchased from ATCC. Dulbecco’s modified Eagle’s medium/Nutrient Mixture F-12 (DMEM/F-12 HAM), low-gelling temperature agarose, and gentamicin were purchased from Millipore Sigma, penicillin/streptomycin was obtained from (Gibco), LDH kit and WST-1 reagent were purchased from Roche. Bleomycin sulphate was purchased from Santa Cruz Biotechnology, ketamine and xylazine from Fort Dodge Animal Health, Fort Dodge, IA) and eFluor 780-conjugated fixable viability dye from Invitrogen. TruStain FcX anti-mouse CD16/32 antibody (Fc Block), FITC anti-mouse/human CD11b antibody, Alexa Fluor® 647 anti-mouse CD11c antibody, Alexa Fluor® 700 anti-mouse CD45 antibody were purchased from BioLegend. All other chemical materials, including the common reagents for synthesis, buffers, and media, were obtained from commercial sources as detailed in Supporting Information. Milli-Q^®^ water was used in dilutions and preparations of buffers and stock solutions unless otherwise specified.

### Preparation and Usage of Reactive Nitrogen and Oxygen Species

The information given here concerns the fluorescence plate reader measurements and GV-based redox sensing. All of the reactive nitrogen and oxygen species were prepared and handled under oxygen-free conditions by bubbling high-purity N_2_ gas into the reagent solutions and buffers. The stock solutions of ONOO^−^, ^•^NO, H_2_O_2_, and O_2_^•–^ were prepared separately, while ^1^O_2_ and HO^•^ were generated *in situ* in the presence of the probe (**TEG-TC-ONOO^−^** or the GVs containing **DPPC-TC-ONOO^−^**). The stock solutions were used immediately upon preparation. ONOO^−^ solution was prepared according to a previously reported protocol.^67^ In a 20 mL vial, 50 mM of NaNO_2_ (0.5 mL) and 50 mM of H_2_O_2_ (0.5 mL) were stirred vigorously. To this mixture was rapidly added 1 M hydrochloric acid solution (1 mL), which was immediately followed by the addition of 1.5 M NaOH solution (1 mL). The resulting pH depends on how fast the NaOH solution is added following the addition of HCl. For a time gap of 1.5 seconds, the final pH of the ONOO^−^ stock solution was measured as 12.8, with a ONOO^−^ concentration of 55 mM. For a time gap of 18 seconds, the pH was 10.6, with 2.2 mM. The concentration of ONOO^−^ was determined by UV-Vis spectrophotometry, using the previously reported extinction coefficient (ε_302_) of 1670 M^-1^cm^-1^.^68•^ NO solution (1.8 mM) was prepared based on a previously reported protocol.^39^ Specifically, 2.0 g NaNO_2_ and 5.0 g KI were dissolved in 3 mL of water, followed by slow addition of 20 mL of an aqueous H_2_SO_4_ solution (2 M). The evolving gas was passed through a Tygon^®^ tubing filled with NaOH pellets and bubbled into deoxygenated Tris buffer (50 mM, pH 7.5). H_2_O_2_ solution (8 mM) was prepared by diluting a commercial H_2_O_2_ solution (30 w/w%, 9.8 M). O_2_^•–^ solution (8 mM) was prepared from KO_2_ dissolved in DMSO. The KO_2_ powder (2.6 mg) was added into DMSO (2 mL) and the resulting mixture was sonicated until the powder fully dissolved and the solution turned clear. The resulting solution (∼18 mM) was then diluted as necessary. HO^•^ was generated *in situ*, with the probe being present, following a previously reported protocol.^39^ Specifically, we added 2.5 µL of aqueous Fe(ClO_4_)_2_ solution (18 mM) to 147.5 µL mixture of Tris buffer (50 mM, pH 7.5), **TEG-TC-ONOO^−^** (1 µM), and H_2_O_2_ (36 mM). For confocal microscopy of GVs, 5 μL of H_2_O_2_ (16 mM) and 5 μL of iron perchlorate (8 mM) were mixed, which was used as the stock solution used for cell-free assays. ^1^O_2_ was generated *in situ*, with the probe being present, according to a previously reported protocol.^39^ Specifically, we added 2.5 µL of aqueous NaOCl (18 mM) to a 147.5 µL mixture of Tris buffer (50 mM, pH 7.5), **TEG-TC-ONOO^−^** (1 µM), and H_2_O_2_ (36 mM).

### Preparation, Usage, and Confocal Imaging of Giant Vesicles

GVs were prepared from stock solutions of POPC (5 mM), **DPPC-TC-ONOO^−^** (0.25 mM), and Liss-Rhod PE dye (45 µM) in CHCl_3_. A mixture of POPC, **DPPC-TC-ONOO^−^**, and Liss-Rhod PE (98.5: 1.0: 0.5 molar ratio) was taken into a gas tight Hamilton syringe and spread on the conductive side of an indium tin oxide (ITO) coated glass slide (only the central area). The glass slide was then placed under high vacuum for 1 hour, resulting in complete removal of CHCl_3_. The electroformation chamber (Supplemental Figure 7) was assembled by the following steps: A short piece of copper tape was attached to each side of the silicon gasket, which was then placed on the conductive side of an untreated ITO-coated glass. The lipid-containing ITO-coated glass slide was taken from the high vacuum and placed on the top of the silicon gasket. The chamber was clamped with three binder clippers (two for the sides and one for the bottom) to ensure proper sealing. The chamber was then filled with a hydrating buffer while avoiding air bubbles. 250 µL of aqueous sucrose solution (400 mM) was added and the chamber was sealed with a small piece of clay. The sealed chamber was connected through its copper tapes to a function generator using alligator clippers, which completed the electroformation circuit. Electroformation process was carried out at 2 V and 10 Hz for 3 hours. Upon completion, the two side binder clippers were removed and vesicles were harvested with a needle syringe. 50 μL aliquots of the vesicle solution (pH 8.5) were treated with 0.5 μL of the stock solution of the redox agent of choice. The reaction mixture was left undisturbed for 5 minutes. A 5 μL sample of the resulting mixture was then collected and mounted between a cover glass and cover slip (1.2 mm thickness), which was subsequently sealed using a clear nail polish. Z-stack images were acquired in Lightning mode using two tracks: DPPC-TC (405/475 nm) and rhodamine (560/610 nm) channels.

### Protocols for Cell Studies

100k HeLa or RAW264.7 cells were seeded into a 35 mm glass microscope dish and cultured overnight in DMEM with 10% FBS, 37 °C, 5% CO_2_. A mixture of **DPPC-TC-ONOO^−^** (5%), DOTMA (47.5%) and DOPE (47.5%) was dried under reduced pressure, then hydrated with 300 mM sucrose to get a final lipid concentration of 125 µM. Sonication of this mixture over 60 minutes provided LNPs. The LNP solution was then diluted with Opti-MEM™ to a final LNP concentration of 50 µM. This mixture was directly added to live cells grown in glass-bottomed dishes and the cells were incubated at 37 °C for 3 hours for the complete uptake of LNPs. After 3 hours, the cell media was replaced with fresh Opti-MEM media containing appropriate stimulants, and cells were incubated overnight at 37 °C. For endogenous generation of peroxynitrite through stimulation in HeLa cells, a solution of IFN-γ (100 ng/mL) and LPS (1 mg/mL) in Opti-MEM (250 µL) was first added to HeLa cells. The resulting samples was incubated for 12 hours, then the media was decanted. Prior to imaging, a solution of PMA (10 nM) in HBSS (250 µL) was added and the resulting samples were incubated for 60 minutes at 37 °C. For RAW cells, LPS (100 ng/mL) in Opti-MEM (250 µL) was added to the cells and the resulting samples were incubated for 16 hours overnight. For iNOS inhibition in cells, 1400W (20 µM) was introduced into Opti-MEM. Cells were then washed and the organelle tracker, dissolved in HBSS, was added.

### Animal Use, PCLS, Instillation, and BAL Extraction

All experiments were conducted with 6– to 8-week-old C57BL6/J (WT) and iNOS^−/–^ mice bred in-house but originally derived from Jackson Laboratories (Bar Harbor, ME, USA). Food and water were provided *ad libitum* and mice were housed under standard conditions. All experiments were performed in compliance with Rutgers University Institutional Animal Care and the biosafety protocols approved by the institutional biosafety committee following the guidelines by U.S. National Institutes of Health Guide for the Care and Use of Laboratory Animals.

To prepare PCLS, mice were first anesthetized with an intraperitoneal injection of ketamine (135 mg/kg) and xylazine (30 mg/kg) and euthanized via exsanguination. The trachea was cannulated, and 1.5% low-melting point agarose was instilled into the lung and allowed to congeal. The lungs were separated and further embedded in additional agarose for stability when slicing. Slicing was performed using a Krumdieck Tissue Slicer (Alabama Research & Development) to achieve a slice thickness of 300 μm. PCLS were cultured in DMEM/F-12 HAM supplemented with 100 U/mL penicillin/streptomycin and 50 μg/mL gentamicin at 37°C and 5% CO_2_. PCLS were exposed to NM (50 μM mechlorethamine hydrochloride in DMEM/F-12 HAM or DMEM/F-12 HAM alone (control) for 1 hour at 37°C. After 1 day, PCLS were incubated with LNPs (75 μM) for 1 hour, subsequently stained with MitoTracker Red for 2 minutes, and imaged on a Leica TCS SP8 confocal microscope. PCLS with airways of comparable sizes were selected to be imaged. PCLS viability after incubation with LNPs was determined using established lactate dehydrogenase (LDH) and water-soluble tetrazolium salt-1 (WST-1) assays (46). PCLS were prepared from both C57Bl6/J and *Nos2^−/–^*mice.

Mice were anesthetized with isoflurane and received a single 50 µL intratracheal instillation of bleomycin (ITB, 3 U/kg in DMSO/PBS (6:94 v/v)) or DMSO/PBS (6:94 v/v) (control).^69^ To ensure complete dose retention after the instillation, mice were observed for 10 mins post-recovery. Mice were re-anesthetized 72 hours later (d3), and received a 50 µL intratracheal instillation of LNPs (0.5 mM) or sucrose (300 mM, control). The optimal LNP concentration was determined to be 0.5 mM based on a cell viability assay (Supplemental Figure 16). After 3 hours, mice were sacrificed by a single intraperitoneal injection of xylazine (30 mg/kg) and ketamine (135 mg/kg) and exsanguination. Following exposure of the abdominal cavity and thoracotomy, cardiac perfusion was performed (3 mL 1× PBS). 5×1 mL of ice-cold PBS was instilled through a 20 gauge canula to collect BAL fluid. The collected fluid was centrifuged at 300g for 8 minutes to pellet cells for flow cytometry.

### Flow Cytometry

Cells collected from the BAL fluid were diluted in staining buffer (5% FBS in 1× PBS, 0.2% sodium azide) to a volume of 100 µL. samples were treated for 10 minutes at 4 °C with TruStain FcX anti-mouse CD16/32 (Fc Block, 1:100) to inhibit non-specific binding during staining and analysis. Samples were treated with the mixture containing the following antibodies (1:100) for 30 minutes at 4 °C: CD11b, CD11c, and CD45 in dark. Samples were centrifuged for 6 minutes at 400 g, staining buffer was used to wash the cells. Cells were then stained with eFluor 780-conjugated fixable viability dye for 30 minutes at 4 °C. After being fixed with 3% paraformaldehyde for 20 minutes at 4 °C and cells were washed with cold PBS and resuspended in 100 µL PBS. Using a Gallios 10-color flow cytometer, cells were examined. Using Kaluza software, cells were initially sorted based on size and complexity, screened for viability and myeloid-derived origin. They were analyzed for CD11b and CD11c expression (Supplemental Figure 18) and categorized for different phenotypes. CD11b+ population was used for **DPPC-TC** signal quantification (Figure 6B and Table 2).

## ASSOCIATED CONTENT

Additional chemicals; General synthetic procedures; NMR spectra; LCMS analysis; UV and fluorescence spectrophotometric characterizations; preparation and DLS analysis of giant vesicles; staining protocols; MTT assay; supporting confocal images of HeLa and RAW 264.7 cells; Viability and addition confocal data for PCLS; Viability of BAL cells, gating strategy for flow cytometry (PDF).

The Supporting Information is available free of charge at https://pubs.acs.org. Supplemental Figures 1−18 and Supplemental Table 1

## AUTHOR INFORMATION

### Author Contributions

E.C.I. conceived and supervised the study. B.G. and H.E. conducted the syntheses and spectroscopic and spectrophotometric analyses of the probes. B.G. established protocols for the preparation of giant vesicles. B.G., T.A., and M.R.L.S. performed confocal imaging. T.A., M.R.L.S., and C.G. conducted cell work. T.A. performed the p*K*_a_ measurement, MTT assay, and flow cytometry (with assistance from E.R.S.). A.B. and T.A. performed *ex vivo* work. E.A. and E.R.S. performed *in vivo* experiments. A.J.G. oversaw *ex vivo* and *in vivo* studies. All of the authors contributed to the interpretation of data. E.C.I. wrote the manuscript with input from all authors.

### Notes

E.C.I., B.G., and H.E. are co-inventors of a provisional patent application filed by Rutgers University on the subject of this work.

## ACKNOWLEDGMENT

We thank G. Hall for allowing access to the fluorometer. We thank K-B. Lee for allowing access to the microplate reader and we acknowledge M. Chen and L. Wang for their assistance with fluorescence assays. We thank D. L. Laskin and K. Hast for useful discussions. This work was supported by the US National Institutes of Health / National Institute of Biomedical Imaging and Bioengineering, Trailblazer Award (EB029548), the Rutgers Center for Lipid Research, the American Cancer Society, Institutional Research Grant Early Investigator Award, and the Rutgers Cancer Institute of New Jersey NCI Cancer Center Support Grant (P30CA072720) (to E.C.I.); Steven A. Cox Scholarship for Cancer Research (to T.A.); National Heart, Lung, and Blood Institute (HL086621) (to A.J.G.); and National Institute of Environmental Health Sciences (training grant ES007148 for E.R.S. and program grant ES005022).

## Notes

### Summary of Updates

1. Conflict of interest statement 2. Author list 3. Biological studies, which now include ex vivo and in vivo experiments 4. Title

