## Supplementary Information for "Direct Assessment of Nitrative Stress in Lipid Environments: Applications of a Designer Lipid-Based Biosensor for Peroxynitrite"

| Contents | Page |
|---|---|


**Chemicals** **S4**
**General Synthetic Methods** **S5**
**Preparation Procedures and Characterization Data for Small Molecules** **S6–S33**
Compound **3** S6
**TEG-TC-ONOO$^-$** S7
Compound **S2** S8
Compound **4** S9
**DPPC-TC-ONOO$^-$** S10
**TEG-TC** S11
**DPPC-TC** S12
$^{1}$H, $^{13}$C, $^{19}$F, $^{31}$P NMR Spectra S13–S33
**LCMS Analysis of Reaction of TEG-TC-ONOO$^-$ with ONOO$^-$** **S34**
Supplemental Figure 1
**Determination of p$K_a$ of TEG-TC** **S35**
Supplemental Figure 2
**pH dependence of fluorescence intensity for TEG-TC** **S36**
Supplemental Figure 3
**Determination of Relative Fluorescence Quantum Yield** **S37**
**Microplate Fluorescence Measurements** **S37**
**Spectrophotometric Characterizations by Absorbance and Emission** **S38**
Supplemental Figure 4
**UV-Vis Spectrophotometric Investigation of Peroxynitrite** **S39**
Supplemental Figure 5
**Effect of Peroxynitrite on TEG-TC-ONOO–** **S40**
Supplemental Figure 6
**Preparation and DLS Analysis of Giant Vesicles** **S41**
Supplemental Figures 7 and 8
**Staining Cells with Organelle Trackers or Actin Dye** **S42**
**MTT Assay for HeLa and RAW 264.7** **S43**
Supplemental Figure 9 and Supplemental Table 1
**Supporting Confocal Images (for HeLa and RAW 264.7)** **S44–S47**
Supplemental Figures 10–13
**Cellular Viability in PCLS Using LDH Leakage and WST-1 Reduction** **S48**
Supplemental Figure 14
**Supporting Confocal Images of PCLS** **S49**
Supplemental Figure 15
**Viability Analysis of BAL Cells** **S50**
Supplemental Figure 16

**Gating Strategy for Flow Cytometry Data** **S51**
Supplemental Figure 17
**CD11b versus CD11c Expression** **S52**
Supplemental Figure 18
**References** **S52**

### Chemicals

Reagents: 1-2-dipalmitoyl-*rac*-glycerol, pyridine, and 7 M ammonium hydroxide, triethylamine (TEA), dimethyl sulfoxide [(DMSO), molecular biology grade], copper (II) sulfate ($CuSO_4$), 2-{2-[2-(2-propynyloxy)ethoxy]-ethoxy}ethanol, hydrogen peroxide ($H_2O_2$), sodium hydroxide (NaOH), sodium nitrite ($NaNO_2$), potassium superoxide ($KO_2$), sodium ascorbate, 1,1,1-trifluoro-4-(4-hydroxyphenyl)butan-2-one, triphosgene, Oxone, sucrose, ethylene chlorophosphite, triphenylphosphine oxide ($Ph_3PO$), sodium bicarbonate ($NaHCO_3$), Coumarin 343, and sodium sulfate ($Na_2SO_4$) were purchased from Millipore-Sigma. 3-azido-7-hydroxycoumarin was purchased from Biosynth International Inc. Tris(3-hydroxypropyltriazolyl-methyl)amine (THPTA) was purchased from Click Chemistry Tools. 1-Palmitoyl-2-oleoyl-*sn*-glycero-3-phosphocholine (POPC), 1,2-palmitoyl-*sn*-glycero-3-phosphocholine (DPPC), and 1,2-dioleoyl-*sn*-glycero-3-phosphoethanolamine-*N*-(lissamine rhodamine B sulfonyl) (ammonium salt) (here referred to as Liss-Rhod PE) were purchased from Avanti Polar Lipids. 3-(4,5-Dimethylthiazol-2-yl)-2,5-diphenyl tetrazolium bromide (MTT) cell proliferation assay kit was purchased from ATCC.

Buffers, Solvents, Media: Phosphate-buffered saline (PBS), 4-morpholineethanesulfonic acid (MES), tris(hydroxymethyl)aminomethane (Tris), dichloromethane ($CH_2Cl_2$), ethyl acetate (EtOAc), benzene, toluene, tetrahydrofuran (THF), and chloroform ($CHCl_3$) were purchased from Millipore Sigma. Acetonitrile ($CH_3CN$) and methanol (MeOH, HPLC grade) were purchased from Fisher Scientific. Deuterated solvents were purchased from either Cambridge Isotope Laboratories or Millipore Sigma. Deuterated solvents contained 0.05% (v/v) TMS as a secondary internal reference. Water was deionized and filtered to a resistivity of 18.2 ΩM with a Milli-Q® Plus water purification system (Millipore, Massachusetts). Buffers were prepared freshly in Milli-Q® water and their pH were adjusted using HCl or NaOH using a Thermo Scientific Orion Star pH meter. Dulbecco's Modified Eagle Medium (DMEM) with and without phenol red was purchased from Corning. Hanks' Balanced Salt Solution (HBSS) and fetal bovine serum (FBS) were purchased from VWR International.

Chemicals used for the animal work were outlined in Materials and Methods in the main manuscript.

### General Synthetic Methods

All reactions were performed under a dry nitrogen atmosphere unless otherwise stated. All glassware was oven-dried before use. Purification of the synthesized compounds was performed using a Büchi Reveleris® flash chromatography system equipped with a FlashPure EcoFlex Diol (50 µm spherical) column or a silica (50 µm irregular) column. Nuclear magnetic resonance (NMR) spectroscopic analyses was carried out using a Bruker Avance Neo 500 MHz spectrometer. NMR data is provided for new compounds. $^{1}$H NMR spectra were acquired at 500 MHz, $^{13}$C NMR spectra were acquired at 126 MHz, $^{31}$P NMR spectra were acquired at 202 MHz, and $^{19}$F NMR spectra were acquired at 471 MHz. Chemical shifts (δ) for $^{1}$H NMR spectra were referenced to $(C\boldsymbol{H}_3)_4Si$ at δ = 0.00 ppm or to $C\boldsymbol{H}Cl_3$ at δ = 7.26 ppm. $^{13}$C NMR spectra were referenced to $\boldsymbol{C}DCl_3$ at δ = 77.23 ppm. $^{31}$P NMR spectra were referenced externally to $Ph_3PO$ dissolved in $CDCl_3$ at δ = 29.06 ppm. The external reference solution was placed in a co-axial NMR tube (~0.1 mL effective volume), which was then inserted into the NMR tube containing the analyte solution (~0.4 mL). The following abbreviations are used to describe $^{1}$H NMR resonances: s (singlet), d (doublet), t (triplet), m (multiplet), dd (doublet of doublets), br (broad), and app (apparent). Coupling constants (*J*) are reported in Hz. Liquid chromatography followed by high-resolution mass spectroscopy (LC-HRMS) analysis from electrospray ionization (ESI) was carried out on a Waters Acquity-Xevo G2-XS QTof instrument. Low-resolution mass spectroscopy (LRMS) analysis was performed using a Finnigan LCQ™ DUO mass spectrometer. Both fluorescence and UV-Vis absorbance measurements were conducted using a Spectra Max ID3 instrument (Molecular Devices). Dynamic light scattering (DLS) measurements were carried out on a Zetasizer Nano C (Malvern Panalytical). Confocal fluorescence microscopy was performed on a Leica TCS SP8 microscope equipped with a 63x, 1.40 NA oil immersion objective and a Tokai Hit stage-top incubator. Images were processed using LasX Lightning deconvolution. All statistical analyses were performed using GraphPad Prism software version 9.

### Preparation Procedures and Characterization Data for Small Molecules

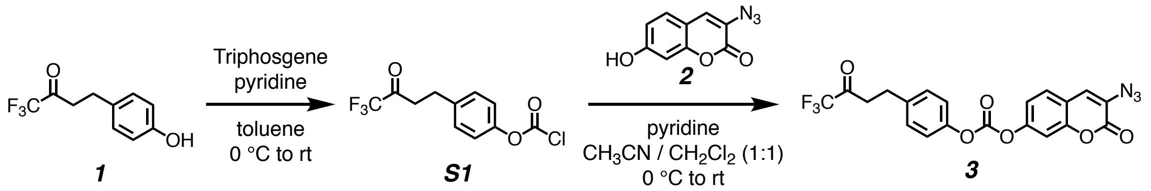

**Synthesis of 3-azido-2-oxo-2*H*-chromen-7-yl (4-(4,4,4-trifluoro-3-oxobutyl)phenyl) carbonate (compound 3):** To a dry 5-mL round-bottom flask was added triphosgene (41 mg, 0.140 mmol, 0.60 equiv) and 0.5 mL of toluene under a nitrogen-rich atmosphere. The solution was cooled to 0 °C and pyridine (28 μL, 0.340 mmol, 1.5 equiv) in 0.5 mL of toluene, followed by 1,1,1-trifluoro-4-(4-hydroxyphenyl)butan-2-one (50 mg, 0.230 mmol, 1.0 equiv) in 1.0 mL of toluene were added dropwise. The reaction mixture was allowed to stir at room temperature (rt) and monitored by TLC until completion (3 h). The resulting mixture was concentrated under reduced pressure, affording the chloroformate intermediate **S1**. The thick residue was re-dissolved in 1.0 mL of $CH_2Cl_2$ and cooled to 0 °C while being stirred vigorously and purged with nitrogen gas continuously. To this mixture, 3-azido-7-hydroxycoumarin (**2**) (32 mg, 0.160 mmol, 1.0 equiv) in 0.5 mL $CH_3CN$ and pyridine (25 μL, 0.460 mmol, 2.0 equiv) in 0.5 mL of $CH_2Cl_2$ were added dropwise. The reaction was allowed to stir at rt for 4 h. The crude mixture was then concentrated to an oil under reduced pressure and purified by silica gel column chromatography (mobile phase: $CH_2Cl_2$ / MeOH, step gradient from 0 to 2% MeOH). The product fractions were collected and concentrated under reduced pressure, providing the carbonate **3** (43 mg, 60%) as a white solid.

**$^{1}$H NMR** (500 MHz, $CDCl_3$): δ 7.47-7.45 (d, *J* = 8.6 Hz, 1H), 7.33 (d, *J* = 2.3 Hz, 1H), 7.28 (s, 1H), 7.27–7.21 (m, 5H), and 3.06–3.02 (m, 4H).

**$^{13}$C NMR** (126 MHz, $CDCl_3$): δ 190.3 (C=O), 157.0 (C=O), 151.8, 151.6, 151.4, 149.5, 137.7, 129.6, 129.5, 128.0, 126.4, 124.9, 121.2, 118.3, 117.5, 109.6, 37.9, and 27.7.

**$^{19}$F NMR** (471 MHz, $CDCl_3$): δ -79.2 (s).

**HRMS** (ESI) m/z: Calculated for $C_{20}H_{11}F_3N_3O_6^-$, $[M – H]^-$, requires 446.0605; found 446.0668.

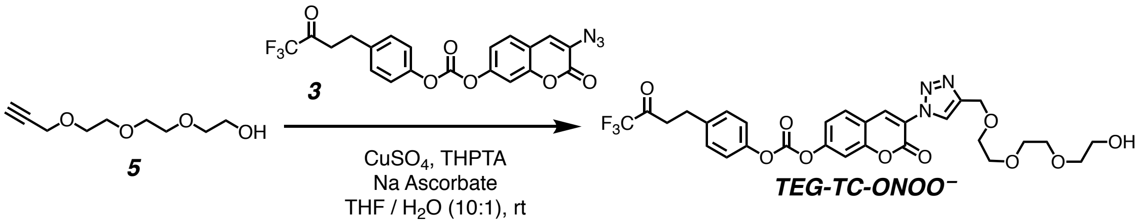


**Synthesis of 3-(4-((2-(2-(2-hydroxyethoxy)ethoxy)ethoxy)methyl)-1*H*-1,2,3-triazol-1-yl)-2-oxo-2*H*-chromen-7-yl (4-(4,4,4-trifluoro-3-oxobutyl)phenyl) carbonate (TEG-TC-ONOO⁻):** To a stirred mixture of propargyl-TEG-OH (**5**) (3.2 mg, 0.017 mmol, 1.5 equiv) and the carbonate **3** (5 mg, 0.011 mmol, 1.0 equiv) in 0.3 mL of THF at rt, was added a mixture of copper (II) sulfate (1.0 mg, 0.006 mmol, 0.5 equiv), THPTA (4.8 mg, 0.011, 1.0 equiv), and sodium ascorbate (1.1 mg, 0.006 mmol, 0.5 equiv) in 0.4 mL of $H_2O$. The reaction was stirred at rt for 24 h under a nitrogen atmosphere. The crude product was concentrated under reduced pressure and purified by silica gel column chromatography (mobile phase: $CH_2Cl_2$ / MeOH, step gradient from 0 to 5% MeOH). The product fractions were collected and concentrated under reduced pressure, providing **TEG-TC-ONOO⁻** (4 mg, 57 %) as a clear film.

**$^1$H NMR (**500 MHz, $CDCl_3$): δ 8.70 (s, 1H), 8.60 (s, 1H), 7.74–7.72 (d, *J* = 2.4 Hz, 2H), 7.45 (d, *J* = 8.5 Hz, 1H), 7.38–7.36 (dd, *J* = 8.7, 2.2 Hz, 1H), 7.29–7.23 (m, 4H), 4.80 (s, 2H), 3.76–3.62 (m, 12H), and 3.07–3.03 (m, 4H)

**$^{13}$C NMR** (126 MHz, $CDCl_3$): δ 190.3, 155.4, 153.8, 153.1, 151.1, 149.4, 145.5, 137.8, 132.4, 129.9, 129.6, 129.5, 123.9, 122.8, 121.0, 118.8, 116.2, 109.7, 72.5, 70.7, 70.6, 70.4, 69.9, 64.4, 61.8, 37.9, and 27.7.

**$^{19}$F NMR** (471 MHz, $CDCl_3$): δ -79.2 (s).

**LRMS** (ESI) m/z: Calculated for $C_{29}H_{27}F_3N_3O_{10}^-$, $[M – H]^-$, requires 634.17; found 634.19.

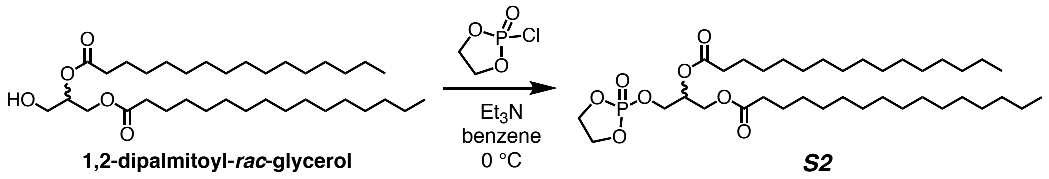

**Synthesis of 3-((2-oxido-1,3,2-dioxaphospholan-2-yl)oxy)propane-1,2-diyl dipalmitate (compound S2):** To a dry 25-mL round-bottom flask was added 1-2-dipalmitoyl-*rac*-glycerol, (500 mg, 0.850 mmol, 1.0 equiv) in 5.0 mL of benzene under a dry nitrogen atmosphere. The mixture was brought to 0 °C and TEA (0.50 mL, 3.75 mmol, 4.4 equiv) was added dropwise. After 15 minutes of stirring, ethylene chlorophosphate (0.75 mL, 8.16 mmol, 9.6 equiv) was added dropwise. The reaction was allowed stirred at rt for 24 h. The solution was quenched with saturated $NaHCO_3$ and washed with EtOAc three times. The collected organic layers were dried with sodium sulfate and concentrated under reduced pressure. The cyclic phosphate triester product **S2** (44 mg, 93%) was obtained as a white solid and used without further purification.

**$^{1}$H NMR (**500 MHz, $CDCl_3$): δ 5.19–5.17 (p, *J* = 4.8 Hz, 1H), 4.39–4.35 (dd, *J* = 11.8, 3.9 Hz, 2H), 4.34–4.12 (m, 5H), 4.11–4.08 (dd, *J* = 11.5, 6.0 Hz, 1H), 2.28 (t, *J* = 7.5, Hz, 2H), 2.25 (t, *J* = 7.5, Hz, 2H), 1.55–1.54 (br s, 4H), 1.21–1.19 (br s, 48H), and 0.82–0.80 (app t, *J* = 6.7 Hz, 6H).

**$^{13}$C NMR** (126 MHz, $CDCl_3$): δ 173.4 (C=O), 173.1 (C=O), 69.5, 69.4, 66.8, 66.7, 66.22 (overlapped), 66.20 (overlapped), 61.5, 34.3, 34.2, 32.1, 29.9 (overlapped), 29.83, 29.82, 29.80, 29.78, 29.65, 29.64, 29.5, 29.45, 29.43, 29.3, 29.2, 25.0, 24.9, 24.8, 22.9, and 14.3.

**$^{31}$P NMR** (202 MHz, $CDCl_3$): δ 17.74 (s).

**LRMS** (ESI) m/z: Calculated for $C_{37}H_{71}O_8PNa^+$, $[M + H]^+$, requires 697.48; found 697.50.

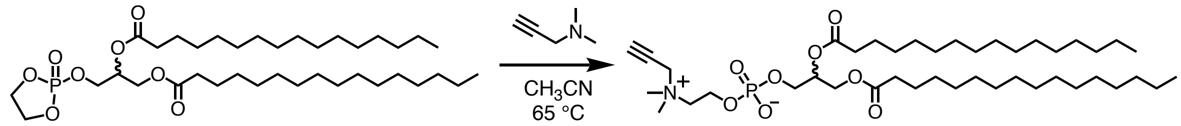


***S2*** ***4***

**Synthesis of 2,3-bis(palmitoyloxy)propyl (2-(dimethyl(prop-2-yn-1-yl)ammonio)ethyl) phosphate (the alkynyl lipid 4):** In a dry 10-mL pressure tube, **S2** (100 mg, 0.160 mmol, 1 equiv) in 2.0 mL of dry $CH_3CN$ was added. 3-Dimethylamino-1-propyne (540 mg, 6.40 mmol, 40 equiv) in 3 mL of dry $CH_3CN$ were added to a dry 5 mL round bottom with sodium sulfate, bubbled with nitrogen and stirred for 30 mins. The mixture was then transferred to the solution containing **S2** and sealed and stirred for 60 h under a dry nitrogen environment at 80°C. The reaction was diluted in $CHCl_3$, filtered, and concentrated to an oil under reduced pressure. The crude product was purified by silica gel chromatography with a diol column (mobile phase: $CHCl_3$ / MeOH / 7 M $NH_4OH$, step gradient from 0 to 25% MeOH / 7 M $NH_4OH$ mixture, 6:1 v/v). The product fractions were collected and concentrated under reduced pressure, providing the alkynyl lipid **4** (49 mg, 58%) as a white solid.

**$^1$H NMR** (500 MHz, $CDCl_3$): δ 5.22 (m, 1H), 4.69 (s, 2H), 4.42–4.39 (dd, *J* = 12.0, 3.1 Hz, 1H), 4.33 (br s, 1H), 4.14 (dd, *J* = 12.0, 7.1 Hz, 1H), 3.96–3.92 (br s, 1H), 3.90–3.87 (br s, 2H), 3.43 (s, 6H), 2.93 (s, 1H), 2.30 (t, *J* = 7.5 Hz, 2H), 2.28 (t, *J* = 7.5 Hz, 2H), 1.52–1.51 (br s, 4H), 1.18 (br s, 48H), and 0.88 (app t, *J* = 7.0 Hz, 6H).

**$^{13}$C NMR** (126 MHz, $CDCl_3$): δ 173.8 (C=O), 173.4 (C=O), 81.5, 72.3, 70.8, 64.4, 63.7, 63.1, 59.4, 55.5, 51.8, 34.6, 34.4, 32.2, 30.0 (overlapped), 29.92, 29.89, 29.78, 29.76, 29.59 (overlapped), 29.56, 29.41, 29.38, 27.8, 25.2, 25.1, 22.9, and 14.3.

**$^{31}$P NMR** (202 MHz, $CDCl_3$): δ -0.73 (s).

**HRMS** (ESI) m/z: Calculated for $C_{42}H_{81}NO_8P^+$, $[M + H]^+$, requires 758.5694; found 758.5727.

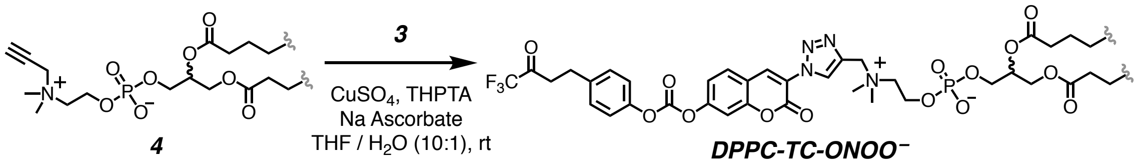


**Synthesis of the DPPC-TC-ONOO⁻:** To a stirred solution of the alkynyl lipid **4** (15.4 mg, 0.034 mmol, 1.3 equiv) and the coumarin azide **3** (20.0 mg, 0.026 mmol, 1.0 equiv) in 0.5 mL of THF were added in 1.3 mL of THF / $H_2O$ (10:1, v/v). The mixture was bubbled with nitrogen before adding copper sulfate (2.0 mg, 0.013 mmol, 0.5 equiv), THPTA (5.6 mg, 0.013, 0.5 equiv), and sodium ascorbate (5.2 mg, 0.026 mmol, 1.0 equiv) into the reaction vessel. After being stirred for 24 h, the reaction mixture was concentrated under reduced pressure and the crude product was purified by silica gel chromatography with a diol column (mobile phase: $CH_2Cl_2$ / MeOH / $H_2O$, step gradient from 0 to 35% MeOH / $H_2O$ mixture, 30:1 v/v). The product fractions were collected and concentrated under reduced pressure, providing **DPPC-TC-ONOO⁻** (8.3 mg, 43%) as a clear film.

$^{1}$**H NMR** (500 MHz, $CDCl_3$ / $CD_3OD$, 3:1 v/v) δ 9.11 (s, 1H), 8.67 (s, 1H), 7.86–7.85 (d, *J* = 8.4 Hz, 1H), 7.55 (s 1H), 7.44 (s, 1H), 7.31–7.29 (d, *J* = 8.5 Hz, 2H), 7.24–7.22 (d, *J* = 8.3 Hz, 2H), 5.30–5.18 (m, 1H), 4.86 (s, 2H), 4.43 (dd, *J* = 12.1, 3.2 Hz, 1H), 4.36 (bs, 2H), 4.17 (dd, *J* = 12.0, 6.9 Hz, 1H), 4.04 (bs, 2H), 3.66 (bs, 2H), 3.41–3.39 (m, 4 H), 3.25 (s, 6H), 2.32 (m, 4H), 1.59 (bs, 4H), 1.26 (bs, 48H), and 0.88 (t, *J* = 7.0 Hz, 6H).

$^{13}$**C NMR** (126 MHz, $CDCl_3$ / $CD_3OD$, 3:1 v/v) δ 177.9, 177.5, 159.5, 158.3, 157.4, 155.3, 152.9, 144.2, 139.6, 138.3, 134.3, 133.4, 133.1, 126.3, 124.6, 123.0, 119.9, 113.7, 74.3, 68.2, 67.5, 66.6, 63.1, 62.4, 55.2, 39.5, 38.1, 37.9, 35.8, 33.53 (overlapped), 33.51, 33.49 (overlapped), 33.38, 33.36, 33.19 (overlapped), 33.15, 32.98, 32.96, 31.9, 28.8, 28.7, 26.5, and 17.7.

$^{31}$**P NMR** (202 MHz, $CDCl_3$ / $CD_3OD$, 3:1 v/v) δ -4.38 (s).

$^{19}$**F NMR** (471 MHz, $CDCl_3$ / $CD_3OD$, 3:1 v/v): δ -77.6 (s).

**HRMS** (ESI) m/z: Calculated for $C_{62}H_{93}F_3N_4O_{14}P^+$, $[M + H]^+$, requires 1205.6373; found 1205.6379.

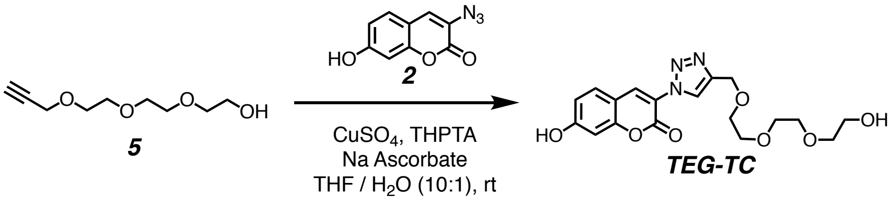


**Synthesis of 7-hydroxy-3-(4-((2-(2-(2-hydroxyethoxy)ethoxy)ethoxy)methyl)-1*H*-1,2,3-triazol-1-yl)-2*H*-chromen-2-one (TEG-TC):** A stirred solution of propargyl-TEG-OH (**5**) (83.2 mg, 0.44 mmol, 1.50 equiv) and 3-azido-7-hydroxycoumarin (**2**) (60.0 mg, 0.295 mmol, 1.00 equiv) in 1.0 mL of THF was bubbled with nitrogen before adding copper sulfate (29.0 mg, 0.15 mmol, 0.50 equiv) and THPTA (65.1 mg, 0.15 mmol, 0.50 equiv) in 0.5 mL of water. Mixture was bubbled with nitrogen again, and sodium ascorbate (59.4 mg, 0.3 mmol, 1.00 equiv) was added in 0.5 ml of water. After being stirred for 12 h, the reaction mixture was concentrated under reduced pressure and the crude product was purified by silica gel column chromatography (mobile phase: $CH_2Cl_2$ / MeOH, step gradient from 0 to 5% MeOH). The product fractions were collected and concentrated under reduced pressure, providing **TEG-TC** (63 mg, 55%) as a clear film.

**$^1$H NMR** (500 MHz, $CD_3OD$) δ 8.57 (s, 1H), 8.49 (s, 1H), 7.64 (d, *J* = 8.6 Hz, 1H), 6.89 (dd, *J* = 8.5, 2.3 Hz, 1H), 6.81 (d, *J* = 2.2 Hz, 1H), 4.73 (s, 2H), 3.79–3.61 (m, 10H), and 3.55 (m, 2H)

**$^{13}$C NMR** (126 MHz, $CD_3OD$) δ 163.5, 156.8, 155.3, 144.6, 135.9, 130.5, 124.6, 119.2, 114.4, 110.4, 102.1, 72.3, 70.2, 70.1, 70.0, 69.4, 63.5, and 60.8.

**HRMS** (ESI) m/z: Calculated for $C_{18}H_{20}N_3O_7^-$, $[M – H]^-$, requires 390.1307; found 390.1308.

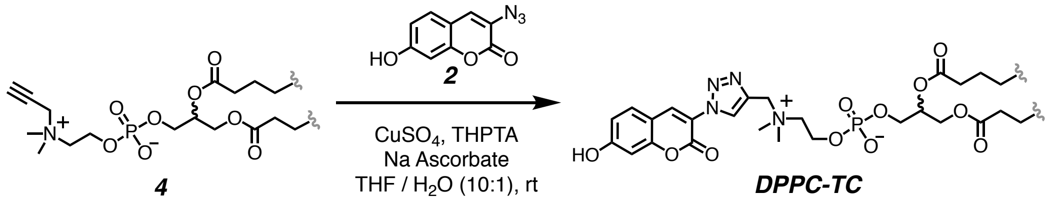


**Synthesis of DPPC-TC:** A stirred solution of the alkynyl lipid **4** (42.0 mg, 0.055 mmol, 1.0 equiv) and 3-azido-7-hydroxycoumarin (**2**) (14.6 mg, 0.072 mmol, 1.3 equiv) in 2.0 mL of THF was bubbled with nitrogen before adding copper sulfate (4.4 mg, 0.027 mmol, 0.5 equiv) and THPTA (12.8 mg, 0.027 mmol, 0.5 equiv) in 0.2 mL of water. Mixture was bubbled with nitrogen again, and sodium ascorbate (11.0 mg, 0.055 mmol, 1.0 equiv) was added in 0.1 mL of water. After being stirred for 12 hours, the reaction mixture was concentrated under reduced pressure and the crude product was purified by silica gel chromatography with a diol column (mobile phase: $CH_2Cl_2$ / MeOH / $H_2O$, step gradient from 0 to 35% MeOH / $H_2O$ mixture, 30:1 v/v). The product fractions were collected and concentrated under reduced pressure, providing **DPPC-TC** (30.1 mg, 57%) as a brown solid.

**$^1$H NMR** (500 MHz, $CDCl_3$ / $CD_3OD$, 3:1 v/v) δ 9.00 (s, 1H), 8.50 (s, 1H), 7.57 (d, *J* = 8.7 Hz, 1H), 6.94 (dd, *J* = 8.6, 2.4 Hz, 1H), 6.89 (d, *J* = 2.4 Hz, 1H), 5.30–5.18 (m, 1H), 4.84 (s, 2H), 4.43 (dd, *J* = 12.1, 3.2 Hz, 1H), 4.36 (bs, 2H), 4.17 (dd, *J* = 12.0, 6.9 Hz, 1H), 4.03 (t, *J* = 6.3 Hz, 2H), 3.65 (bs, 2H), 3.25 (s, 6H), 2.32 (m, 4H), 1.59 (bs, 4H), 1.26 (bs, 48H), and 0.88 (t, *J* = 7.0 Hz, 6H)

**$^{13}$C NMR** (126 MHz, $CDCl_3$ / $CD_3OD$, 3:1 v/v) δ 177.9, 177.5, 167.4, 160.6, 159.2, 140.2, 139.2, 134.5, 132.8, 122.5, 119.0, 114.2, 106.8, 74.2, 68.2, 67.6, 66.6, 63.3, 62.8, 55.2, 38.1, 38.0, 35.8, 33.6, 33.5, 33.39, 33.37, 33.21, 33.17, 33.0, 32.98, 28.8, 28.7, 26.5, and 17.8.

**$^{31}$P NMR** (202 MHz, $CDCl_3$ / $CD_3OD$, 3:1 v/v) δ -0.60 (s).

**HRMS** (ESI) m/z: Calculated for $C_{51}H_{84}N_4O_{11}P^-$, $[M – H]^-$, requires 959.5880; found 959.5889.

Compound ***3*** ([1]H NMR: 500 MHz, $CDCl_3$)

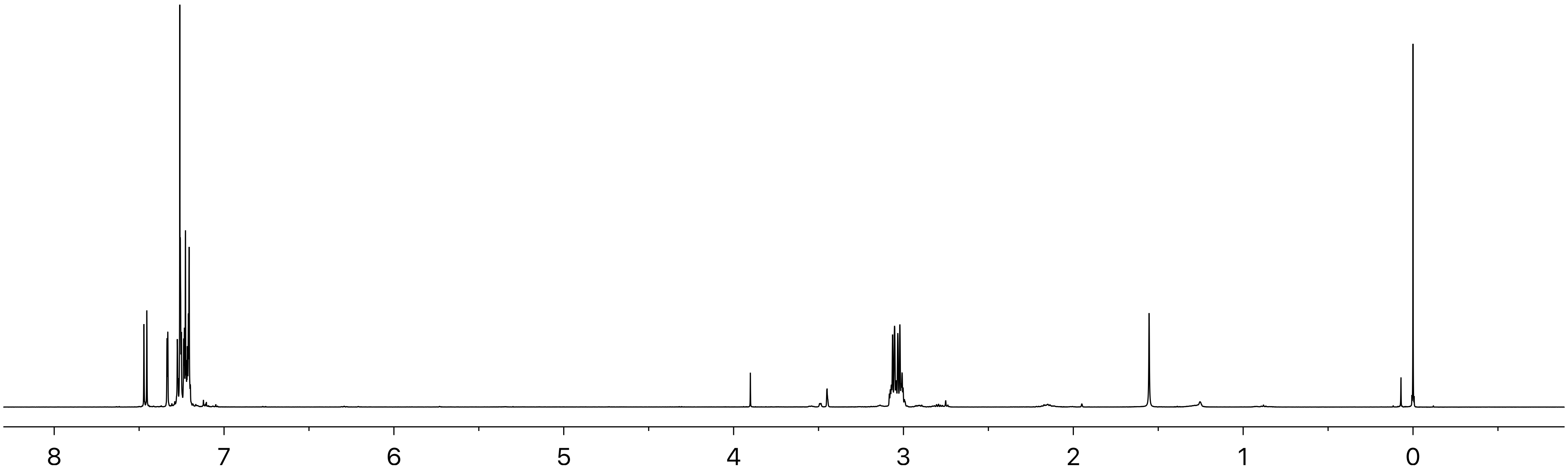

Compound ***3*** ($^{13}$C NMR: 126 MHz, $CDCl_3$)

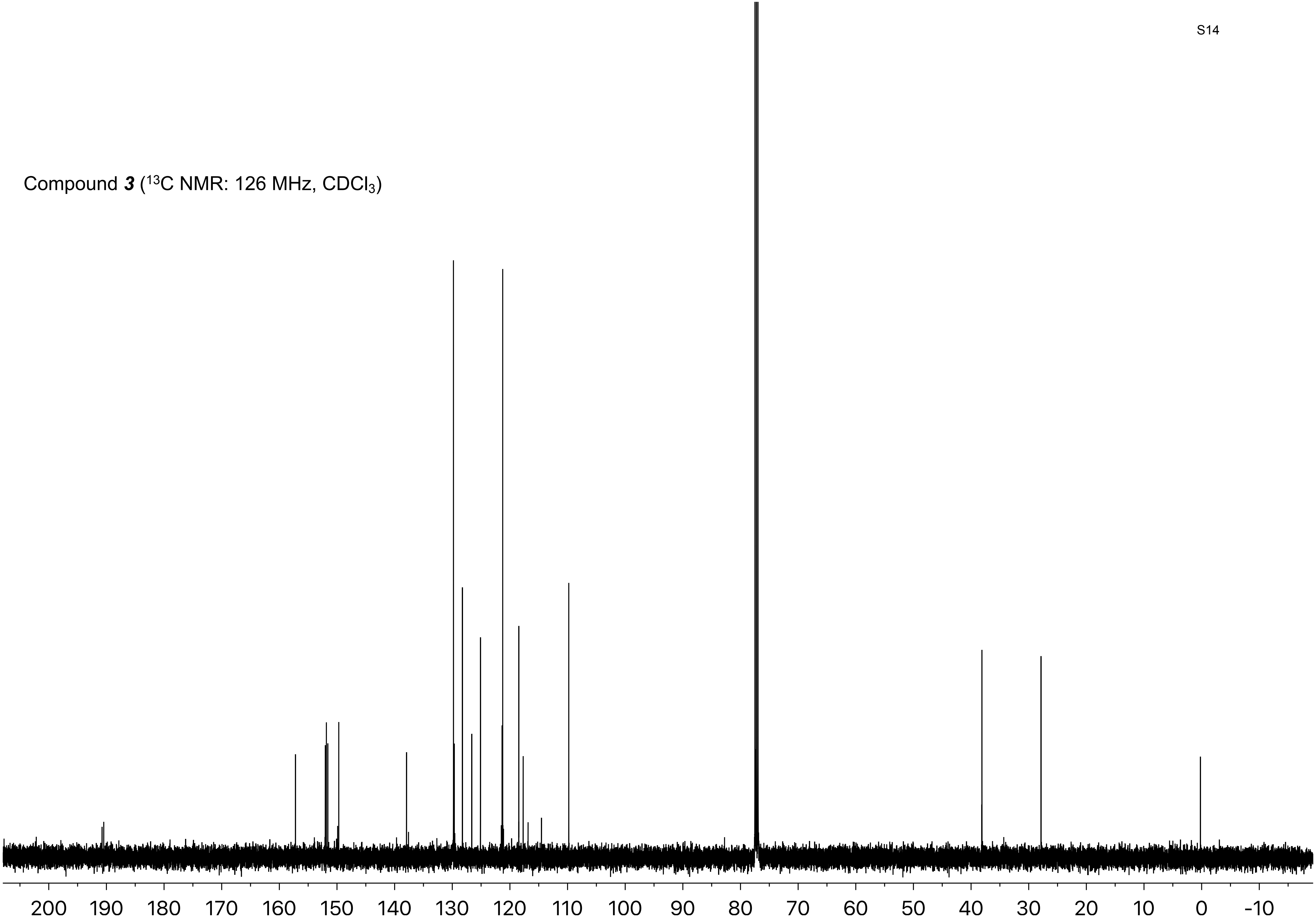

Compound ***3*** ($^{19}$F NMR: 471 MHz, $CDCl_3$)

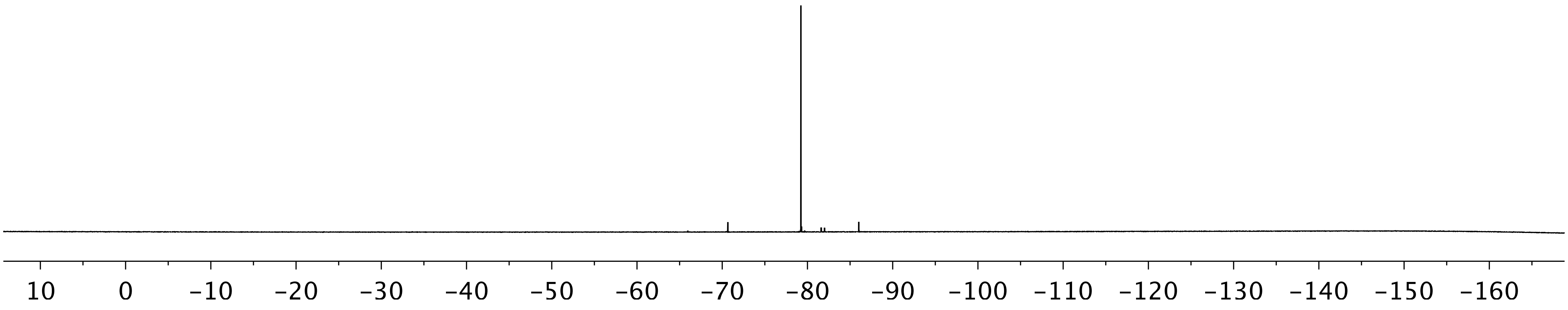

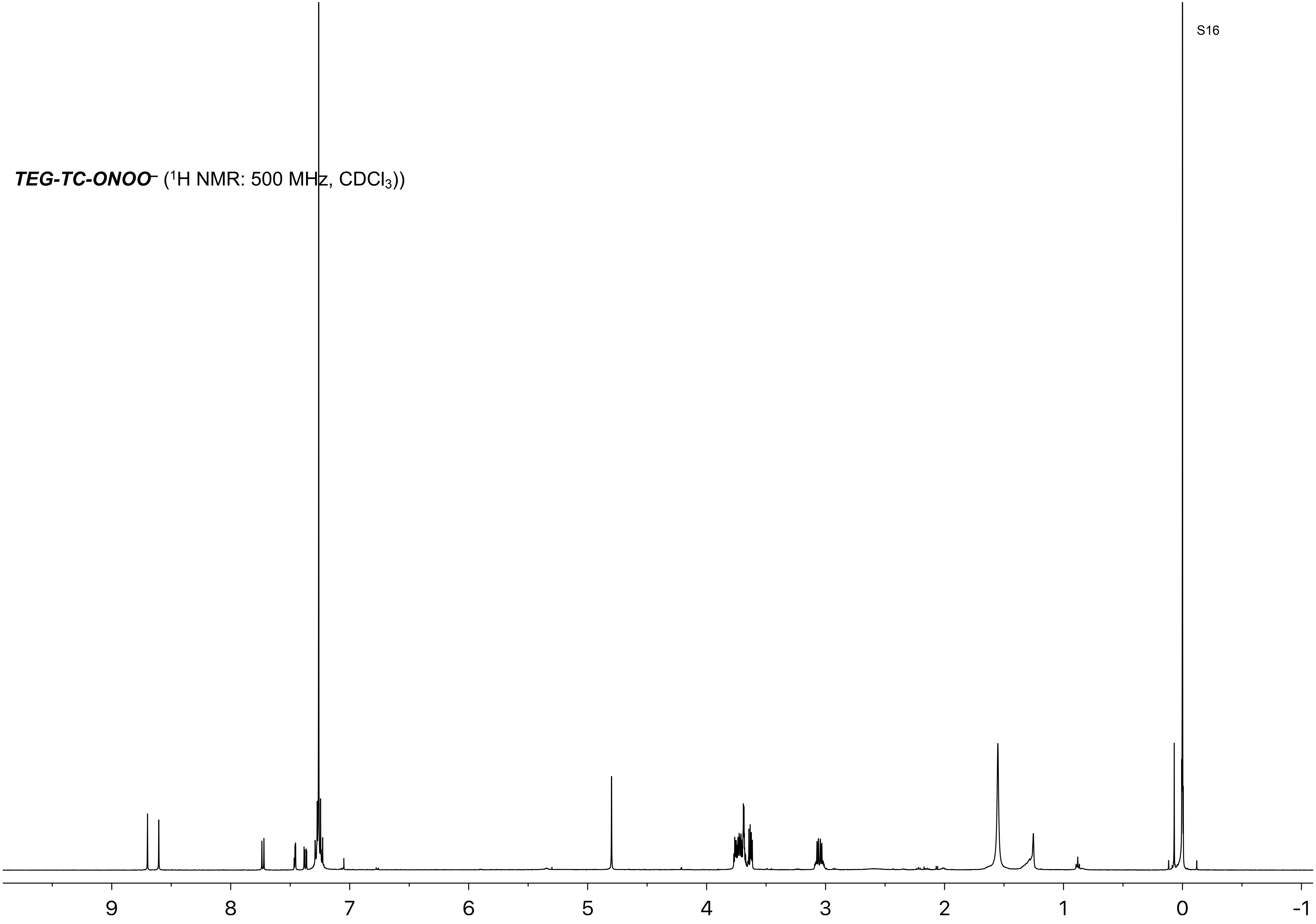
***TEG-TC-ONOO⁻*** ($^1$H NMR: 500 MHz, $CDCl_3$))
9
8
7
6
5
4
3
2
1
0
-1

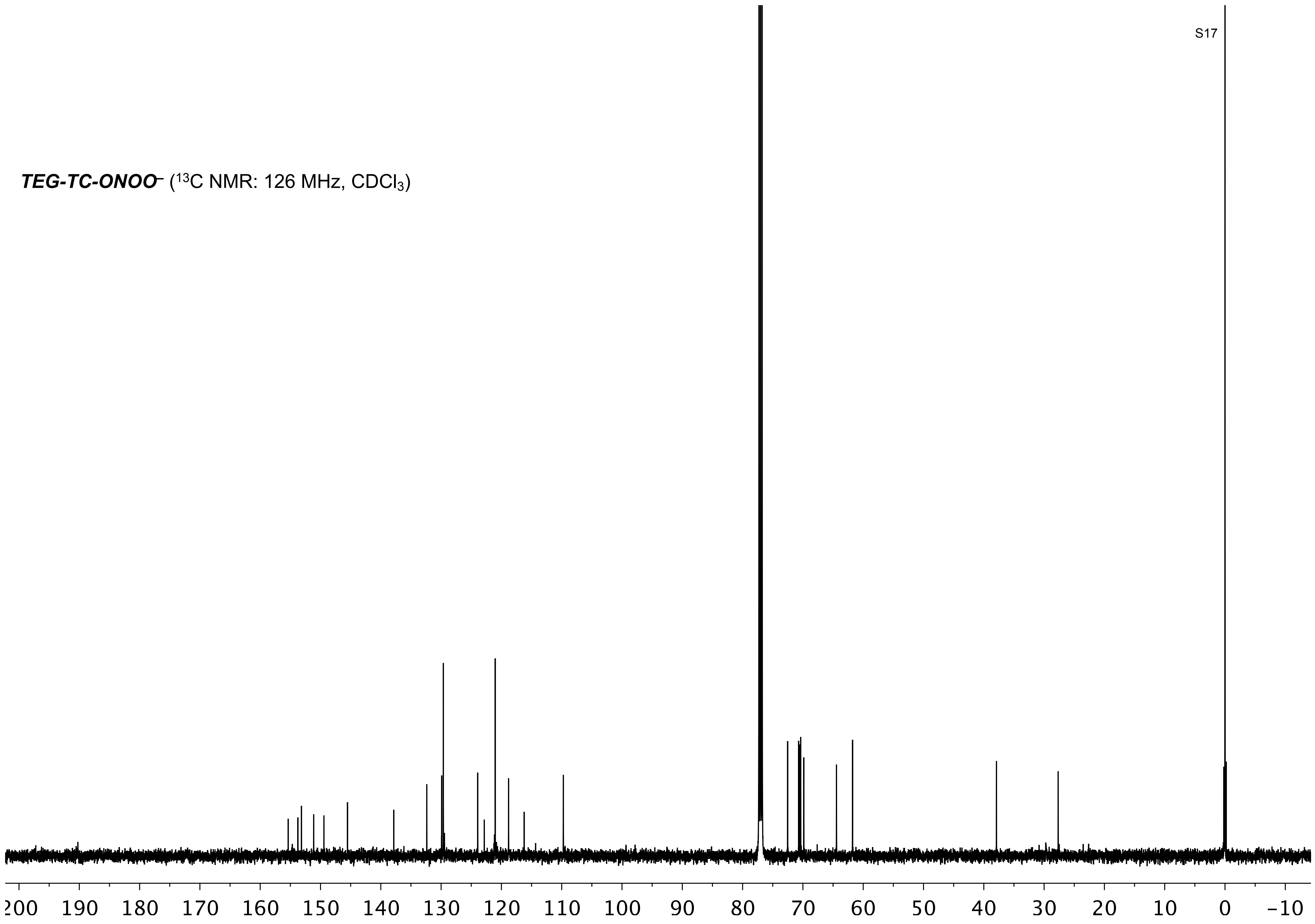

***TEG-TC-ONOO⁻*** ($^{13}$C NMR: 126 MHz, $CDCl_3$)
200 190 180 170 160 150 140 130 120 110 100 90 80 70 60 50 40 30 20 10 0 −10

***TEG-TC-ONOO*⁻** ($^{19}$F NMR: 471 MHz, $CDCl_3$)

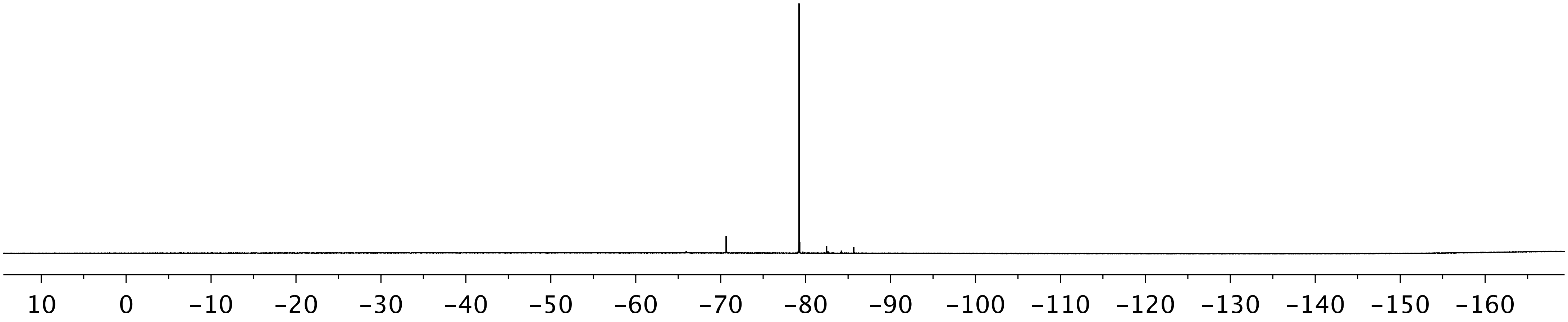

Compound ***S2*** ($^{1}$H NMR: 500 MHz, $CDCl_3$)

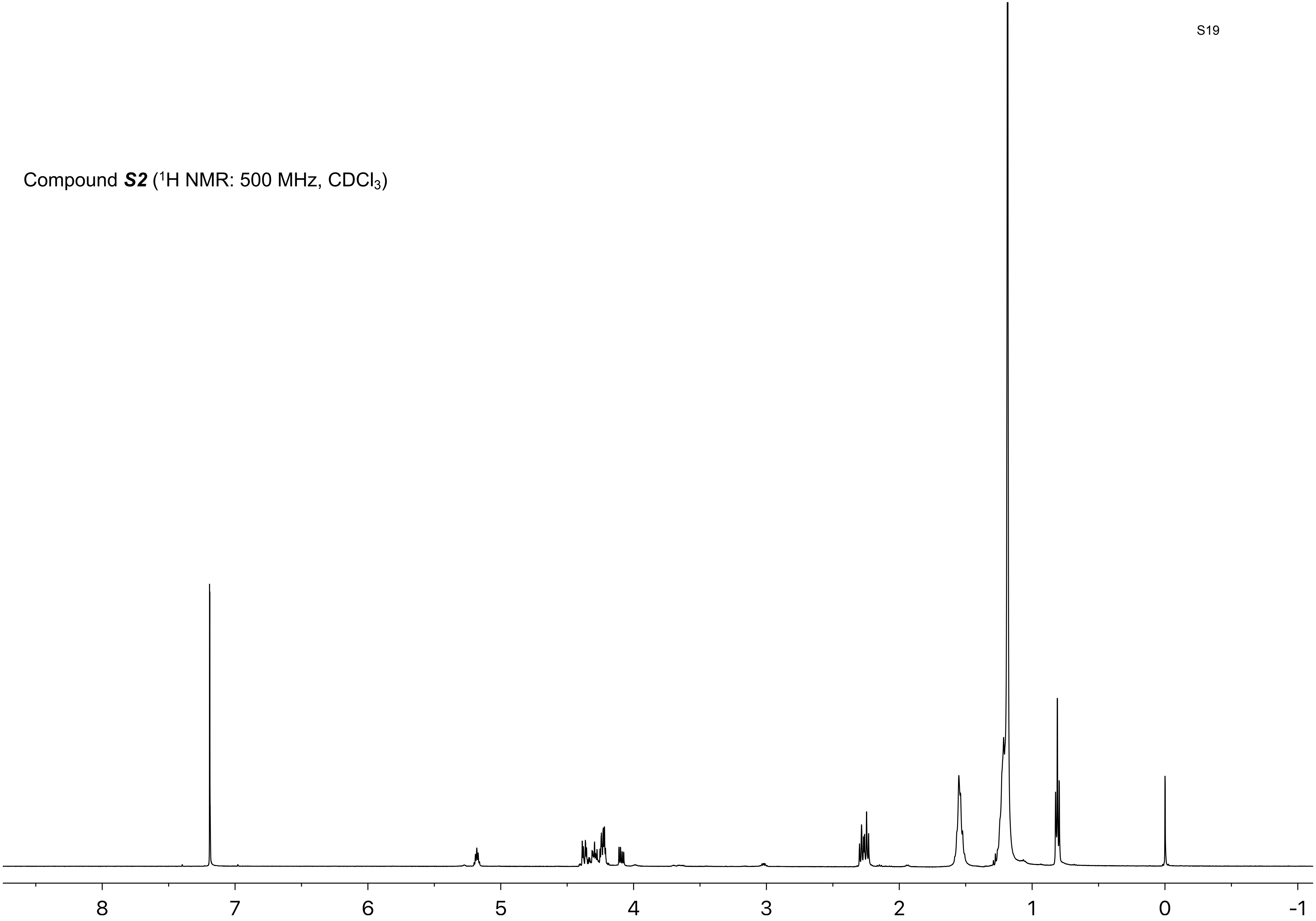

Compound ***S2*** ($^{13}C$ NMR: 126 MHz, $CDCl_3$)

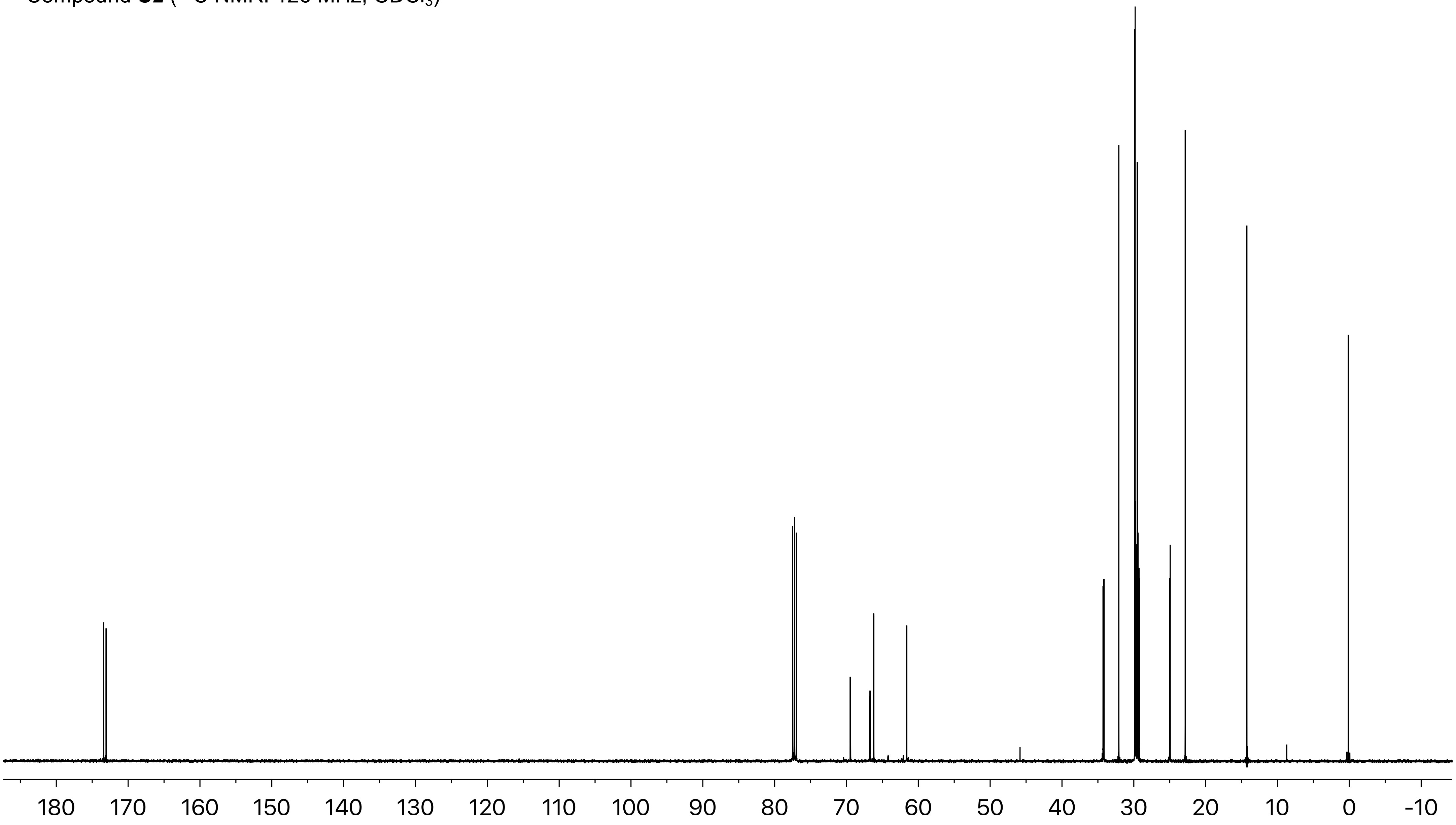

Compound ***S2*** ($^{31}P$ NMR: 202 MHz, $CDCl_3$)

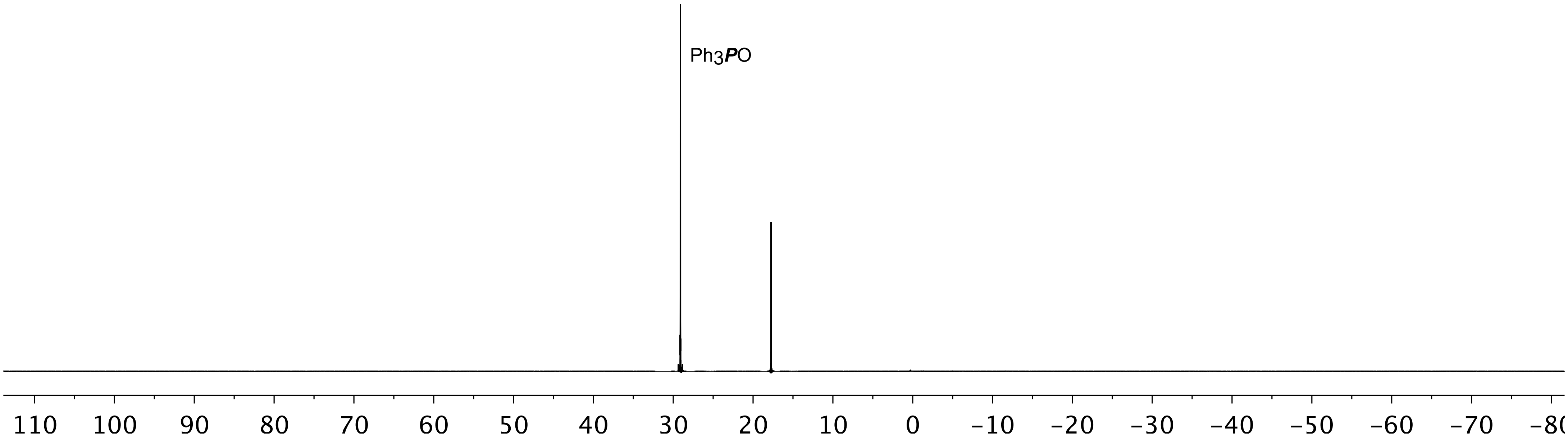

Compound ***4*** ($^1$H NMR: 500 MHz, $CDCl_3$)

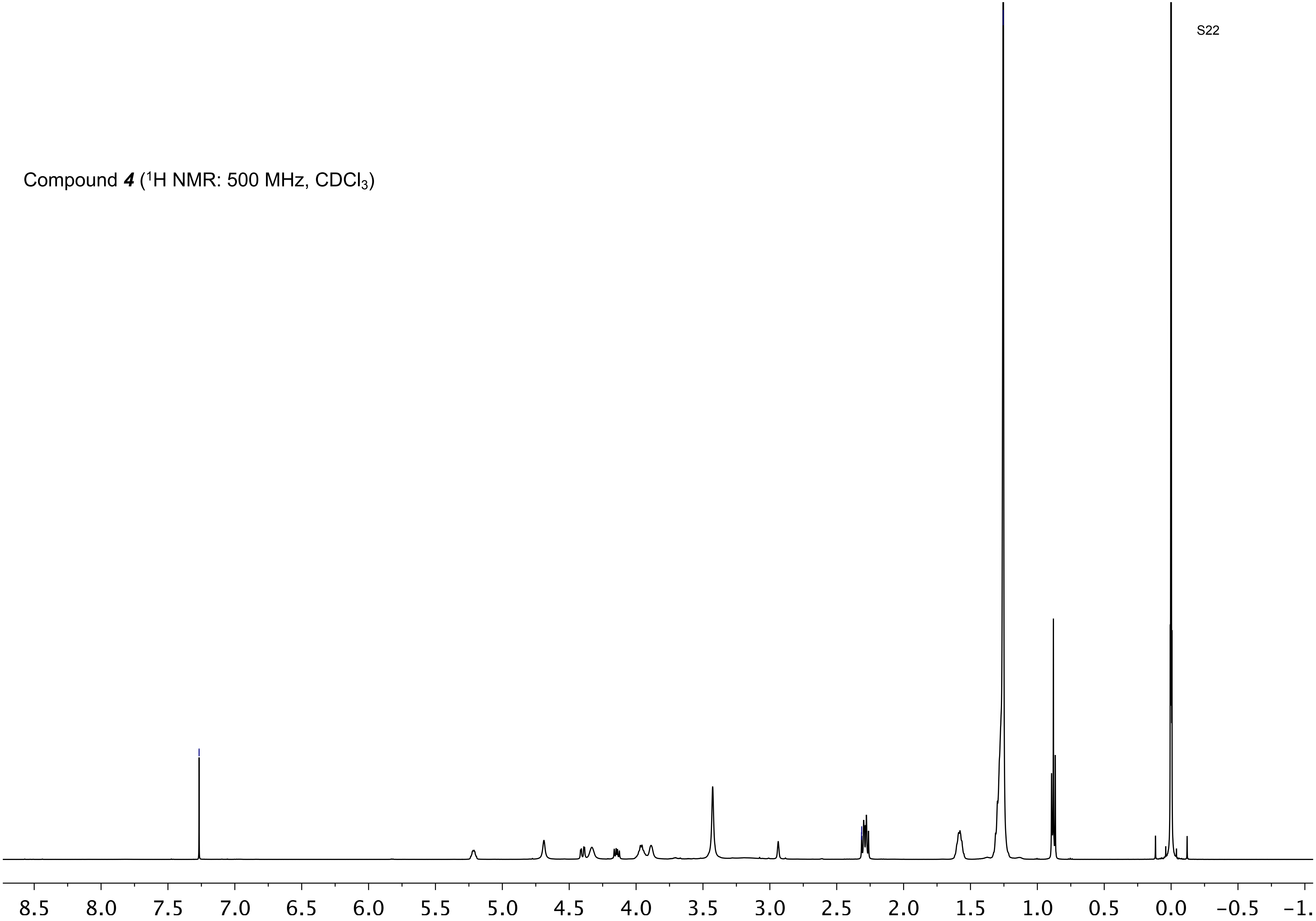

Compound ***4*** ($^{13}C$ NMR: 126 MHz, $CDCl_3$)

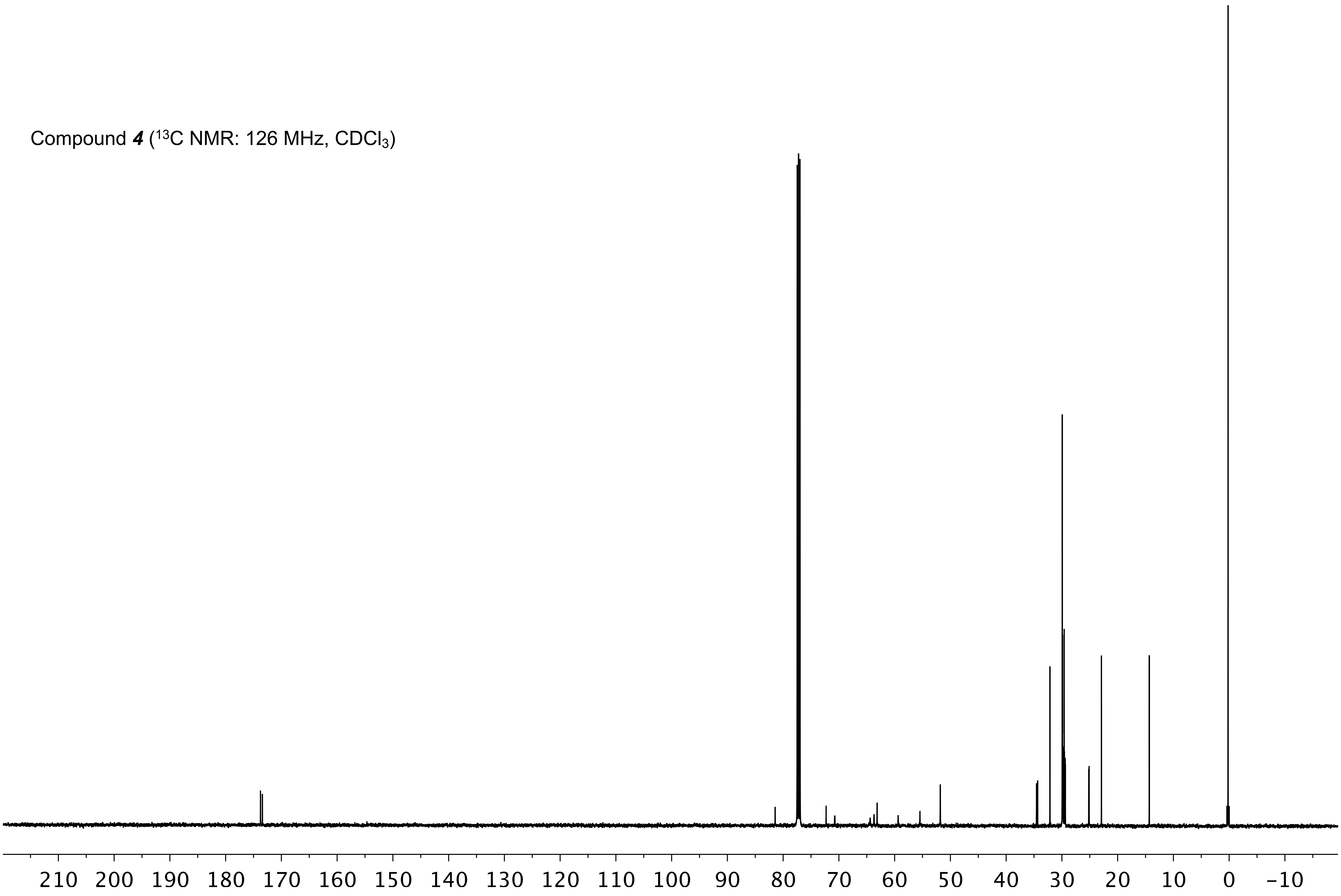

Compound ***4*** ($^{31}$P NMR: 202 MHz, $CDCl_3$)

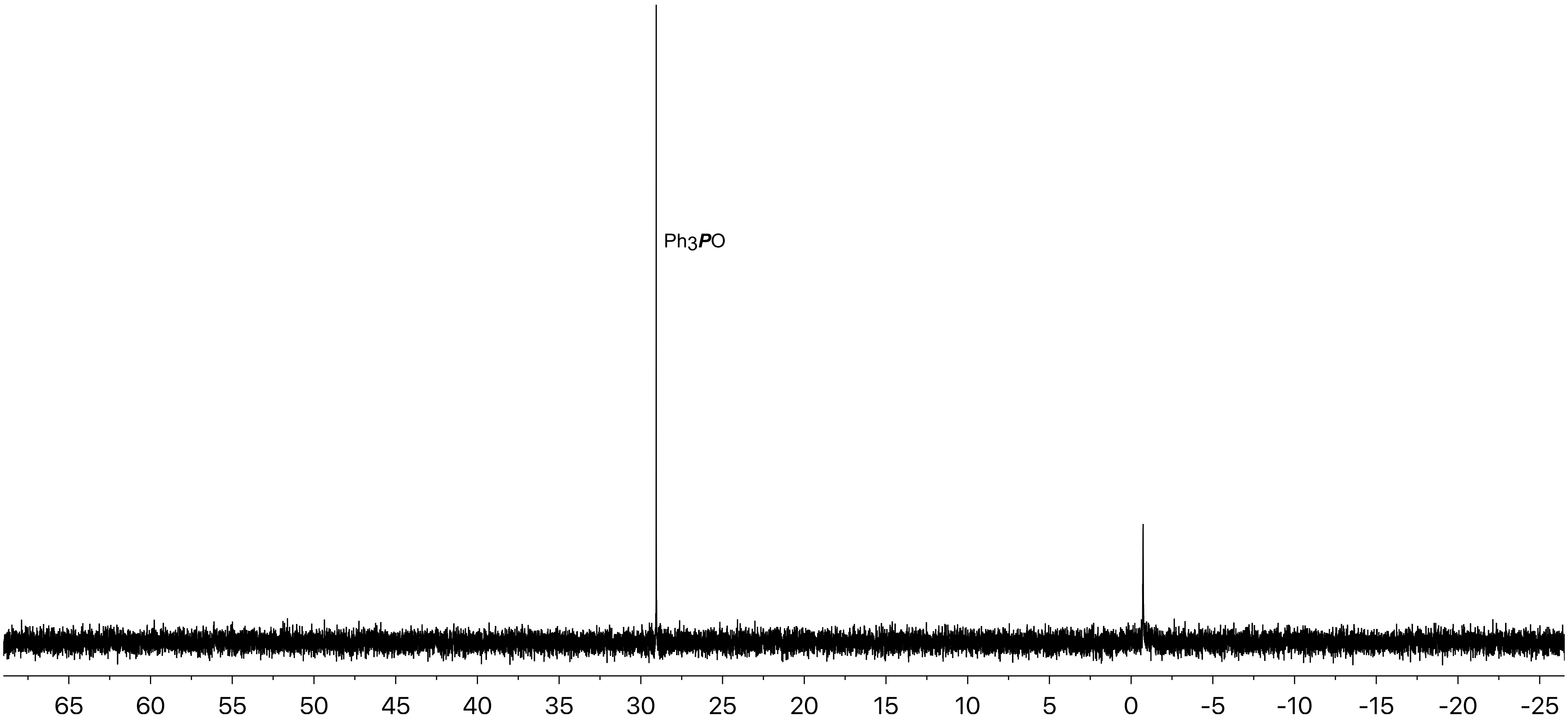

***DPPC-TC-ONOO⁻*** ($^1$H NMR: 500 MHz, $CDCl_3$ / $CD_3OD$, 3:1 v/v)

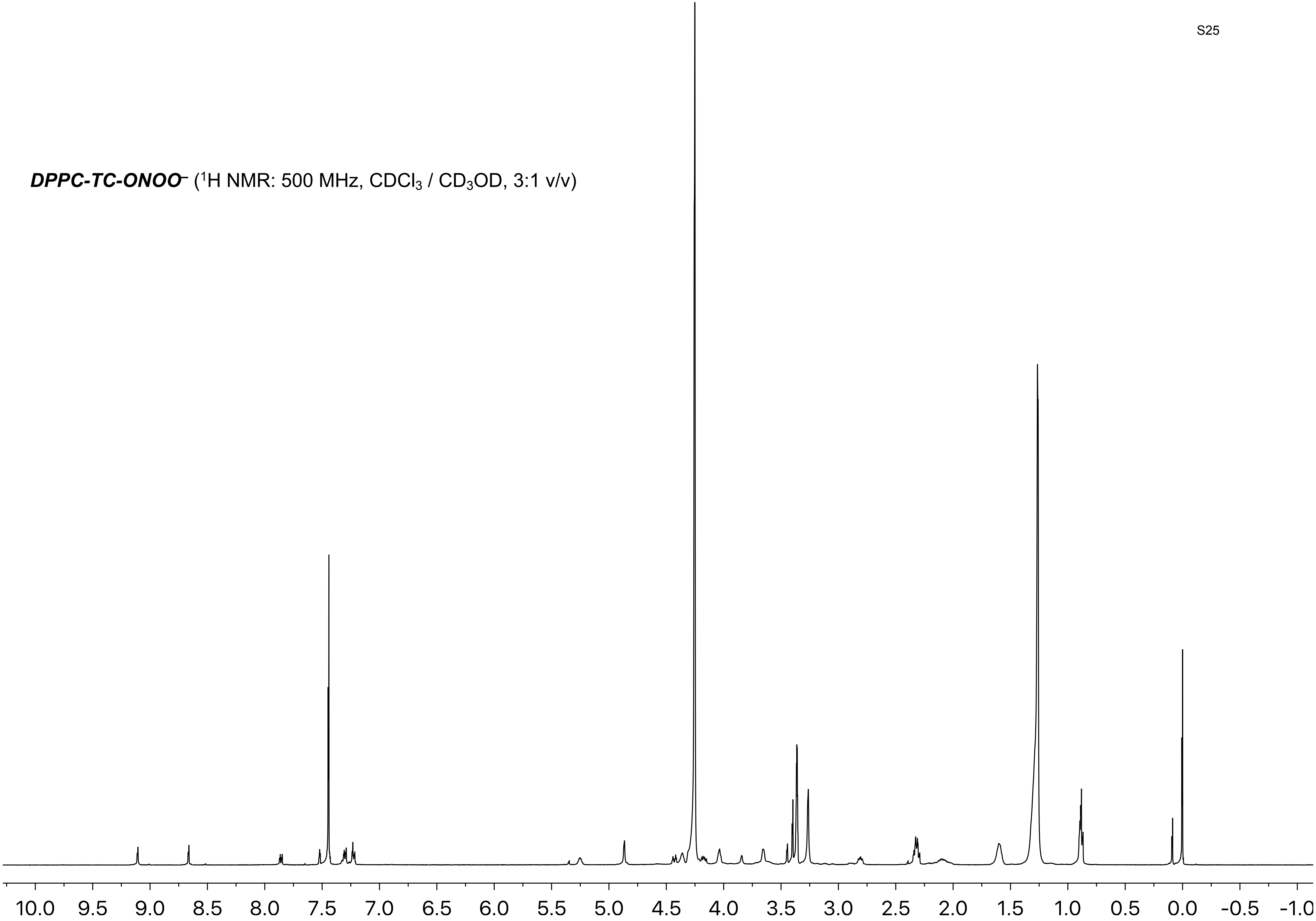

***DPPC-TC-ONOO⁻*** ($^{13}C$ NMR: 126 MHz, $CDCl_3$ / $CD_3OD$, 3:1 v/v)

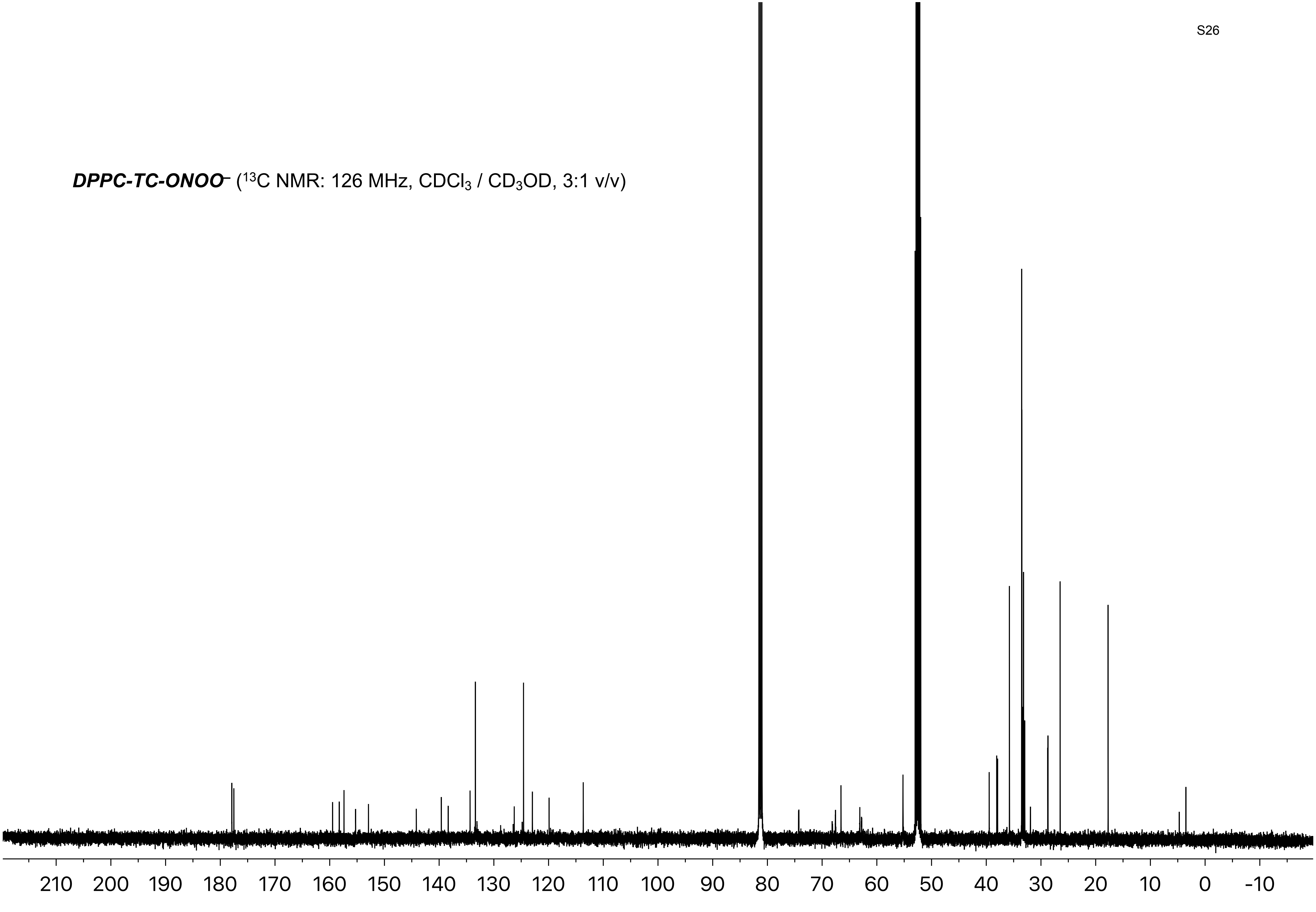

***DPPC-TC-ONOO*** – ($^{31}$P NMR: 202 MHz, $CDCl_3$ / $CD_3OD$, 3:1 v/v)

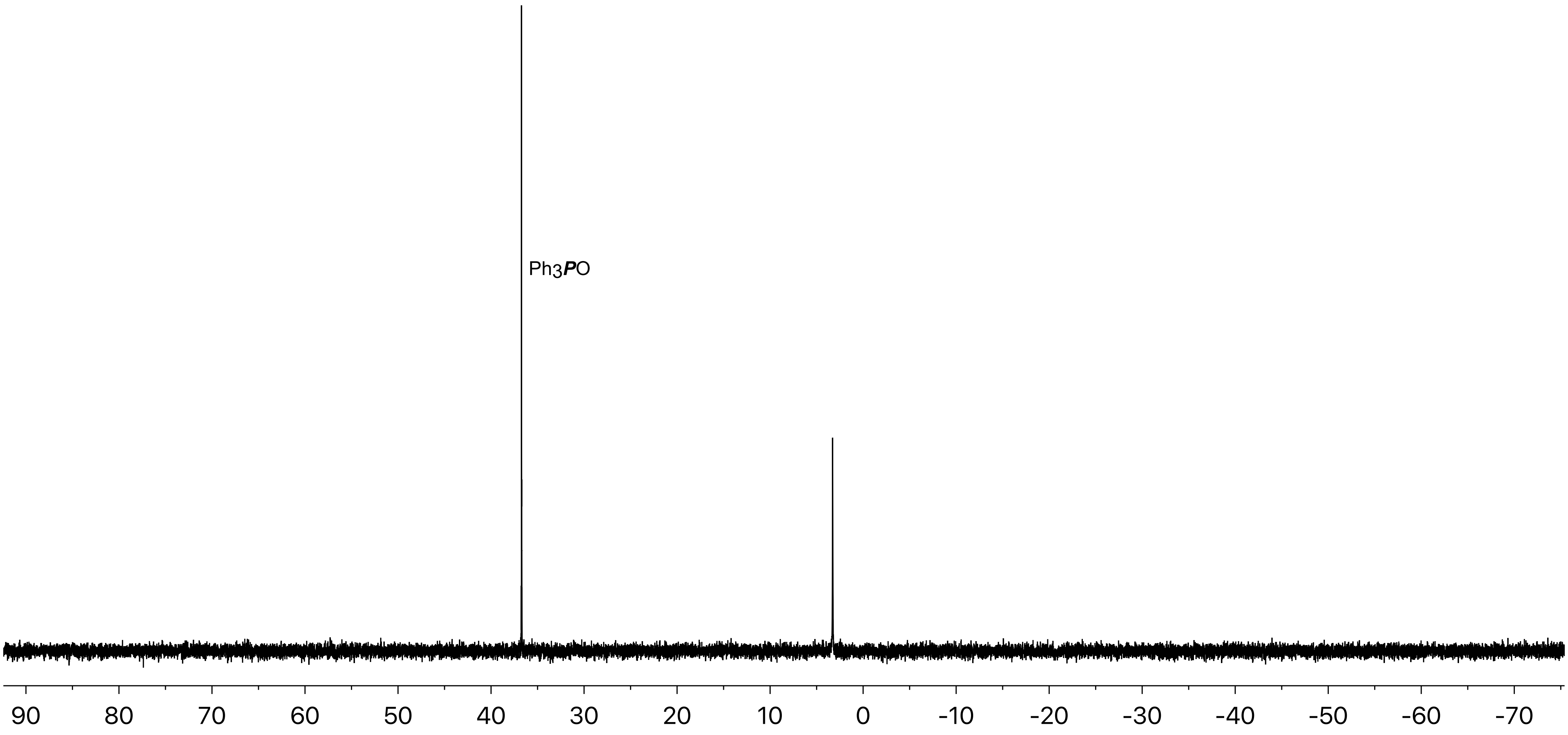

***DPPC-TC-ONOO⁻*** ($^{19}F$ NMR: 471 MHz, $CDCl_3$ / $CD_3OD$, 3:1 v/v)

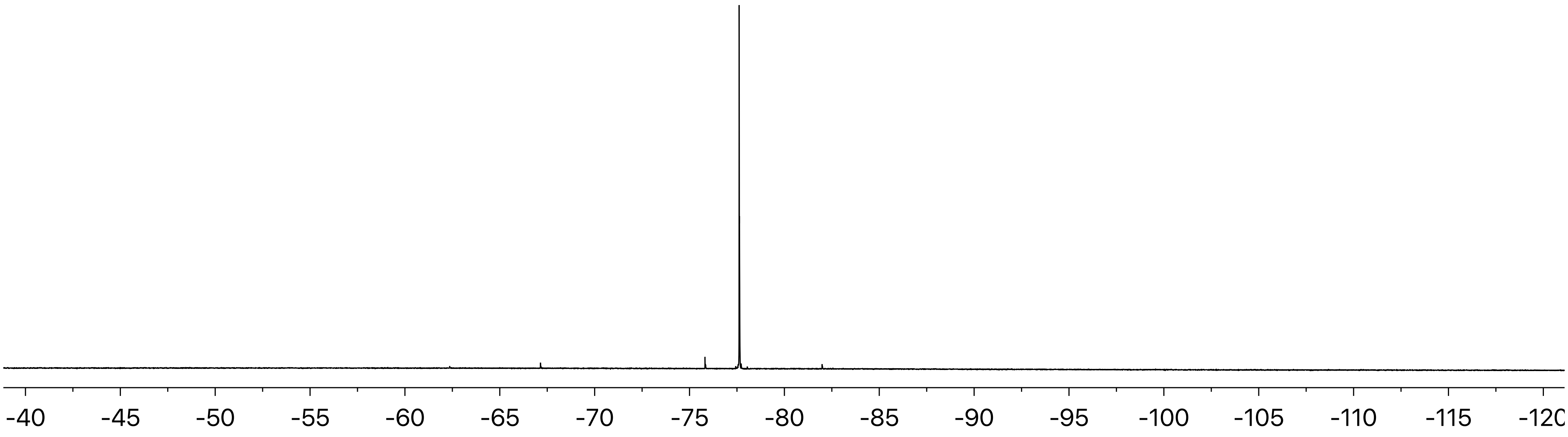

***TEG-TC*** ([1]H NMR: 500 MHz, $CD_3OD$)

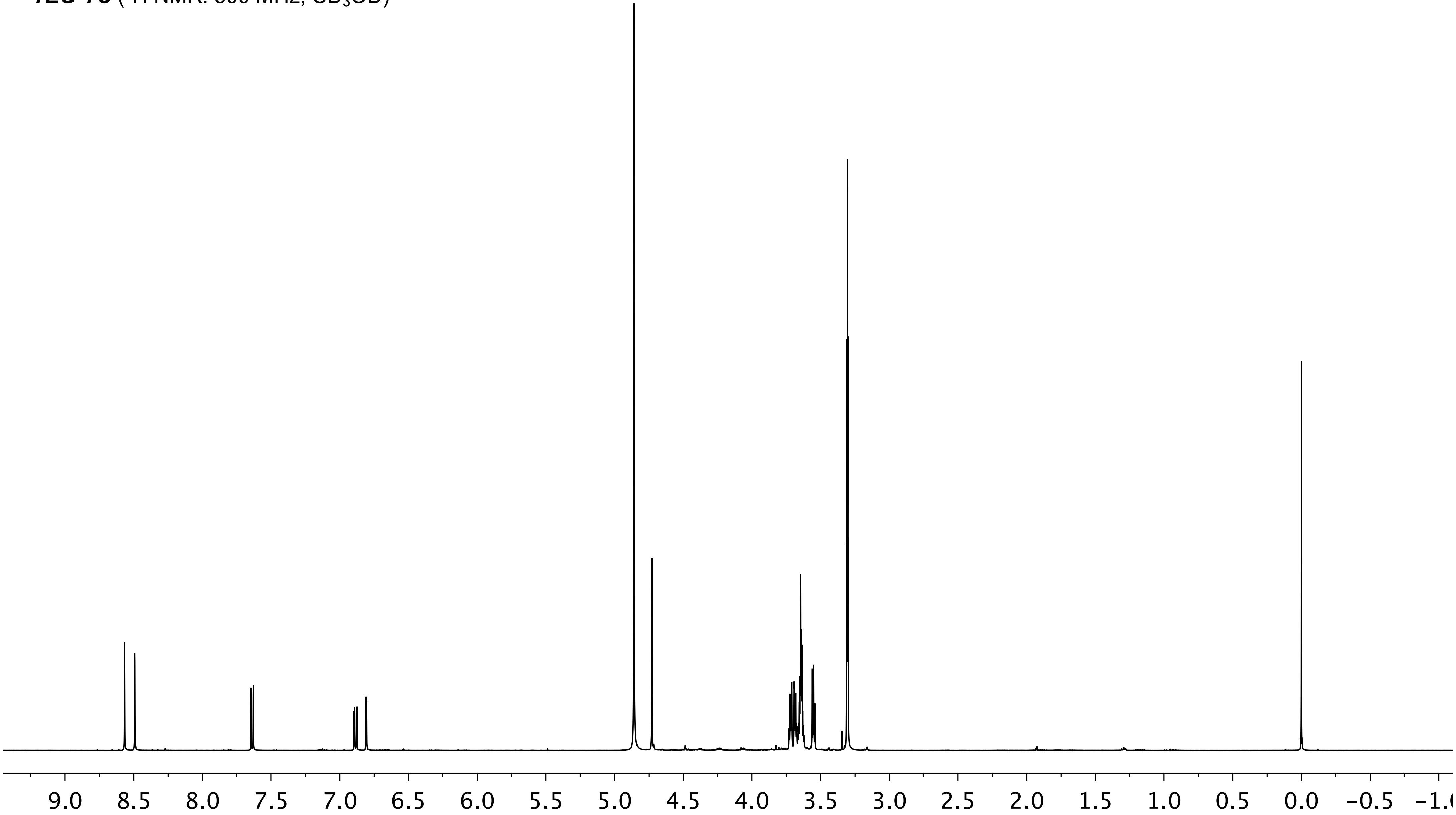

***TEG-TC*** ($^{13}C$ NMR: 126 MHz, $CD_3OD$)

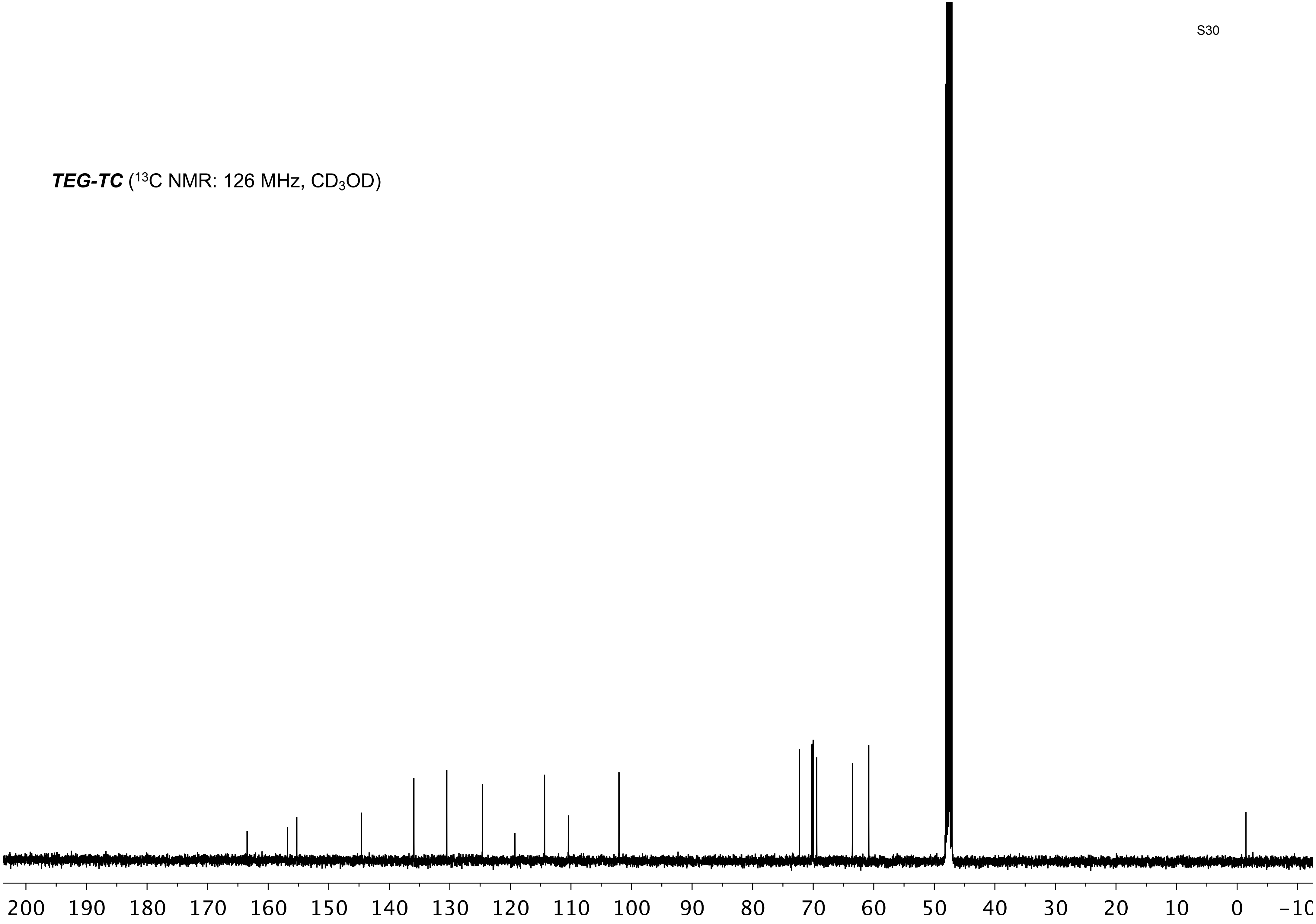

***DPPC-TC*** ($^1$H NMR: 500 MHz, $CDCl_3$ / $CD_3OD$, 3:1 v/v)

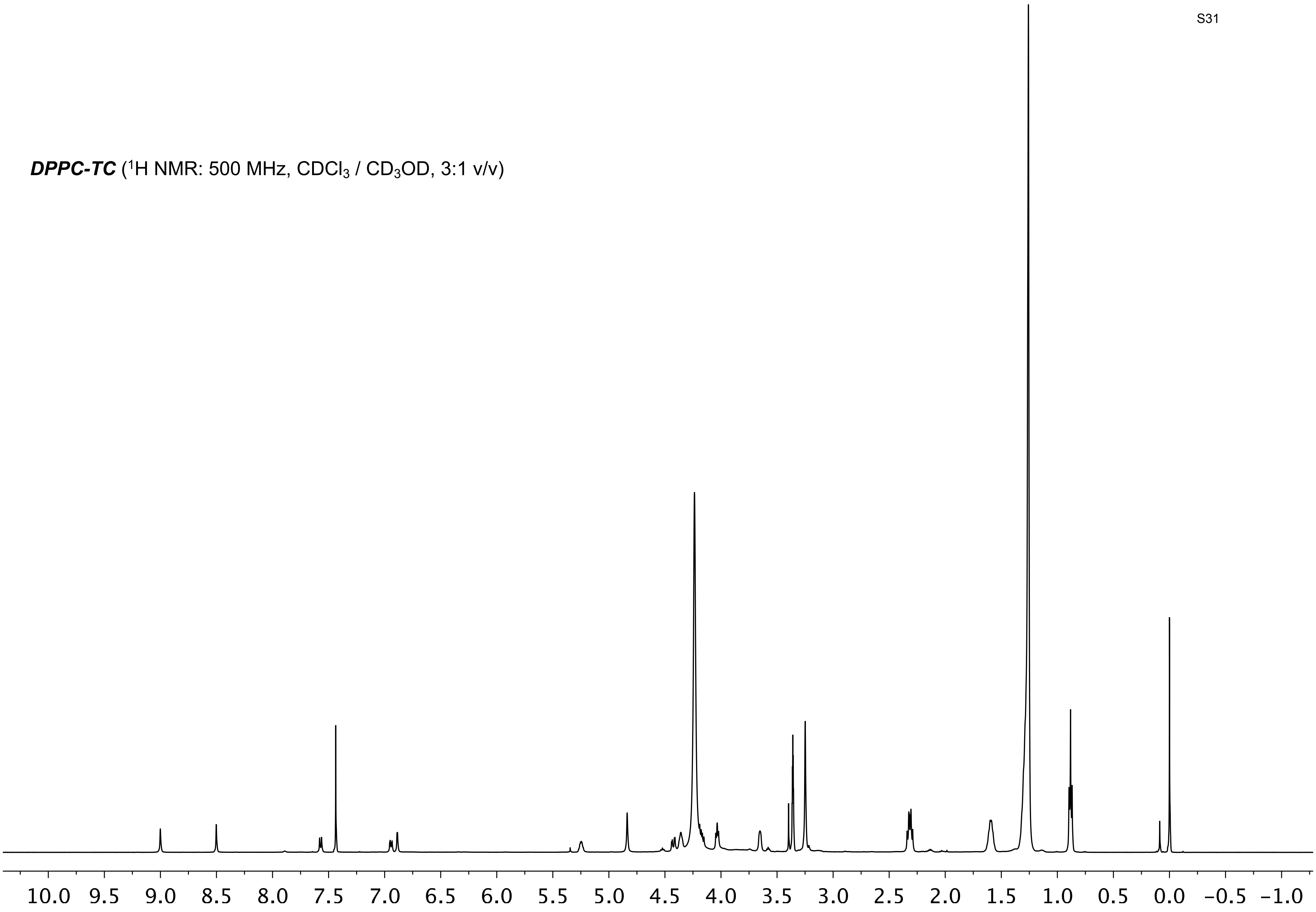

***DPPC-TC*** ($^{13}$C NMR: 126 MHz, $CDCl_3$ / $CD_3OD$, 3:1 v/v)

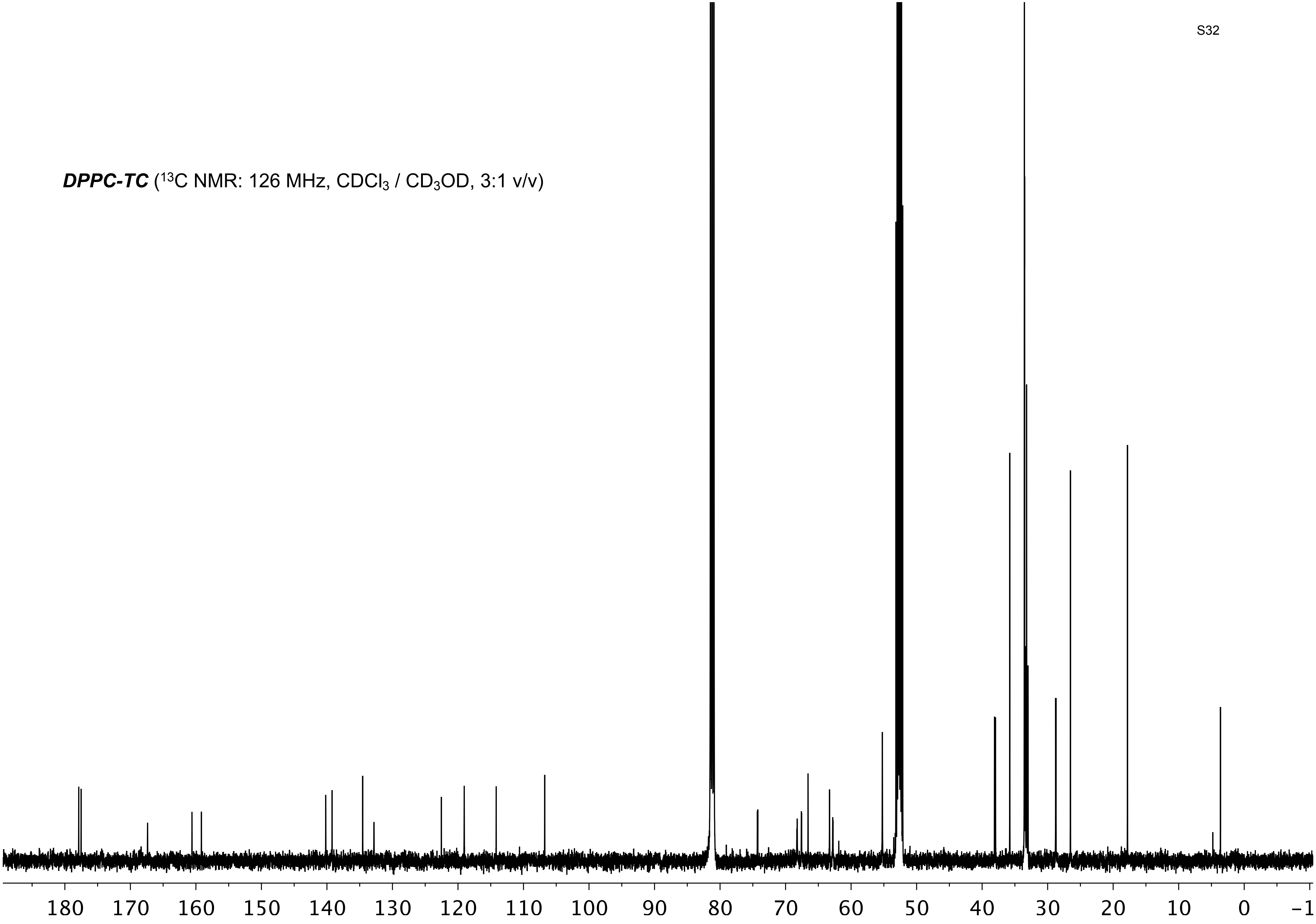

***DPPC-TC*** ($^{31}P$ NMR: 202 MHz, $CDCl_3$ / $CD_3OD$, 3:1 v/v)

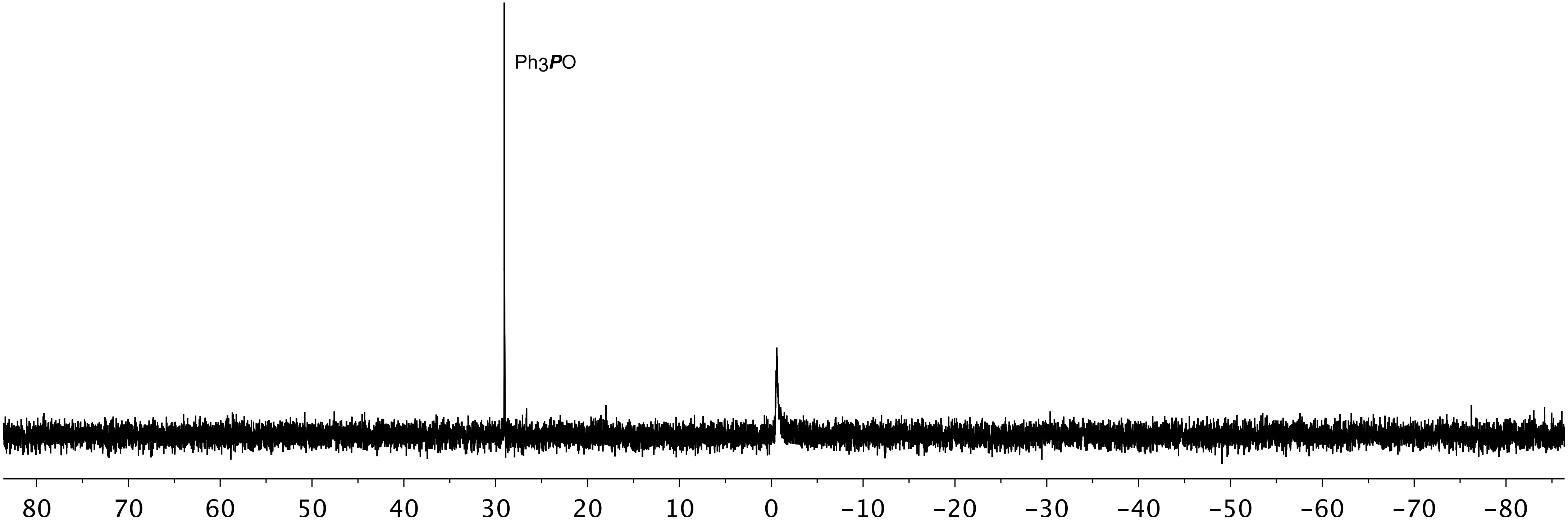

### LCMS Analysis of Reaction of TEG-TC-ONOO⁻ with ONOO⁻

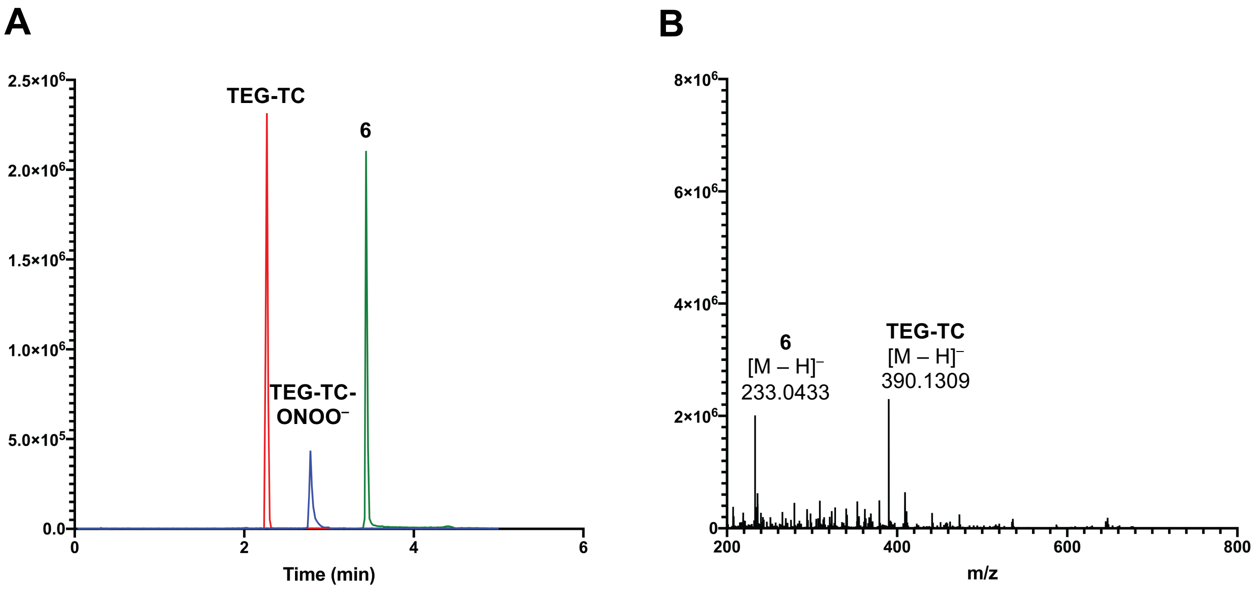


**Supplemental Figure 1. (A)** Overlay of extracted ion chromatograms of compounds **TEG-TC** (red), **TEG-TC-ONOO⁻** (blue), and the oxa-spiro[4,5] decenone **6** (green) as the major products from the reaction (5 minutes) between **TEG-TC-ONOO⁻** (10 µM) and ONOO⁻ (600 µM). Reaction solvent: $H_2O$/MeOH (1:4 v/v). Aqueous part contained Tris (50 mM, pH 7.5). **(B)** HRMS analysis of the reaction aliquot taken in 5 minutes. Spectra are shown for the retention times of 2–4 minute window. HRMS analysis was carried out in negative ionization mode.

**TEG-TC:**

**HRMS** (ESI) m/z: Calculated for $C_{18}H_{20}N_3O_7^-$, [M - H]⁻, requires 390.1307; found 390.1309.

The oxa-spiro[4,5] decenone **6:**

**HRMS** (ESI) m/z: Calculated for $C_{10}H_8F_3O_3^-$, [M – H]⁻, requires 233.0431; found 233.0433.

### Determination of p$K_a$ of TEG-TC

The absorbance of the model coumarin **TEG-TC** (20 μM) at 405 nm was measured in buffer solutions with pH values: 5.2, 6.0, 7.0, 7.5, 8.0, 8.5, 9.0, and 9.5. The buffers at pH 5.2 and 6.0 were obtained using 50 mM MES, while those at pH 7.0–9.5 were obtained using 50 mM Tris. Resulting data was fitted to a sigmoidal curve (Supplemental Figure 2), providing a p$K_a$ of 7.1 ± 0.1.

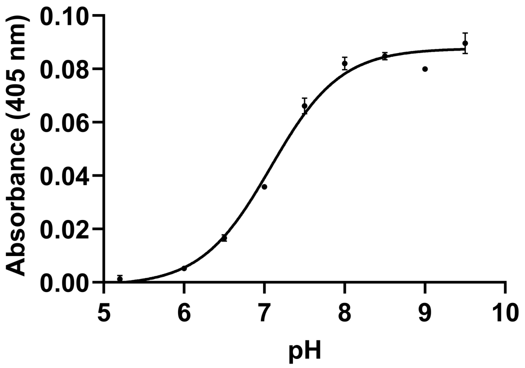


**Supplemental Figure 2.** Absorbance (405 nm) of **TEG-TC** vs. pH. For each absorbance measurement, a 20 μM solution of **TEG-TC** was prepared in buffers with a pH ranging from 5.2–9.5. The measured pH values were used to plot a sigmoidal dose-response curve ($R^2$ = 0.989). Error bars represent standard deviation, $n$ = 3.

### pH dependence of fluorescence intensity for TEG-TC

The fluorescence intensity output of the model coumarin **TEG-TC** (1 μM) at 475 nm was measured by 405 nm excitation wavelength in buffer solutions with pH values: 5.8, 6.8, 7.5, 7.8, and 8.8. The buffer at pH 5.8 was obtained using 50 mM MES, while those at pH 6.8–8.8 were obtained using 50 mM Tris.

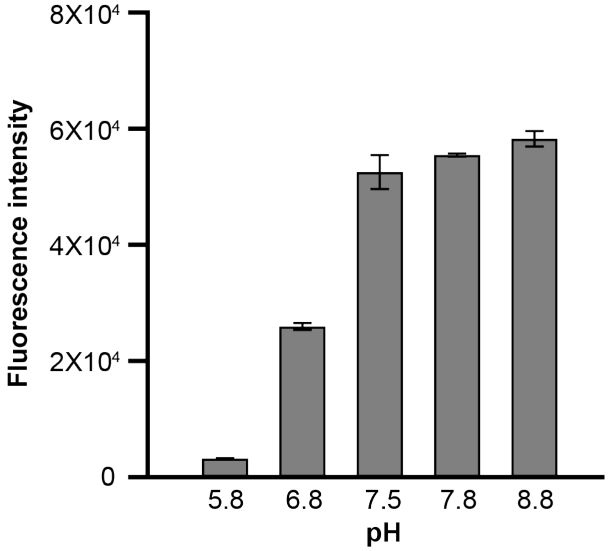


**Supplemental Figure 3.** Fluorescence intensity (405 nm excitation, 475 nm emission) of **TEG-TC** (1 μM) vs. pH. Error bars represent standard deviation, *n* = 3.

### Determination of Relative Fluorescence Quantum Yield

The relative fluorescence quantum yields for **TEG-TC-ONOO⁻**, **TEG-TC**, **DPPC-TC-ONOO⁻**, and **DPPC-TC** were determined using a previously reported equation (1) shown below:

$$Q_s = Q_r\,(m_s/m_r)(n_s/n_r)^2$$

where $Q_s$ is the quantum yield of the unknown, $Q_r$ is the quantum yield of a known dye, $m_s$ and $m_r$ are gradient of the plot of integrated fluorescence intensity against absorbance of the unknown and known dyes, respectively, and lastly, n (s and r) is the index of refraction. Absorbance (405 nm) and fluorescence intensity ($\lambda_{ex/em}$ of 405/475 nm) were measured. Coumarin 343 was used as the reference standard for $Q_r$ (0.63) in ethanol which was found directly from Sigma Aldrich. Ethanol was used for both the standard and the samples; the refractive index ratio is equal to 1. The quantum yields for **TEG-TC-ONOO⁻**, **TEG-TC**, **DPPC-TC-ONOO⁻**, and **DPPC-TC** were 0.08, 0.66, 0.09, and 0.64, respectively.

### Microplate Fluorescence Measurements

Stock solutions of the redox-active species were prepared at 18 mM concentration. 1.5 μL of **TEG-TC-ONOO⁻** in DMSO (100 μM) and 5 μL of the stock solution of the redox-active species were added to a 96-well plate containing Tris buffer (50 mM, pH 7.5), for a total volume of 150 μL. At this final volume, the concentrations of **TEG-TC-ONOO⁻** and the redox-active species were predicted to be 1 μM and 600 μM, respectively. Fluorescence intensity ($\lambda_{ex/em}$ 405/475 nm) of the samples were measured on a microplate reader (Molecular Devices SpectraMax iD3 or Tecan Infinite Pro 2000) at the 1 and 60-minute marks. For peroxynitrite, additional measurements were taken at 3, 5, 10, 20, 30, 40, and 50 minutes. This protocol was also employed for **TEG-TC** (positive control). $F/F_0$ was calculated by taking the final fluorescent measurements at the 60-minute mark over the initial fluorescence measurements at $t_0$.

### Spectrophotometric Characterizations by Absorbance and Emission

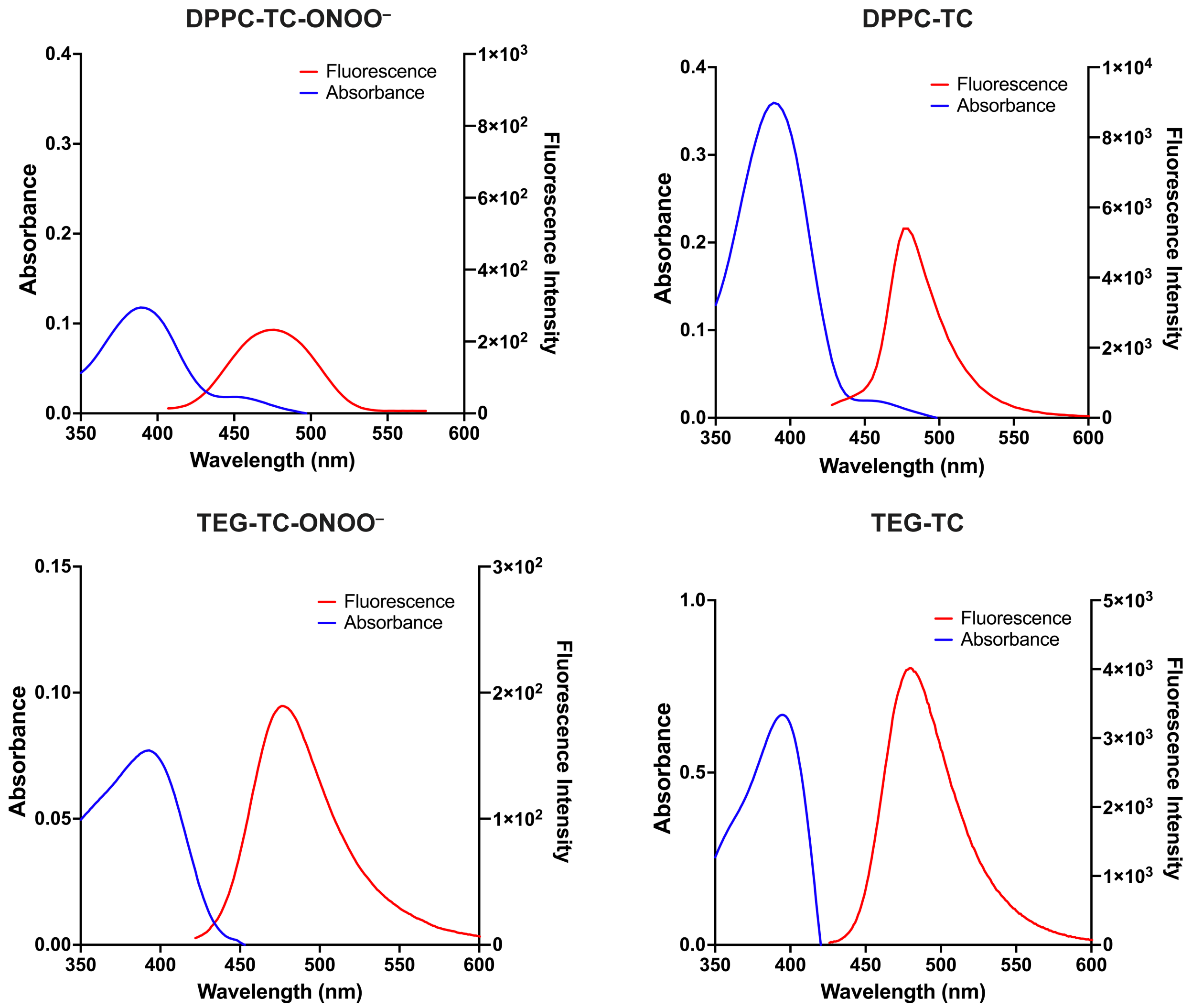


**Supplemental Figure 4.** Absorbance and emission profiles of the probes and their uncaged analogs.

### UV-Vis Spectrophotometric Investigation of Peroxynitrite

Two sets of data were acquired based on $CO_2$ treatment: (1) $CO_2$-rich and (2) $CO_2$-deprived. (1) For the $CO_2$-rich condition, the peroxynitrite-free aqueous buffer was first bubbled with $CO_2$ gas from dry ice and the pH was adjusted to 7.5 by careful addition of NaOH. (2) For the $CO_2$-deprived condition, the peroxynitrite-free aqueous buffer was first boiled to eliminate $CO_2$, and the pH was adjusted to 7.5 by careful addition of HCl. A freshly prepared $ONOO^-$ solution (2.2 mM, pH 10.6) was added to each buffer and the absorbance at 302 nm was recorded over 10 minutes. Based on dilution, the concentration of $ONOO^-$ at the moment of mixing was estimated to be 100 μM.

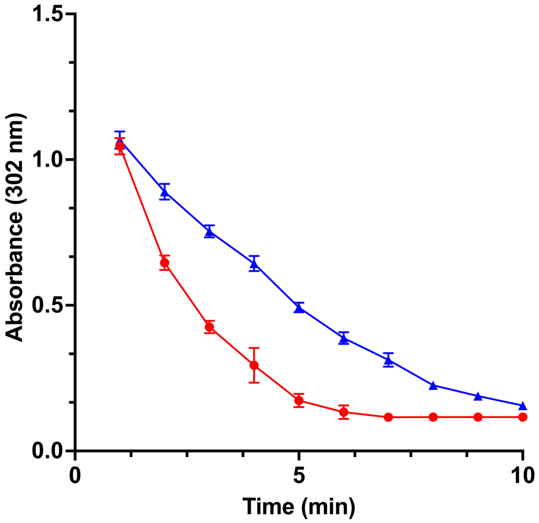


**Supplemental Figure 5.** Absorbance (302 nm) of peroxynitrite versus time at pH 7.5 for $CO_2$-rich (red circle) and $CO_2$-deprived (blue triangle). Error bars represent standard deviation, $n$ = 3.

### Effect of Peroxynitrite on TEG-TC-ONOO⁻

UV-Vis spectrophotometric measurements were carried out to assess the conversion of the model probe **TEG-TC-ONOO⁻** to the uncaged coumarin **TEG-TC** using peroxynitrite. Absorbance was measured with 3 nm intervals at 0, 5, 30, and 60 minutes.

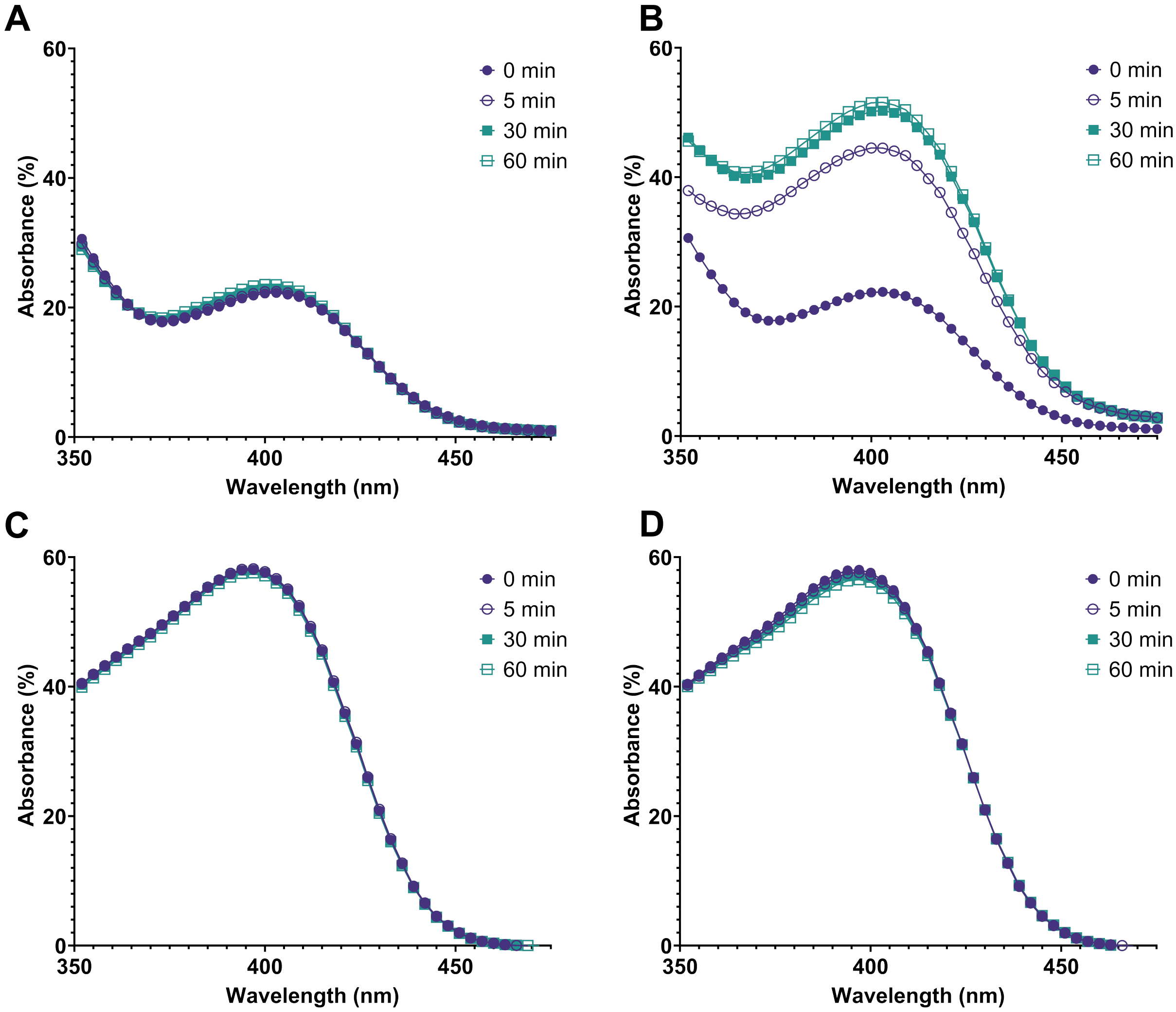


**Supplemental Figure 6.** Time course for the UV-Vis absorbance of the aqueous mixtures including **(A) TEG-TC-ONOO⁻** (200 μM), **(B) TEG-TC-ONOO⁻** (200 μM) and peroxynitrite (600 μM), **(C) TEG-TC** (200 μM), and **(D) TEG-TC** (200 μM) and peroxynitrite (600 μM). Percent conversion of the probe **TEG-TC-ONOO⁻** into the coumarin **TEG-TC** by peroxynitrite was 67% at 5 minutes and 88% at 60 minutes. All mixtures were prepared in Tris buffer (150 mM, pH 7.5). Data acquired at room temperature. Blank included Tris buffer and DMSO. Probe **TEG-TC-ONOO⁻** and the coumarin **TEG-TC** were initially dissolved in DMSO, then introduced into the buffer.

### Preparation and DLS Analysis of Giant Vesicles

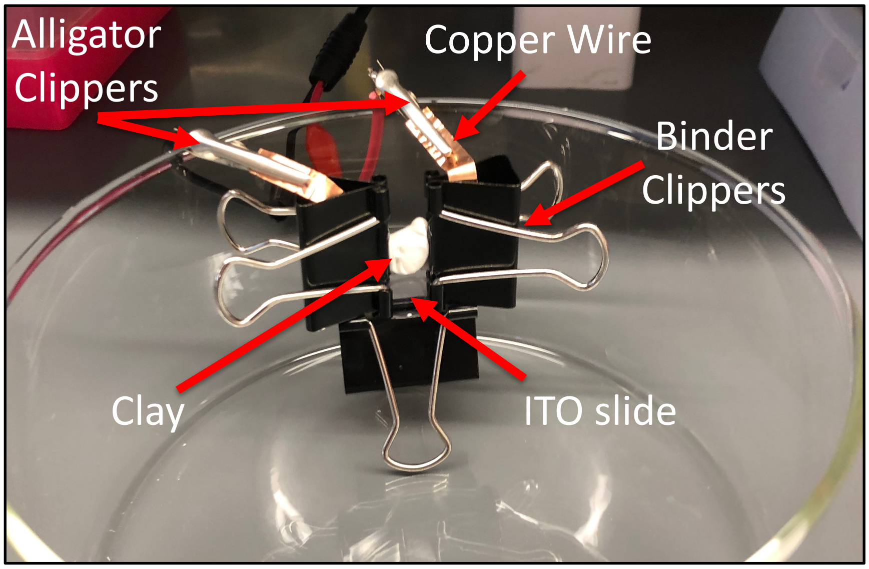


**Supplemental Figure 7.** The electroformation chamber used to form GVs.

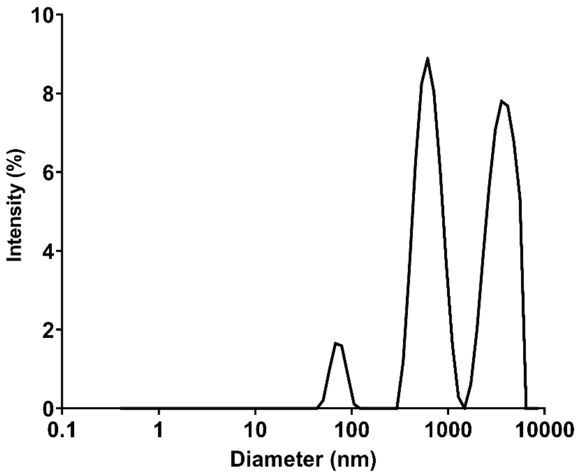


**Supplemental Figure 8.** DLS analysis of the vesicles prepared via electroformation. Approximately 48% of the vesicles had an average size of 5 μm in diameter.

### Staining Cells with Organelle Trackers or Actin Dye

ER-Tracker™ Green (1 μM, 200 μL, 510/540 nm excitation/emission, ER stain) applied at 37 ºC for 10 min; MitoTracker Deep Red FM (250 nM, 200 μL, 644/665 nm, Mitochondrion stain) applied at room temperature for 5 min; CellLight™ Golgi-RFP, BacMam 2.0 (15 μL in Opti-MEM media, 555/584 nm, Golgi stain) applied at 37 ºC, overnight; LysoTracker™ Deep Red (100 nM, 200 μL, 647/670 nm, lysosome stain) applied at 37 ºC for 60 min; CellMask Deep Red Actin dye (1 μM, 200 μL, 669/710 nm, actin stain) applied at 37 ºC for 10 min. After the stated incubation time, the dye solution was removed, cells were washed 3 times with PBS, and 1 mL of HBSS was added prior to imaging.

### MTT Assay for HeLa and RAW 264.7

Cytotoxicity of the probe **DPPC-TC-ONOO⁻**, its uncaged analog **DPPC-TC**, the oxa-spiro[4,5] decenone **6**, and LNPs against HeLa and RAW 264.7 cells was determined using a standard MTT assay. **DPPC-TC-ONOO⁻** and **DPPC-TC** were each mixed with DOPE and DOTMA to form LNPs as described below. HeLa (10,000 cells/well) or RAW 264.7 (25,000 cells/well) were seeded in a 96-well tissue culture plate and cultured overnight to allow the attachment of the cells to the surface. The cell media was then replaced with 100 µL DMEM with 10% FBS containing the liposome or compound (**6** or tamoxifen) of interest. After 24 hours, 10 µL of the 3-(4,5-dimethylthiazol-2-yl)-2,5-diphenyl tetrazolium bromide (MTT reagent) was added to each well and the plate was incubated at 37 °C for 3 hours to yield formazan crystals. To these wells, 100 µL of "detergent reagent" (from the MTT assay kit) was added and the plate was incubated in dark at rt for 2 hours to allow the dissolution of the formazan crystals. The absorbance of each well was recorded at 570 nm on a plate reader. The $OD_{570}$ value for the sample containing cell-free media (DMEM with 10% FBS) was subtracted from each reading. The $OD_{570}$ values were then normalized to the value for the sample containing untreated cells. Using these values, we plotted the dose-response curves of HeLa (Supplemental Figure 8A) or RAW 264.7 cells (Supplemental Figure 8B), which provided $IC_{50}$ values (Supplemental Table 1) through a non-linear regression fitting model.

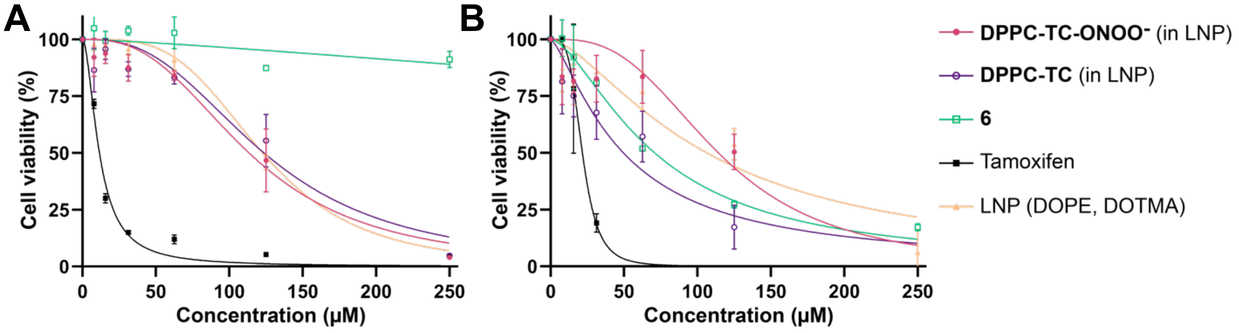


**Supplemental Figure 9.** MTT assay for **(A)** HeLa and **(B)** RAW 264.7 cells after 24 hours of treatment. For each $OD_{570}$ value, the % cell viability vs. concentration plots were obtained using GraphPad Prism 8 software. Error bars represent standard error of mean, $n$ = 3.

**Supplemental Table 1.** Summary of $IC_{50}$ for HeLa (rows 1-2) and RAW 264.7 (rows 3-4).

| | $IC_{50}$ (µM) with 95% CI | | | | |
|---|---|---|---|---|---|
| | **DPPC-TC-ONOO⁻** | **DPPC-TC** | **6** | **Tamoxifen** | **LNP (DOPE, DOTMA)** |
| **HeLa** | 113 ± 14 | 124 ± 18 | >250[a] | 12 ± 1 | 123 ± 6 |
| **RAW 264.7** | 114 ± 28 | 49 ± 13 | 71 ± 8 | 22 ± 2 | 106 ± 28 |

[a] Compound displayed no detectable level of cytotoxicity at 250 µM, which constitutes for the upper bound $IC_{50}$ value. CI: Confidence interval.

### Confocal Images of Live HeLa Cells with DPPC-TC or DPPC-TC-ONOO⁻

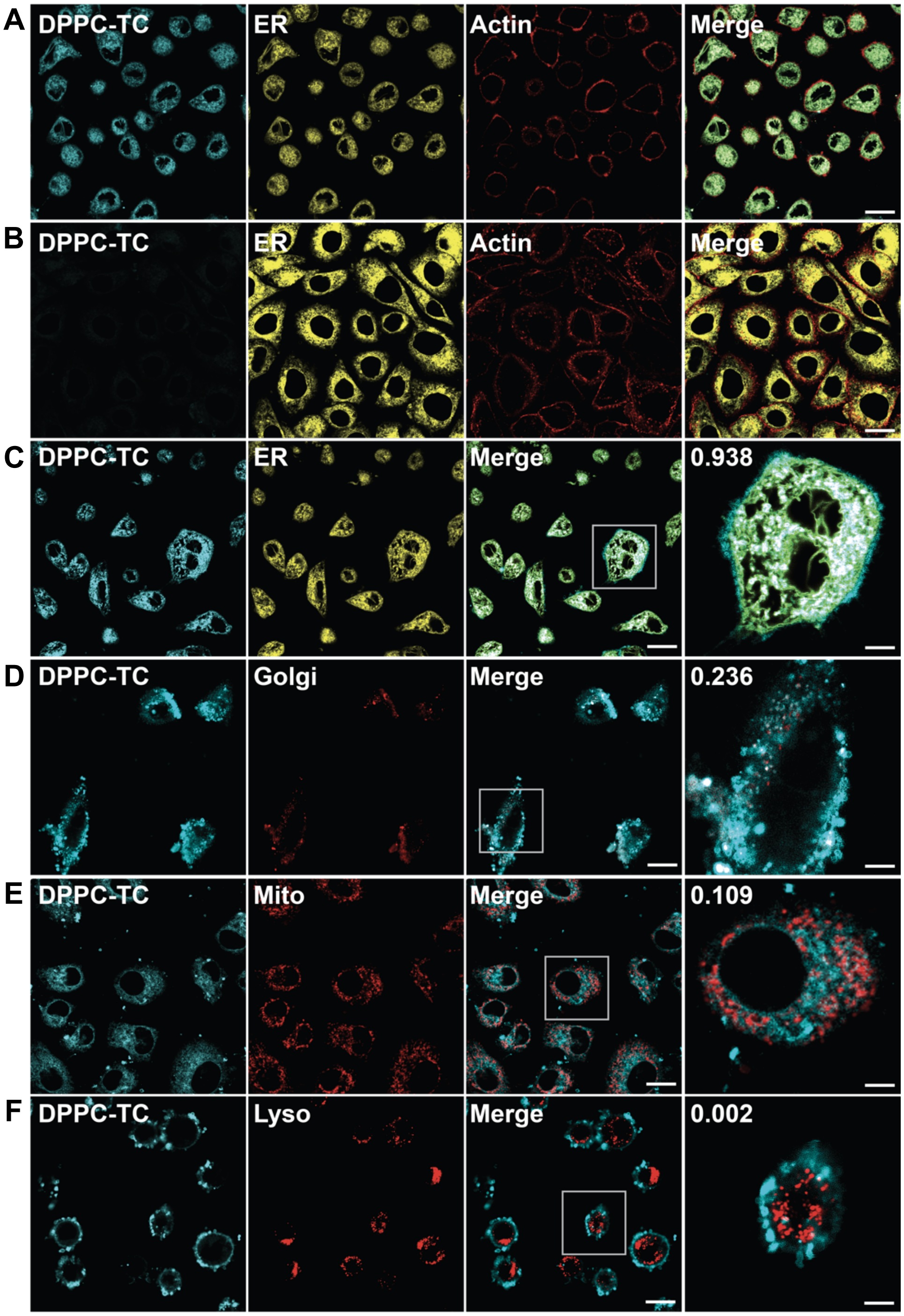


**Supplemental Figure 10. Confocal imaging of lipid environments targeted by ONOO⁻ in live HeLa.** LNPs obtained from a 47.5 : 47.5 : 5.0 molar ratio of DOTMA, DOPE, and **(A) DPPC-TC** or **(B–F) DPPC-TC-ONOO⁻**. **(A-B)** Cells treated with LNPs, left unstimulated. **(C–F)** Quantitative colocalization study of cells treated with LNPs and stimulated with IFN-γ/LPS/PMA. PCC was calculated for assessment of colocalization. DPPC-TC channel: 405/475 nm. Scale bars = 20 μm or 5 μm (zoomed area).

**Confocal Images of Live RAW 264.7 Cells with DPPC-TC or DPPC-TC-ONOO⁻**

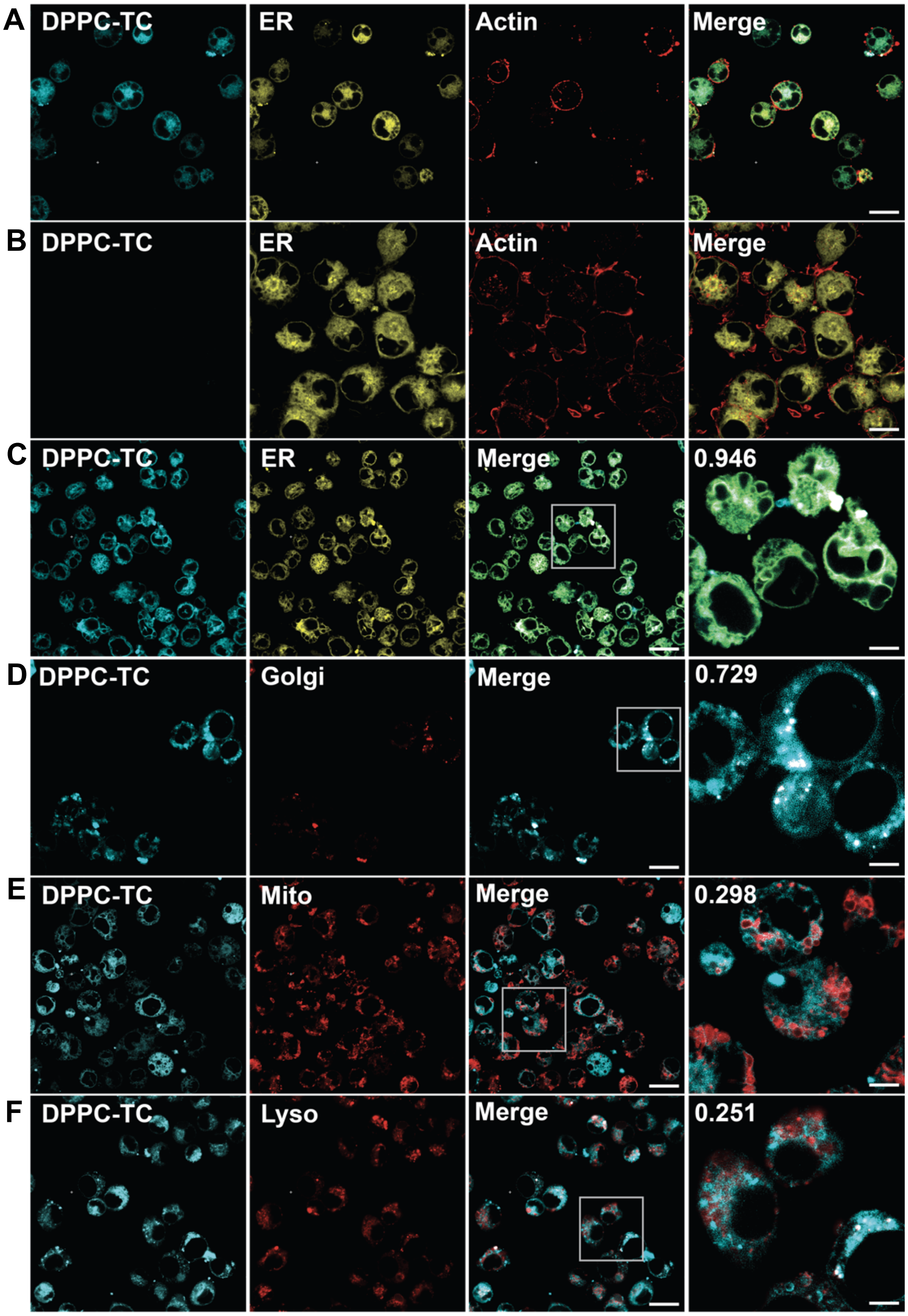


**Supplemental Figure 11. Confocal imaging of lipid environments targeted by ONOO⁻ in live RAW 264.7 cells.** LNPs obtained from a 47.5 : 47.5 : 5.0 molar ratio of DOTMA, DOPE, and **(A) DPPC-TC** or **(B–F) DPPC-TC-ONOO⁻**. **(A-B)** Cells treated with LNPs, left unstimulated. **(C–F)** Quantitative colocalization study of cells treated with LNPs and stimulated with LPS. PCC was calculated for assessment of colocalization. DPPC-TC channel: 405/475 nm. Scale bars = 20 µm or 5 µm (zoomed area).

### Confocal Images of Live Cells using LNPs with Liss-Rhod PE

Investigation of Liss-Rhod PE localization in both HeLa and RAW 264.7 cells was carried out using the protocol described in Materials and Methods, under the section “Protocols for cell studies”.

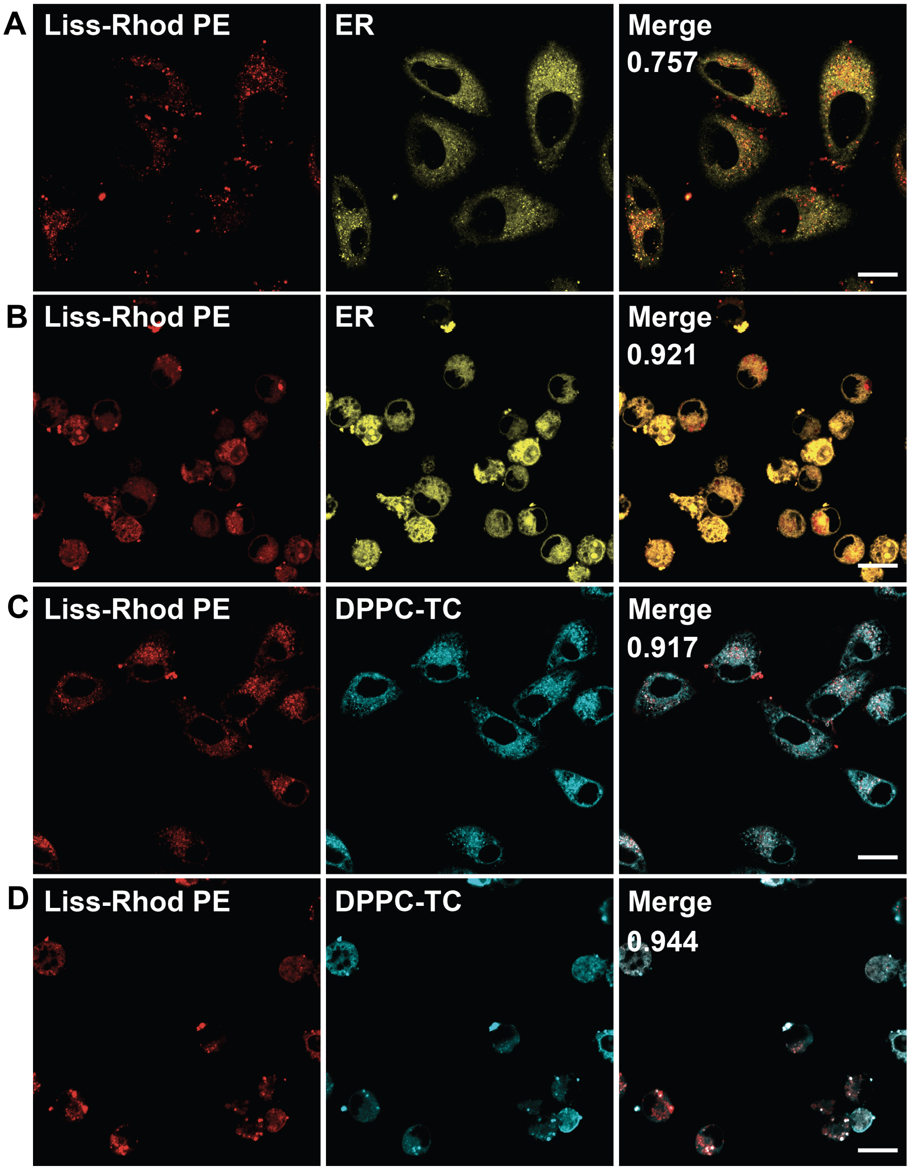


**Supplemental Figure 12.** Confocal images of HeLa **(A, C)** and RAW 246.7 **(B, D)** cells incubated with LNPs containing Liss-Rhod PE and then endogenously stimulated to generate $ONOO^{-}$. **(A, B)** Cells were labeled with ER tracker. LNPs included a mixture of Liss-Rhod PE / DOPE / DOTMA at a 0.5 : 49.75 : 49.75 molar ratio. **(C, D)** LNPs included a mixture of Liss-Rhod PE / **DPPC-TC-ONOO⁻** / DOPE / DOTMA at a 0.5 : 4.5 : 47.5 : 47.5 molar ratio. Scale bars = 20 μm.

### Confocal Images of Live Cells with TEG-TC or TEG-TC-ONOO⁻

Both HeLa and RAW 264.7 cells were seeded into a 35 mm glass microscope dish and cultured overnight in DMEM with 10% FBS, 37 °C, 5% $CO_2$. A 20 μM solution of either **TEG-TC** or **TEG-TC-ONOO⁻** was directly added to live cells grown in glass-bottomed dishes and the cells were incubated at 37 °C for 20 minutes. For endogenous stimulation of peroxynitrite in HeLa cells, a solution of IFN-γ (100 ng/mL) and LPS (1 mg/mL) in Opti-MEM (250 μL) was added to the cells. The sample was incubated for 12 hours, then the media was decanted. Prior to imaging, a solution of PMA (10 nM) in HBSS (250 μL) was added and the resulting sample was incubated for 60 minutes at 37 °C. For endogenous generation of peroxynitrite through stimulation in RAW cells, LPS (100 ng/mL) in Opti-MEM (250 μL) was added to the cells and the resulting samples were incubated for 16 hours overnight.

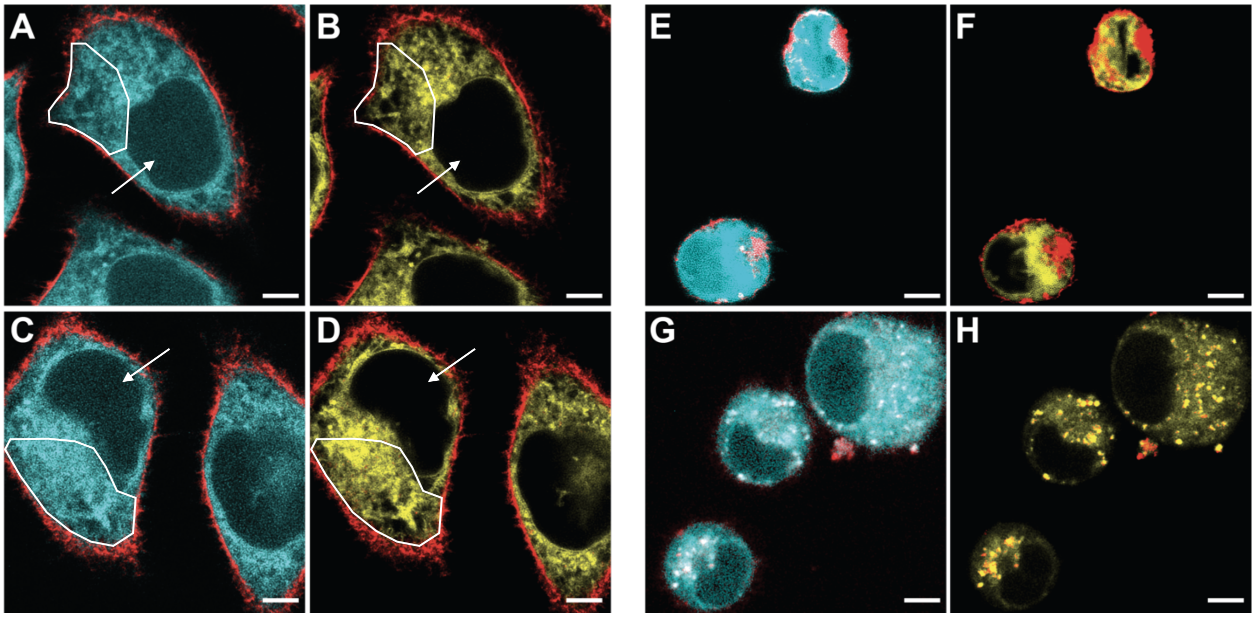


**Supplemental Figure 13.** Confocal images of HeLa cells treated with **(A, B) TEG-TC** only or **(C, D)** IFN-γ/LPS/PMA, followed by **TEG-TC-ONOO⁻**, and RAW 246.7 cells treated with **(E, F) TEG-TC** only or **(G, H)** LPS, followed by **TEG-TC-ONOO⁻**. Cell cytoskeleton was assessed with CellMask™ Deep Red Actin Tracking Stain. TEG-TC channel: 405/475 nm. White arrows and polygons **(A–D)** indicate representative locations that highlight signal intensity differences between the TEG-TC and ER channels. Scale bars = 5 μm.

### Cellular Viability in PCLS Using LDH Leakage and WST-1 Reduction

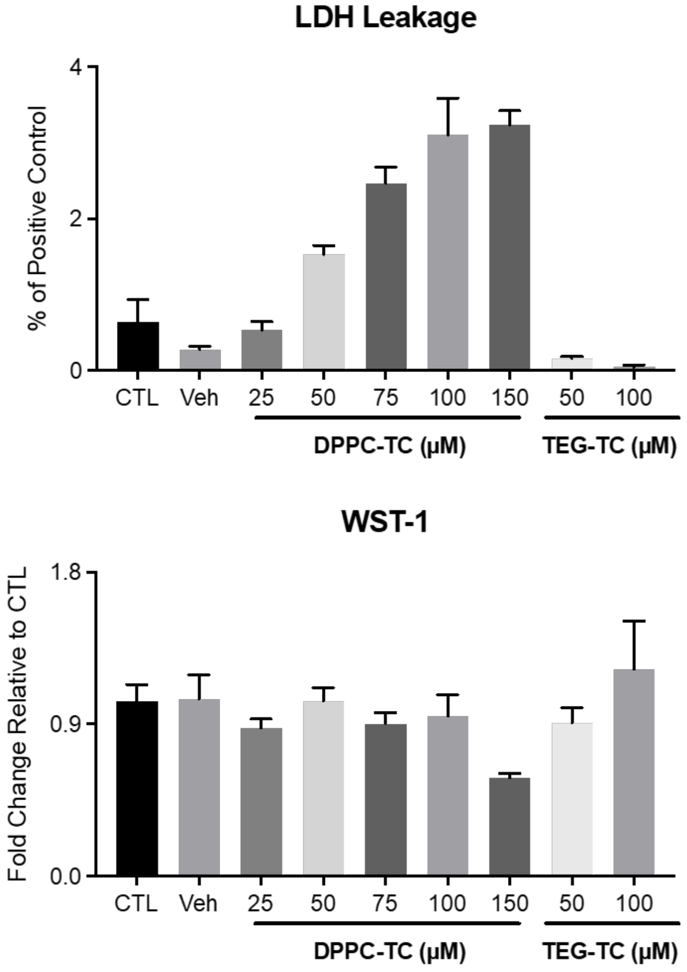


**Supplemental Figure 14.** Cytotoxicity of **DPPC-TC** in PCLS. Lung slices were incubated for 1 hour with varying doses of **DPPC-TC** or **TEG-TC** and assessed for viability by measuring LDH release to the medium or WST-1 reduction within the slice using established techniques (PMID 36692165). LDH activity was measured in the medium using the cytotoxicity detection assay (Roche) using a spectramax spectrophotometer. The quantity of released LDH is expressed as a percentage of LDH in the medium from lysed slices. WST-1 reduction was measured following incubation with Cell Proliferation agent, WST-1 (Roche) and is expressed as a ratio of reduction in untreated slices.

### Supporting Confocal Images of PCLS

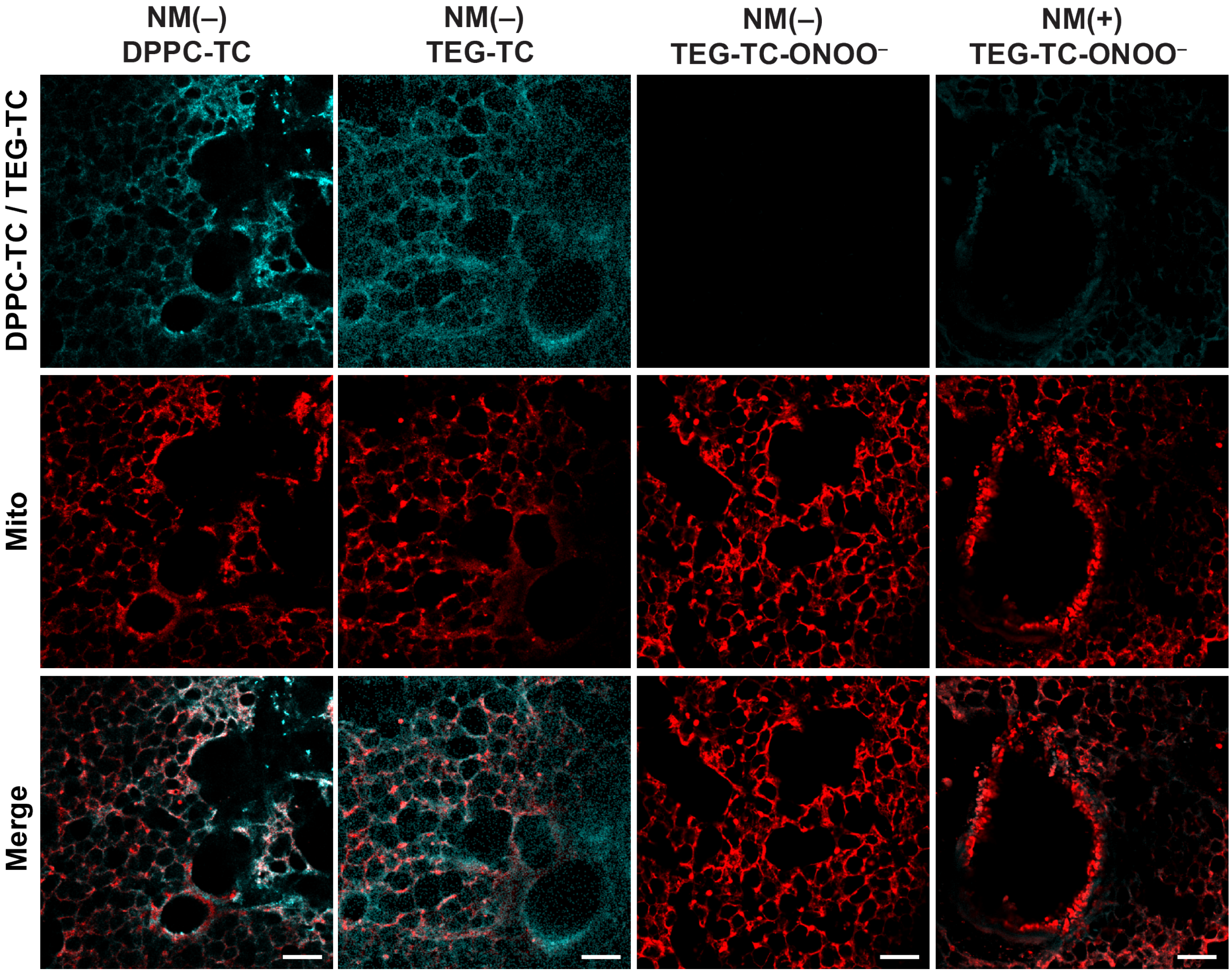


**Supplemental Figure 15.** Confocal images of PCLS incubated with **DPPC-TC** (amphiphile, positive control), **TEG-TC** (non-amphiphile, positive control), or **TEG-TC-ONOO$^-$** (non-amphiphilic probe). NM (+): nitrogen mustard exposure. Scale bar = 100 μm. The channel “DPPC-TC/TEG-TC” represents excitation at 405 nm.

### Viability Analysis of BAL Cells

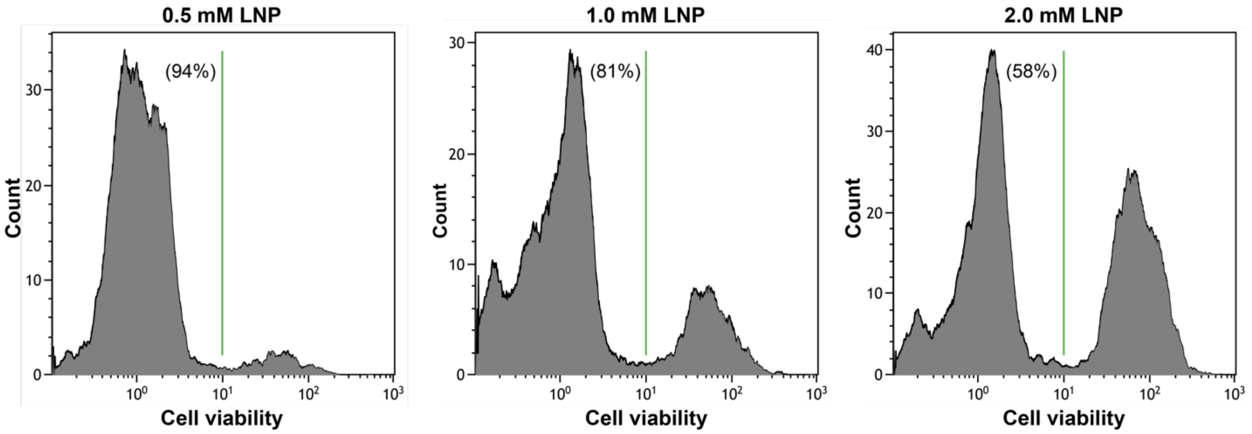


**Supplemental Figure 16.** Viability of BAL cells from mice instilled with various concentrations of LNPs.

### Gating Strategy for Flow Cytometry Data

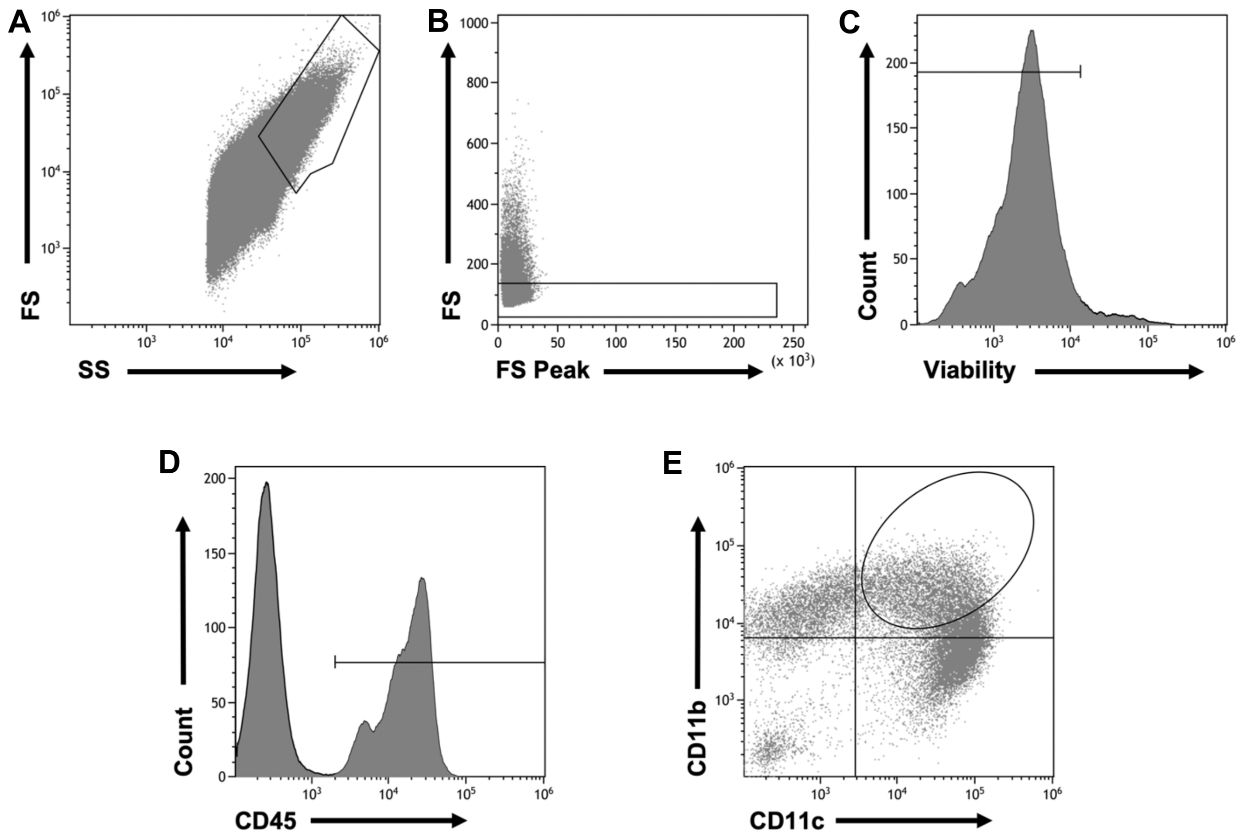


**Supplemental Figure 17.** Analyses of alveolar macrophages in BAL fluid using flow cytometry. BAL was performed 3 hours post the final LNP instillation. Cells were isolated and immunostained with fluorescent antibodies (CD45, CD11b and CD11c) followed by a viability dye (Fixable Viability Dye eFluor 780). The immunostained BAL samples were analyzed using a Gallios 10-color flow cytometer (Beckman Coulter, CA, USA). The flow cytometry data were analyzed using Kaluza software, and cells were gated based on **(A, B)** size and complexity, **(C)** viability, and **(D)** CD45 positivity. The myeloid lineage was determined based on the positive staining with CD45. **(E)** The CD45+ cells were further analyzed for CD11b and CD11c expression.

### CD11b versus CD11c Expression

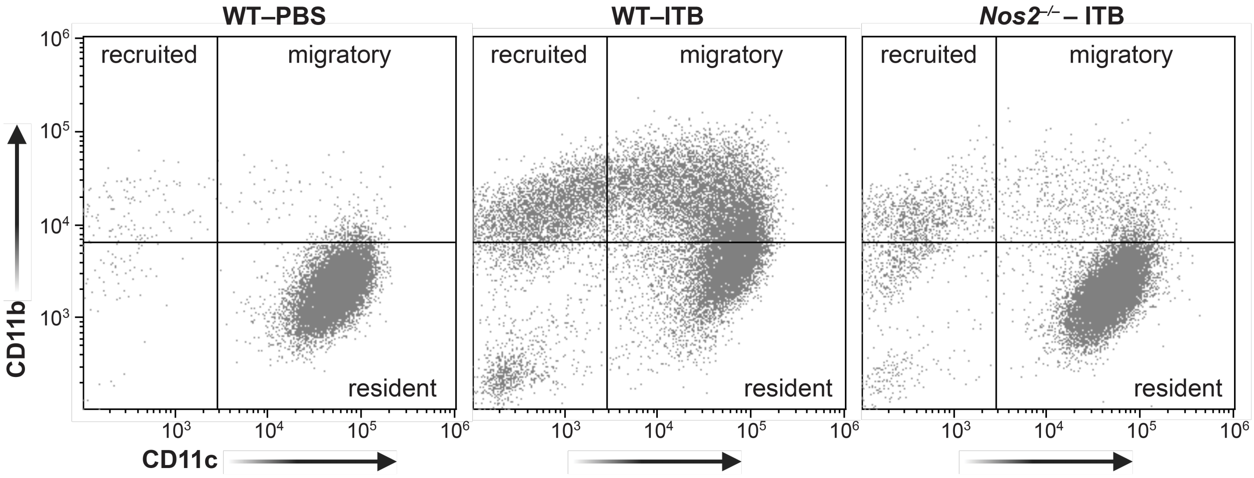


**Supplemental Figure 18.** Scatter-plot analyses of CD11b versus CD11c expression in BAL cells.
